# Proteinaceous Metal-Binding Eph-Ephrin Tetramerization is Modulated by Copper and Chelators

**DOI:** 10.64898/2025.12.08.693004

**Authors:** Amav Khambete, Hanghang Wang, Miah Henkemeyer, Mark Henkemeyer

## Abstract

Eph-Ephrin are powerful signaling molecules that form dimers, tetramers, and tetramer superclusters. The tetramerization mechanism — central to how these receptor-ligand molecules become activated — remains poorly understood. We find EphrinB2 is a proteinaceous chelator that binds to two juxtaposed arginine-rich electropositive pockets on the surface of EphB1 and EphB2 receptors when presented as two precisely aligned EphB-EphrinB2 dimers. This snaps the two low-affinity dimers into a high-affinity circular tetramer. We show EphrinB2 binds copper, consistent with ancestry to Cupredoxins, explaining why metal chelators such as EDTA, EGTA, and 8-hydroxyquinoline are potent tetramer inhibitors: they bind the EphB arginine-rich, copper-like pocket to compete with EphrinB2. Our findings reveal how small molecules, pH, salts, and copper dynamically modulate the kinetics of Eph–Ephrin binding and provide a mechanistic framework for therapeutic targeting of the tetramer.

**SIGNIFICANCE:** The Eph receptor tyrosine kinases interact with their membrane-anchored Ephrin ligands to form dimers, circular tetramers, and tetramer superclusters at cell-cell contact to become activated and transduce powerful bidirectional signals. We find that copper, pH, salts, and metal chelators modulate dimerization/tetramerization dynamics to affect downstream signaling. Ephrins are copper-binding proteins, consistent with their ancestry to other copper-binding proteins, while the Eph tetramer interface is highly electropositive, mimicking copper and targeted by metal chelators that disrupt tetramer assembly. These findings explain the molecular mechanisms that drive Eph-Ephrin dimer/tetramer assembly and regulate bidirectional signaling.

## INTRODUCTION

Eph receptors, the largest family of receptor tyrosine kinases, and their corresponding Ephrin ligands form a conserved group of signaling molecules vital for cell-cell communication (1, 2). Found throughout the metazoan lineage (3), Eph-Ephrin signaling regulates embryonic development and adult tissue maintenance (4). Unlike systems that use soluble ligands, Eph-Ephrin interactions are contact-dependent because both molecules are tethered to the plasma membrane, allowing direct communication between adjacent cells and transmission of bidirectional phosphotyrosine signals that are propagated into both the receptor-expressing cell (forward signaling) and the ligand-expressing cell (reverse signaling) (5).

Studies of Eph-Ephrin are confounded by the large number of Eph receptors (14) and Ephrin ligands (8), their promiscuous binding with A, B, and cross A/B subclasses (6), their ability to form *in-trans* interactions between contacting cells and *in-cis* interactions within the same cell (7), and their ability to crosstalk with many other receptor systems and signaling pathways (8). Eph-Ephrin bidirectional signaling has strong effects on cytoskeletal dynamics and can modulate the activities of small GTPases, Ras/MAPK, Src family tyrosine kinases, PI3K, JAK/STAT, Wnt, and SMAD pathways (8, 9). They further control integrins, cadherins, focal adhesions, and tight junctions, and interact with and regulate multiple cell surface receptors, most notably the N-methyl-D-aspartate (NMDA) receptor calcium channel in neurons as well as receptors for Epidermal Growth Factor, Fibroblast Growth Factor, Platelet-Derived Growth Factor, and Transforming Growth Factor (1, 2, 8–10). With the large number of possible Eph-Ephrin interactions, their ability to impact other cell signaling systems, and their strong influence on cell behaviors, Eph-Ephrin directed cell-cell signaling has emerged as a major player in physiology.

Select Eph and Ephrin molecules also become overexpressed and signal excessively in many chronic conditions. EphB1 and its ligand EphrinB2, for instance, become overexpressed/overactive after nerve injury to control the plasticity of neurons that signal chronic pain (11, 12), whereas EphB2 becomes overexpressed/overactive in most, if not all, chronic fibrotic disorders (13, 14). Genetic deletion of *EphB1* in mice strongly blunts models of chronic pain (11, 15, 16), while deletion of *EphB2* or point mutation that kills this receptor’s tyrosine kinase catalytic domain strongly mitigates animal models of inflammatory/fibrotic disorders (17–21). EphrinB2 reverse signaling is also important in chronic conditions, as shown in our accompanying manuscripts on chronic pain (22) and pancreatic cancer cell migration (23).

Eph and Ephrin ectodomains interact upon cell-cell contact to form dimers, circular tetramers, and tetramer superclusters to become activated and transduce bidirectional signals (24). Because both forward and reverse signaling contribute to pathology, we searched for ways to block the receptor-ligand interaction to blunt both signals. This work identified a group of orally available small molecules that act as reversible competitive inhibitors of the Eph-Ephrin circular tetramer (22). We report here that tetramer inhibitors are also metal chelators that bind an arginine-rich electropositive region on the tetramerization interface of the Eph receptor to prevent Ephrin binding. We further find that copper alters dimer/tetramer dynamics and, contrary to assumptions that Ephrins do not interact with metals (3, 24, 25), show that EphrinB2 binds copper. Our studies suggest a conserved proteinaceous copper-binding activity of Ephrins recognizes electropositive regions of the Eph ectodomain to form the high-affinity circular tetramer. Targeting this electrostatic interaction may counteract excessive Eph-Ephrin signaling linked to numerous conditions.

## RESULTS

### Eph-Ephrin dimer/tetramer dynamics are altered by metal chelators and copper

The first 3D crystal structures were those of the EphB2 N-terminal globular ligand-binding domain (PDB: 1NUK) (26) and its co-crystal with EphrinB2 (PDB: 1KGY) (24). In the co-crystal, EphB2 and EphrinB2 assembled into two EphB2-EphrinB2 heterodimers that come together through a distinct interface to form a circular tetramer (Figure 1A; see Video S1). Tetramers further packed into the crystal through additional receptor-receptor and ligand-ligand contacts (24), resembling signaling superclusters observed on contacting cells (27, 28). Co-crystals of the vascular EphB4 receptor bound to EphrinB2 showed only dimers (29), and our biophysical data indicate EphB4-EphrinB2 is more dimer-driven (22). The EphrinB2 ectodomain folds into an eight-stranded *β*-barrel with a Greek key topology similar to the Cupredoxin family of copper-binding proteins (24), though Ephrins are assumed not to bind metals. In Wang et al. (22), we report our discovery and characterization of small molecules that selectively target formation of the Eph-Ephrin circular tetramer. Tetramer inhibitors as exemplified by novel compounds A20-I and 3511-I, contain an 8-hydroxyquinoline (8-HQ) bicyclic ring scaffold (Figure 1B) that was first synthesized almost 150 years ago. In fact, halogenated and other modified 8-HQ-containing compounds have been approved for human use, one sold for decades as an over the counter (OTC) medicine, and are used as a food additive to enhance livestock growth and are under development for multiple indications, including Alzheimer’s disease (30). 8-HQs are characterized by a hydroxyl group in the quinoline ring at position 8 and a nitrogen at position 1. This positioning makes them potent metal binding compounds, that require two or more 8-HQ molecules to cage a metal ion, similar to yet different from the larger and more electron donating EDTA and EGTA metal chelators. EDTA and EGTA trap metal ions within a single molecule.

**Figure 1.**
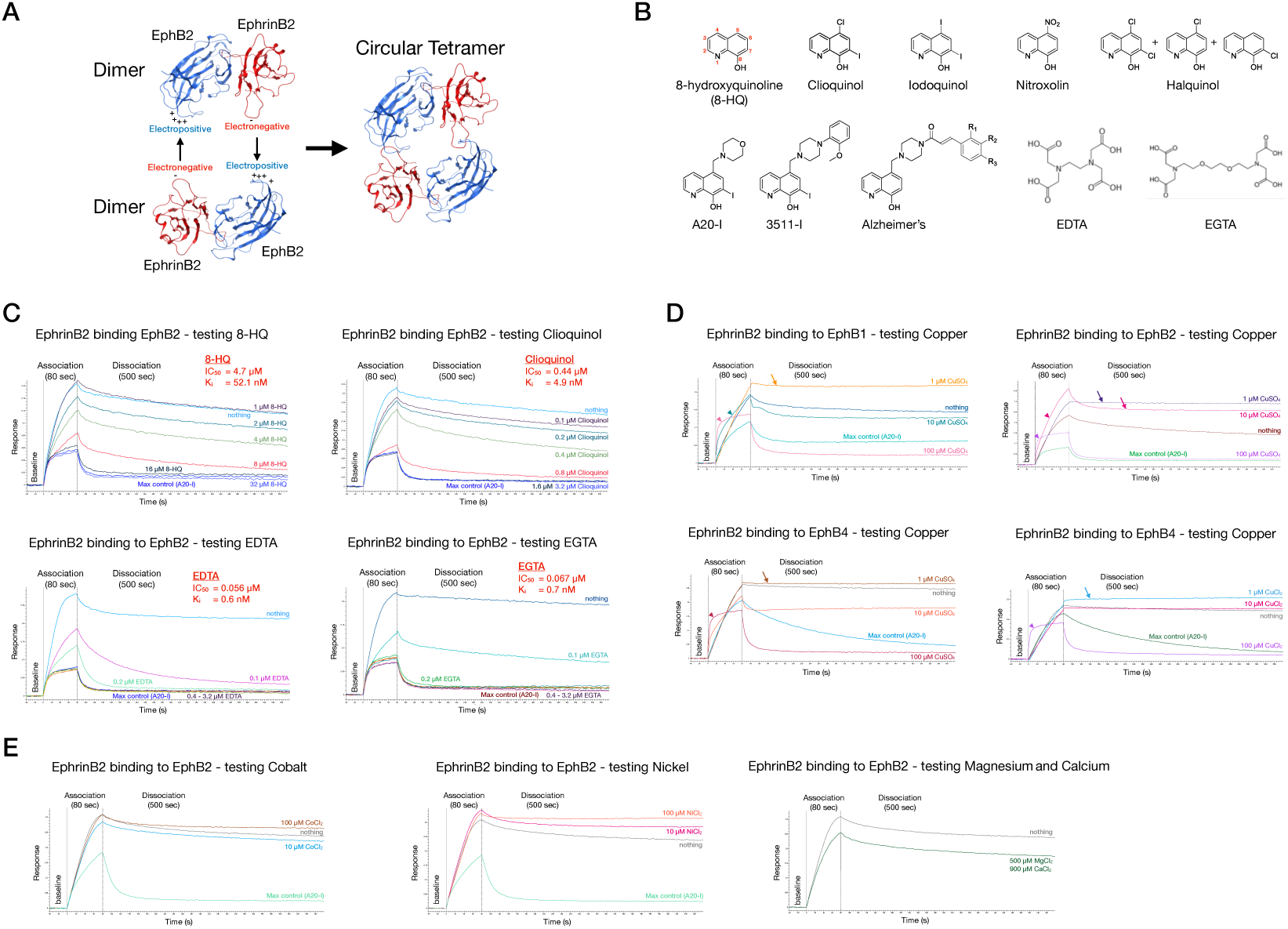
Metal chelators and copper are Eph-Ephrin tetramer inhibitors. (A) Ribbon schematic of the EphB2 (blue) and EphrinB2 (red) ectodomains showing two juxtaposed dimers forming the circular tetramer through tandem electrostatic interactions (PDB 1KGY). (B) The A20-based tetramer inhibitors A20-I and 3511-I contain 8-hydroxyquinoline (8-HQ), a bidentate metal chelator through its nitrogen and 8-hydroxyl oxygen. The multidentate chelators EDTA and EGTA have more donor atoms. (C) Biolayer interferometry (BLI) of the effects of 8-HQ, Clioquinol, EDTA, and EGTA on EphrinB2-EphB2 binding kinetics and tetramer formation. Immobilized human EphrinB2-Fc was exposed to 25 nM human EphB2-His with or without indicated concentrations of compound to determine tetramer-specific IC_50 a_nd K_i v_alues as described (22). Without compound, the sensorgram response over time showed complex 2:1 binding kinetics with formation of a a long-lasting tetramer, whereas sufficient inhibitor yielded mainly short-lived dimers. 8-HQ was a poor tetramer inhibitor, whereas EDTA and EGTA were superior. (D) BLI shows low copper enhances tetramer formation, while high copper favors dimer formation. EphB1-His, EphB2-His, and EphB4-His (25 nM) were tested with 1, 10, or 100 μM copper. Low copper maintained an elevated sensorgram response through dissociation (arrows), whereas high copper produced a steep rise and equally rapid loss (arrowheads). (E) High concentrations of cobalt, nickel, magnesium, and calcium had little effect on EphrinB2-EphB2 dimer/tetramer dynamics.

As the Eph-Ephrin tetramer inhibitors we identified were related to the above known 8-HQ chemicals, we first confirmed that they were metal chelators with copper binding activity (Figure S1A). We then used biolayer interferometry (BLI) and an Octet RED384 or R8e instrument (Sartorius) to assess in real-time the effect of various small molecule chelators on the complex dimer/tetramer binding kinetics of EphB1, EphB2, and EphB4 receptor ectodomains interacting with their common EphrinB2 ligand and as described (22). The BLI experiments were typically done by loading onto AHC or AHC2 biosensors a commercially available Ephrin ectodomain fused to Fc (e.g. human EphrinB2-Fc, R&D Systems) and then, after baseline measurements in buffer, exposing the Ephrin immobilized chips in timed association-dissociation steps to an Eph ectodomain binding partner (e.g. human EphB1-His, EphB2-His, or EphB4-His, Sino Biological) and as schematized (Supplementary Figure S1B). Here, testing the human EphB1-EphrinB2, EphB2-EphrinB2, and EphB4-EphrinB2 protein-protein interactions determined that A20-I, Clioquinol, EDTA, and EGTA at 3.2 μM were all potent tetramer inhibitors disrupting all three interactions, especially tetramer-driven EphB1-EphrinB2 and EphB2-EphrinB2 (Figure S1C). Our program Find50.py (22) then determined the tetramer-specific half maximal inhibition concentration (IC_50)_ and inhibition constant (Ki) of chelators. While 8-HQ itself was a relatively weak tetramer inhibitor with an IC_50 o_f 4.7 μM, Clioquinol with an IC_50 o_f 0.44 μM had the typical strong activity similar to our other 8-HQ-based compounds (22), while EDTA and EGTA were superior, with IC_50 v_alues below 0.1 μM and subnanomolar K_i v_alues (modified Cheng-Prusoff equation) (Figure 1C). We also assessed other non-8-HQ based chemicals for tetramer inhibitor and copper binding activity, identifying several common FDA approved drugs as metal chelators but not as tetramer inhibitors (Figure S1D). We did find approved metal chelators used for thalassemia (iron overload), Wilson’s disease (copper overload), and heavy metal poisoning, and a novel chelator molecule termed L08 identified in our high-throughput screen (HTS) (22) were all potent Eph-Ephrin tetramer inhibitors (Figure S1E and Table S1). The above data shows bidentate 8-HQ metal chelators like A20-I and Clioquinol have a strong effect on the tetramer, and that this ability is potentiated in multidentate chelators such as EDTA and EGTA.

The ability of small molecule chelators to disrupt the formation of Eph-Ephrin tetramers indicated possible role for metals in regulating the protein-protein interactions. To directly test this, we used BLI to assess dimer/tetramer dynamics in the presence of copper. This revealed a curious biphasic effect with low concentrations of copper leading to stabilization of the tetramer and high concentrations leading to accelerated dimer formation with complete loss of the tetramer species (Figure 1D).

The biphasic effect of copper was observed for all three human EphB1-EphrinB2, EphB2-EphrinB2, and EphB4-EphrinB2 interactions. BLI studies of EphB1 and EphB2 interacting with the related EphrinB1 ligand showed high copper also inhibited tetramer formation (Figure S1F), though low copper did not increase it, suggesting EphrinB2 and EphrinB1 respond differently to copper. Metals cobalt, nickel, magnesium, and calcium at high concentrations exhibited little effect on Eph-Ephrin binding (Figure 1E), while zinc and iron did (Figure S1G). Like what we showed for 8-HQ-based chemicals and EDTA (22), copper can also affect the ability of soluble EphrinB2-Fc ectodomain to bind and stimulate EphB2 catalytic forward signaling in Cos1 cells that endogenously express the receptor (Figure S1H). Like the BLI studies, we found copper exhibited a biphasic effect on EphB2 tyrosine kinase activity, with low concentrations enhancing phosphotyrosine signaling and high concentrations inhibiting it. Altogether, the ability of both metal chelators and copper to affect Eph-Ephrin dimer/tetramer dynamics suggests a novel molecular mechanism regulates the interactions of these molecules and modulates dimer and tetramer formation. This, in turn, will affect the ability of these receptors and ligands to form tetramer superclusters, become activated, and transduce forward and reverse signals into the cells that express them.

### Eph receptors bind chelators

The amino acid sequence of the EphB2 receptor tetramer interface is electropositive as it contains a cluster of four arginines — three from the F’-G loop (Arg87, Arg88, Arg89) and one from the K-L loop (Arg179) — which form a salt bridge with the EphrinB2 C-D loop aspartic acid (Asp69) (24) (Figures 2A-D; see Videos S2, S3, S4, S5, and S6). In addition to the strong Arg87_R-_Asp69_L e_lectrostatic bond and hydrogen bonds through Arg89_R-_Lys71_L,_ the EphB2 and EphrinB2 tetramer interfaces also associate through polar and van der Waals contacts. Notable are the stacking of the Phe128_R-_Tyr37_L a_romatic rings and the Phe135_R-_Glu34_L i_nteraction (see Video S7). As chelator molecules bind to positively-charged metal ions, we hypothesized they mimic the EphrinB2 C-D loop and interact with the electropositive arginine-rich tetramer interface of the EphB2 protein, which we propose looks like copper, displacing the electrostatic bond that normally drives tetramer formation. To test this, we synthesized a novel 8-HQ compound that contains a primary amine to enable immobilization on AR2G biosensors for BLI studies. The primary amine was first tested for tetramer inhibitor activity and found to have a relatively weak IC_50 o_f 3.9 μM for the EphB1-EphrinB2 interaction and 2.1 μM for the EphB2-EphrinB2 interaction (Figure 2E), similar to that observed for the 8-HQ base chemical (Figure 1C). We then immobilized the primary amine on AR2G biosensors and tested its interaction with the human EphB1, EphB2, EphB4, and EphrinB2 ectodomains, finding that EphB2 and EphB1 exhibited obvious 1:1 binding with fast-on/fast-off association/dissociation kinetics (Figure 2F).

**Figure 2.**
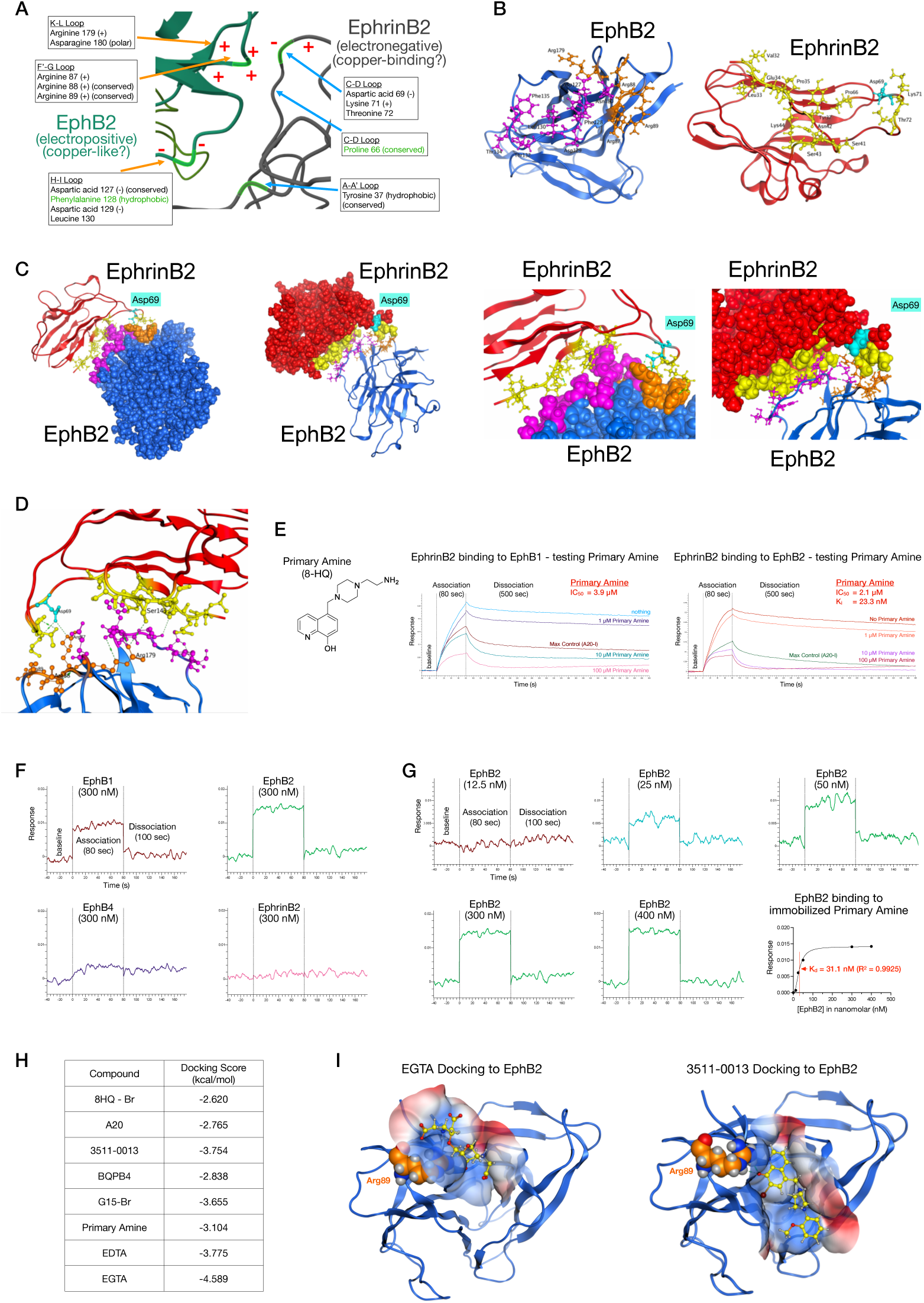
EphB2 tetramerization interface is electropositive and binds chelators. (A) Schematic of the EphB2-EphrinB2 tetramer interface, with the electropositive EphB2 loops F’-G and K-L interacting with the electronegative EphrinB2 loop C-D. Green amino acids show large conformational changes upon tetramerization. (B) Structures of monomeric EphB2 (left/blue; 1NUK) and EphrinB2 (right/red; 1IKO) ectodomains with key tetramer interface residues highlighted (EphB2 arginines, orange; EphrinB2 C-D loop aspartic acid, light blue). (C) Structures of EphB2-EphrinB2 co-crystals (1KGY) highlighting the tetramer interface amino acids, shown as full ectodomains and zoomed in on the tetramer interface. (D) Electrostatic map of the EphB2-EphrinB2 tetramer interface pinpoints the interaction between Arg87 of EphB2 and Asp69 of EphrinB2. (E) BLI of a novel 8-HQ compound with a primary amine on the EphrinB2–EphB1 and EphrinB2–EphB2 interactions. The tetramer IC₅₀ was 3.9 μM for EphB1 and 2.1 μM for EphB2, with an apparent Kᵢ of 23.3 nM. (F) AR2G biosensors loaded with the primary amine 8-HQ compound were exposed to 300 nM human EphB1-His, EphB2-His, EphB4-His, and EphrinB2-His. EphB2 and EphB1 showed 1:1 binding, while EphB4 and EphrinB2 did not. (G) BLI of ARG2 immobilized primary amine 8-HQ compound binding to 12.5–400 nM human EphB2-His showed saturable 1:1 binding with a Kᵢ of 31.1 nM (R² = 0.9925), similar to the Kᵢ calculated from IC₅₀ data in (E). (H) MOE docking of EGTA, EDTA, and six 8-HQ compounds to the EphB2 tetramer interface. All dock via ionic and hydrogen bond interactions with the conserved Arg89. (I) MOE docking of EGTA and 3511-0013 into the EphB2 tetramer interface (blue electropositive, red electronegative). Oxygens in the carboxylates of EGTA and in 3511-0013 form ionic and hydrogen bonds with Arg89.

Immobilized primary amine was then tested for binding to increasing amounts of EphB2, giving a dissociation constant (K_d)_ of 31.1 nM (Figure 2G). This indicates high-affinity binding of the primary amine to the EphB2 ectodomain, consistent with the estimated Kᵢ of 23.3 nM independently calculated from the above IC₅₀ data in Figure 2E.

EphB2 has four arginines in the tetramer interface, EphB1 has three arginines, and EphB4 only has two arginines. Because EphB4 does not make strong tetramers with EphrinB2 (Figure S1C) or appreciably bind the primary amine (Figure 2F), the higher electropositivity of the EphB2 and EphB1 tetramer interfaces likely drives tetramerization and the ability of the metal chelators to disrupt the tetramer. All 14 Eph receptors are electropositive in their respective F’-G and K-L loops with numerous arginine and lysine residues, especially Arg88 in EphB2, which is conserved in 12 of the 14 receptors (Figures S2A and S2B). Likewise, the tetramer interface of the 8 Ephrin ligands all share a conserved proline residue in their C-D loop and most have an electronegative aspartic or glutamic acid +3 from the proline (Figure S2C). This suggests a universal mechanism of electrostatic interactions may regulate dimer/ tetramer dynamics of all Eph-Ephrin family receptors and ligands. To explore this possibility, available Eph and Ephrin ectodomain crystal structures (Figure S2D) were studied using Molecular Operating Environment (MOE) for 3D visualization, modeling, and simulations. We first superimposed the backbones of Eph crystal structures to evaluate their overall 3D similarity, finding they are all similar throughout the N-terminal globular domain involved in Ephrin binding (Figure S2E). Within this globular ligand-binding domain, the 3D structures of the Eph tetramerization interfaces are more similar to each other than are their corresponding dimerization interfaces, which showed more variation in structure.

MOE in silico mutagenesis assessed the importance of Arg87, Arg89, Arg179, and Asn180 in EphB2 (Figure S2F). A proline at position 87 was predicted to disrupt tetramer affinity; EphB4, which has Pro87, makes weak tetramers with EphrinB2.

Mutating Arg87 to Asn87, Trp87, or Tyr87 maintained strong tetramer affinity, as did mutating Arg179 to Lys179 or mutating Asn180 to Lys180 or Arg180. Electropositivity of the F’-G and K-L loops is thus important for stable tetramers, and Pro87 as in EphB4 may disrupt them. Changing EphrinB2 Asp69 to Glu69, as in EphrinB1, would retain high affinity tetramers (Figure S2G).

We then used MOE docking to predict whether tetramer inhibitors interact with the EphB2 ligand-binding domain, examining 8-HQ based chemicals as well as EDTA and EGTA. Mirroring their tetramer inhibitor activities, 8-HQ gave a relatively weak docking score, followed by A20, and then stronger docking scores for the improved tetramer inhibitors 3511-0013, BQPB4, and G15, with best scores computed for EDTA and EGTA (Figure 2H). All compounds docked to the same electropositive region of the EphB2 tetramer interface and were predicted to form ionic and hydrogen bonds with Arg89, which is conserved in all EphB receptors (Figures 2I, S2H, and S2I; see Videos S8, S9, and S10). Schrödinger Glide and Maestro confirmed these results, with A20 also forming an ionic interaction with Arg89 (see Video S11). Nanoscale Molecular Dynamics (NAMD) 30 nanosecond (ns) simulations of EGTA and 3511-0013 binding the EphB2 ectodomain indicated EGTA has a >2-fold stronger total PBSA binding energy (−0.687 ± 0.036 kcal/mol) compared to that for 3511-0013 (−0.290 ± 0.017 kcal/mol) (Figure S2J). Further, while EGTA remained stably docked to EphB2 throughout the 30 ns simulation, 3511-0013 was less stable and after ∼22 ns completely dissociated from the receptor protein (Figure S2K; see Videos S12 and S13). This is likely because the multidentate EGTA can interact with both Arg89 and Arg87, while 3511-0013 only targets Arg89 (Figure S2L). Tetramer inhibitors thus bind with significant affinity through ionic bonds with the electropositive Eph tetramer interface.

### EphrinB2 binds copper

The BLI studies shown above in Figure 1 indicate copper can influence Eph-Ephrin dimer/tetramer dynamics. This prompted us to revisit the notion that Ephrin proteins do not bind copper (3, 24, 25). We first superimposed the crystal structures of four Cupredoxin proteins with crystal structures of six different Eph receptors and six different Ephrin ligands. Although Cupredoxins have no relation to Eph proteins, they do exhibit high structural similarity to Ephrins, including an overall shared Greek key *β*-barrel topology (Figure 3A), though Ephrins lack the canonical histidine residues typically found in the Cu(II) binding sites of Cupredoxins. Instead, Ephrins as a group share structural features important for dimerization and tetramerization with their cognate Eph receptors (Figure S3A). We used MOE to dock copper with the EphrinB2 structure. This identified a potential interaction of the metal with the C-D loop of the EphrinB2 tetramerization interface, particularly involving an ionic interaction between Cu²⁺ and Asp69, as well as coordinate covalent bonds between Cu²⁺ and both Lys71 and Thr72 (Figures 3B and S3B; see Video S14). NAMD indicated a sustained interaction of copper with Asp69/Lys71/Thr72 throughout a 30 ns simulation (Figure S3C-S3E, see Video S15), more stable and energetically favored (−0.794 0.0172 kcal/ mol) than EGTA or 3511-0013 binding to EphB2.

**Figure 3.**
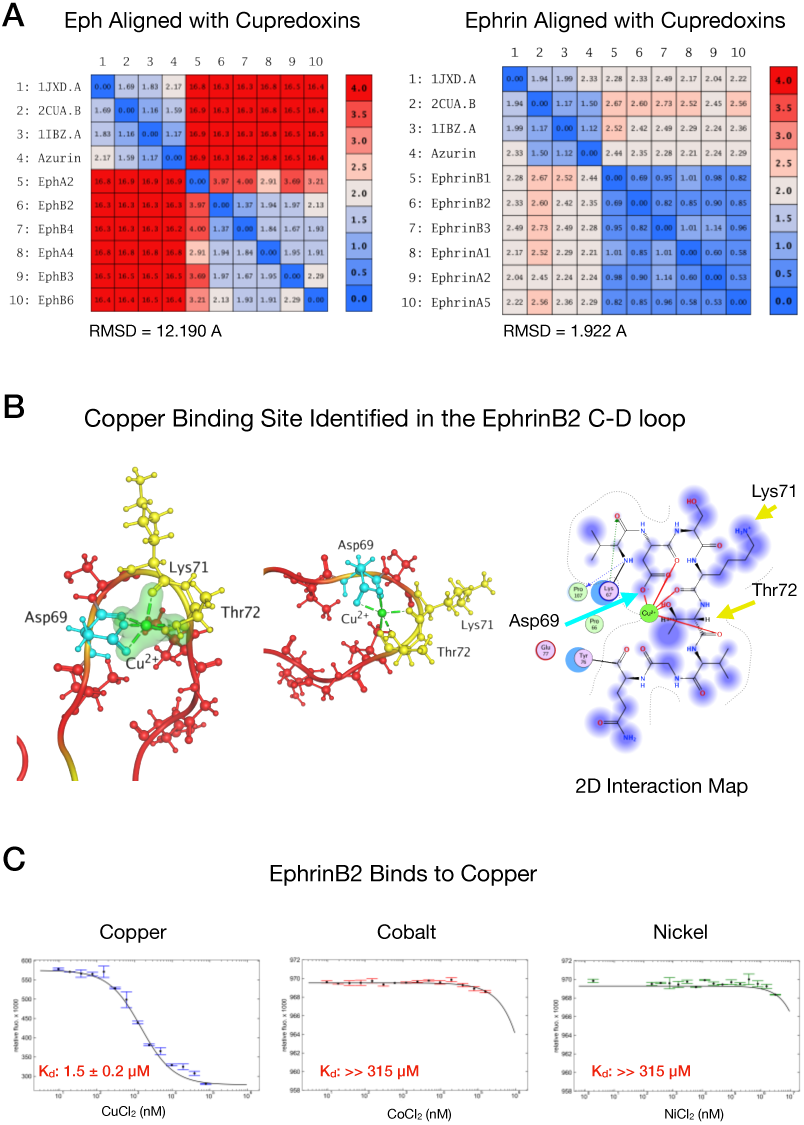
EphrinB2 tetramerization interface is electronegative and binds copper. (A) MOE backbone superimposition of crystal structures comparing Cupredoxins with Eph (left) and Ephrin (right) ectodomains. Ephrin ectodomains exhibit high structural similarity to Cupredoxins, whereas Eph ectodomains do not. (B) MOE SiteFinder identified a copper-binding site in the EphrinB2 C–D loop, where Cu²⁺ is coordinated by oxygen atoms and stabilized by Asp69, Lys71, and Thr72; NAMD confirmed copper remained bound over 30 ns. (C) MST of Alexa Fluor 488 labeled human EphrinB2 with CuCl_2,_ CoCl_2,_ or NiCl_2._ Copper quenched fluorescence in a concentration-dependent manner with a K_d o_f 1.5 μM. Cobalt and Nickel had little or no effect on thermophoresis.

To experimentally test if EphrinB2 can bind copper, the human ectodomain protein was sparsely labeled on lysine residues with Alexa 488 NHS dye and then incubated with increasing concentrations of copper, cobalt, or nickel (Figures 3C, S3F-S3K). This revealed copper was able to quench, in a concentration-dependent fashion beginning at 20-40 nM the cold fluorescence emitted from dye-labeled EphrinB2 with a K_d o_f 1.5 μM, aligning with the above BLI and Cos1 cell studies that showed copper affects Eph-Ephrin interactions and cell signaling at 1-100 μM. Copper’s strong quenching of the dye’s fluorescent signal indicates the EphrinB2 protein is binding to the metal. SD tests using detergent and heat to denature the protein returned the signal, confirming that the quenched fluorescence was due to specific EphrinB2-copper interactions and not protein precipitation or aggregation. Cobalt or nickel showed no effect on cold fluorescence when added to the dye-labeled EphrinB2 protein. Further, while copper quenching of dye-labeled EphrinB2 precluded microscale thermophoresis (MST) studies of copper, MST studies of cobalt and nickel confirmed these metals had little if any effect on EphrinB2 protein movement, except perhaps for the highest concentration tested (312.5 μM). Confirming this, additional BLI studies with up to 500 μM of these two metals verified that neither were able to block the ability of EphB2 to form tetramers with EphrinB2 or EphrinB1 (Figures S3L and S3M).

To probe why copper had a strong effect on fluorescence dye-labeled EphrinB2, MOE simulations indicate the most exposed lysine on the protein’s surface is the lysine residue at position 71 (Figure S3N). This indicates Lys71 is likely a predominant dye attachment residue. Lys71 is also part of the EphrinB2 tetramer interface C-D loop and adjacent to Asp69, within 10 angstroms from the copper binding site, and predicted to form one of the coordinate covalent bonds with copper. This would be consistent with copper binding to EphrinB2 likely producing a significant conformational change in this loop to alter fluorescence from the nearby attached dye. Altogether, our data indicate Ephrins stably bind copper within the C-D loop, and likely explains why this metal has such a strong effect on Eph-Ephrin dimer/tetramer dynamics.

### Tetramerization is extremely stable and driven by electrostatic interactions

NAMD simulations of EphB2-EphrinB2 dimer and tetramer binding interactions indicated the tetramer is much more stable than the dimer, with lower total and electrostatic free energy ΔG (Figures 4A, 4B, 4C, S4A, and S4B; see Videos S16 and S17). Van’t Hoff analysis indicated tetramer formation is >750-fold more stable than the dimer (Figure S4C). BLI experimentally confirmed that the tetramer is essentially irreversible while dimerization rate constants changed with temperature (Figures 4D and S4D). Using a conservative K_D1 v_alue for the EphB2-EphrinB2 tetramer of 0.28 nM (22), free energy calculations based on the BLI data confirmed the tetramer is at least 250-fold more stable than the dimer (Figure S4E).

**Figure 4.**
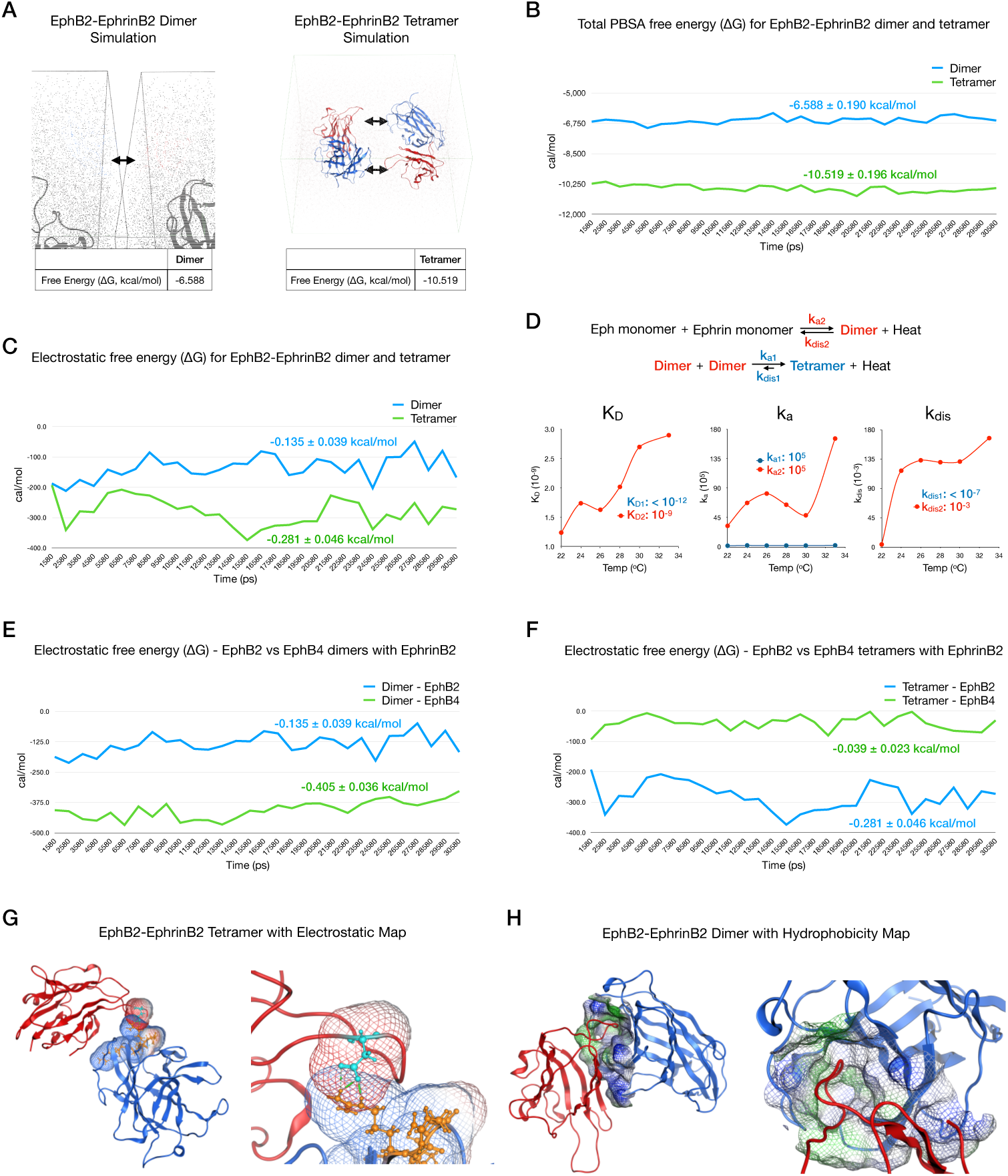
The EphB2-EphrinB2 tetramer is more stable than the dimer and is driven by electrostatic interactions. (A) EphB2 and EphrinB2 structures were prepared in MOE; the dimer (one of each) and tetramer (two juxtaposed dimers) were simulated for 30 ns with CHARMM36 in NAMD. (B) MM/PBSA free energy (ΔG) calculations every 50 ps indicate stable high affinity interactions for both the dimer and tetramer. The difference (ΔΔG = −3.931 kcal/mol) corresponds to a >725-fold increase in tetramer stability. (C) Electrostatic PBSA free energy (ΔG) calculated every 50 ps shows the tetramer interaction is more electrostatically driven than the dimer. (D) BLI indicates the tetramer is stable across temperatures, whereas the dimer is more susceptible to temperature changes. Human EphrinB2-Fc binding to 3.125-100 nM human EphB2-His at 22-33 oC was fit to 2:1 kinetics; the tetramer was effectively irreversible at all temperatures. (E) Electrostatic PBSA free energy (ΔG) shows the EphB4-EphrinB2 dimer interaction is 3x more electrostatically driven compared to the EphB2-EphrinB2 dimer. (F) Electrostatic PBSA free energy (ΔG) of tetramer binding, with EphB4 superimposed onto EphB2, shows the EphB2-EphrinB2 tetramer is >7x more electrostatically driven than the EphB4-EphrinB2 tetramer. (G) Electrostatic map of one EphB2-EphrinB2 tetramer interface focused on the EphB2 arginine-rich pocket and EphrinB2 C-D loop. (H) Hydrophobic grid shell of the EphB2 dimer interface, which mediates hydrophobic interactions with EphrinB2 loops F-G and G-H that are weaker and more temperature dependent than tetramer interactions.

As exemplified in Figure S1C and in more detail in our accompanying manuscript (22), the interaction of EphrinB2 with EphB2 is tetramer-driven and strongly affected by metal chelators, whereas its interaction with EphB4 is more dimer driven and not nearly as affected by chelators. To explore this difference, we conducted additional NAMD simulations to compare EphB2-EphrinB2 and EphB4-EphrinB2 dimer and tetramer dynamics, revealing that in both cases the tetramer exhibited a much higher affinity interaction versus the respective dimer (Figure S4F). Focusing on electrostatic interactions, the EphB4-EphrinB2 dimer has 3x more electrostatic binding energy compared to the EphB2-EphrinB2 dimer (Figure 4E), whereas the EphB2-EphrinB2 tetramer has >7x more electrostatic binding energy than the EphB4-EphrinB2 tetramer (Figure 4F). These data are consistent with experimental evidence that EphB4-EphrinB2 is more dimer-driven and EphB2-EphrinB2 more tetramer-driven.

Because the EphrinB2 C-D loop is negatively charged (Asp69) and the EphB2 tetramer interface has an electropositive pocket (Arg87, Arg88, Arg89, and Arg179), MOE docking and NAMD predict strong ionic salt bridges at the tetramer interface (Figures 4G, S4G; see Video S18). The EphB2 dimerization channel, which receives the inserted EphrinB2 G-H loop, is largely hydrophobic (Figure 4H; see Video S19). Upon protein-protein contact, the EphB2 dimer channel changes conformation dramatically while the tetramer loops change minimally (Figure S4H), as do EphB2 dimer side chains (Figures S4I and S4J), showing the hydrophobic dimerization interface is flexible while the tetramer interface is rigid. However, residues in the EphrinB2 C-D loop that form ionic and hydrogen bonds with EphB2 exhibit significant conformational changes upon binding to the receptor, and residues at both ends of this loop, Pro66 and Tyr76, show major hinge-like movements upon tetramerization (see Video S20). In EphB2, the electropositive arginines moved somewhat and Phe128 moved substantially to form the van der Waals interaction with Tyr37 of the ligand (see Video S21).

MOE SiteFinder identified potential binding pockets on the EphB2 monomer, EphrinB2 monomer, and EphB2-EphrinB2 dimer (Figure S4K). In the EphB2 monomer, SiteFinder identified residues in the dimerization interface, whereas in the EphrinB2 monomer both tetramerization and dimerization interfaces were identified, and in the EphB2-EphrinB2 dimer only the tetramerization interface of EphrinB2 was identified. These data are consistent with an initial Eph-Ephrin interaction favoring dimer formation that then leads to tetramerization.

### Dimer/tetramer dynamics are influenced by pH and salt

Aspartic acid at neutral pH will have a negative charge, whereas in an acidic buffer condition it should be neutral. As we hypothesize EphB2-EphrinB2 tetramerization relies on negatively charged Asp69 in the EphrinB2 C-D loop interacting with positively charged arginines on the EphB2 protein, a low pH may alter dimer-tetramer dynamics. We therefore conducted BLI studies to compare the kinetics of immobilized human EphrinB2 binding to soluble human EphB1 and EphB2 receptors in an acidic environment (Figure 5A). At normal pH of 7.4, the EphB1-EphrinB2 and EphB2-EphrinB2 interactions both exhibited the typical complex association/dissociation kinetics of dimer and tetramer formation, with highly stable tetramers persisting throughout the dissociation. However, at pH 3.6, the association kinetics were completely disrupted, there was a rapid increase in dimer kinetics within seconds of association onset, then an equally rapid loss of signal once the biosensor was placed in dissociation. This indicates stable tetramers do not form in low pH, only weak dimers with a fast-off profile. The kinetics in acidic conditions were similar to BLI runs using high concentrations of copper and show a shift to a more simple 1:1 dimer interaction (Figure 1D). Similar pH-dependent binding dynamics were obtained using immobilized EphB1, EphB2, EphB3, and EphB4 ectodomains binding to soluble EphrinB2 (Figure 5B). The 1:1 kinetics at low pH were reversed by neutralizing the buffer, indicating protein structures were not irreversibly damaged by the acidic environment (Figure 5C). At low pH, dimer binding is thus enhanced while tetramer formation is eliminated, consistent with Asp69 electrostatic interactions being important for tetramer stability and unable to form when aspartic acid is uncharged.

**Figure 5.**
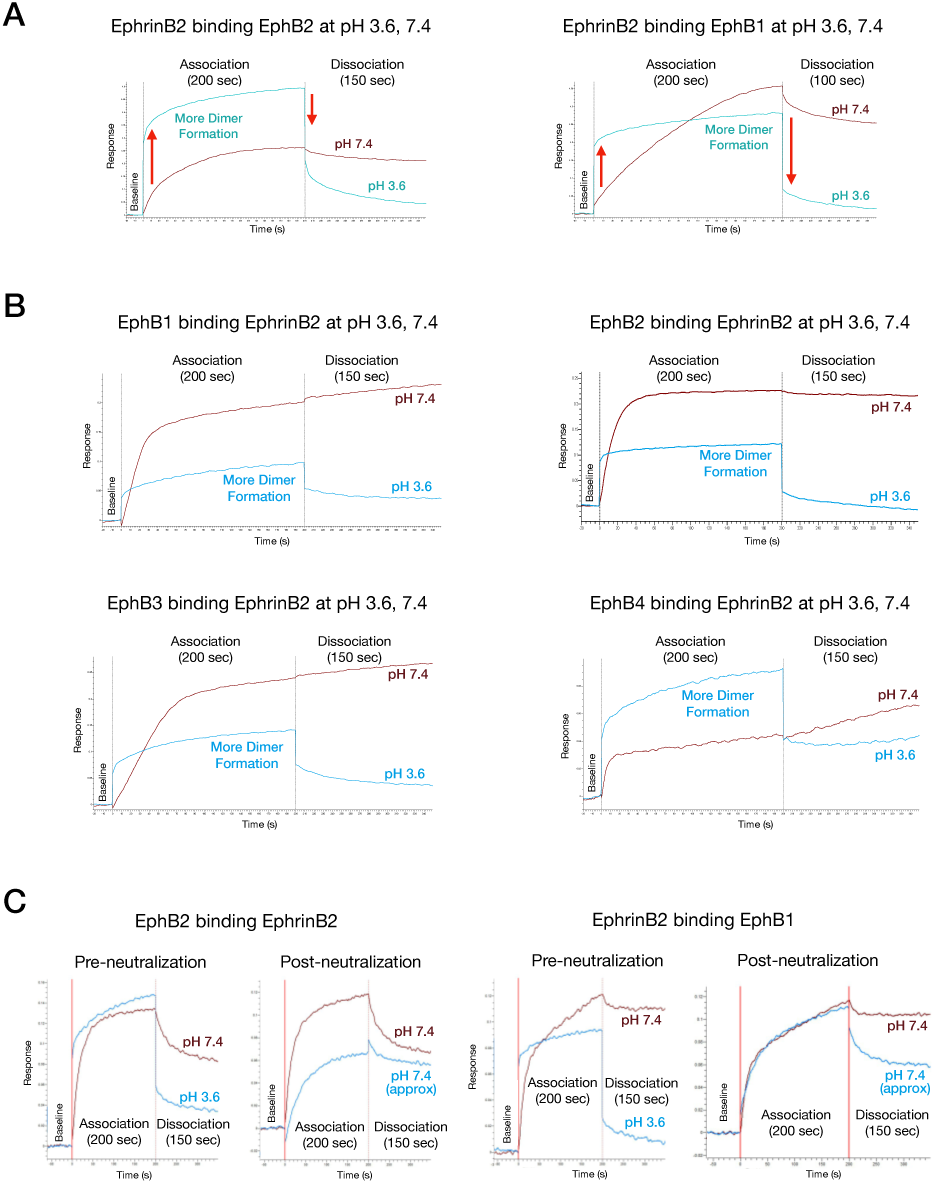
EphB2-EphrinB2 tetramerization is disrupted by low pH. (A) BLI of immobilized human EphrinB2-Fc binding to 50 nM soluble human EphB2-His and EphB1-His at pH 7.4 (brown) or pH 3.6 (blue). At acidic pH, tetramer formation is greatly curtailed while dimer binding is rapid and flatlines. (B) BLI of rat EphB1-Fc, mouse EphB2-Fc, human EphB3-Fc, or human EphB4-Fc binding to 100 nM mouse EphrinB2-His at pH 3.6 (blue) or 7.4 (red). Acidic pH essentially abolishes tetramers. (C) BLI of mouse EphB2-Fc binding to 100 nM mouse EphrinB2-His (left) or human EphrinB2-Fc binding to 50 nM human EphB1-His (right) at pH 7.4 (brown) or 3.6 (blue). Tetramer activity lost at pH 3.6 was restored by neutralization.

We also probed electrostatic interactions with NaCl to mask charge-charge interactions. NaCl reduced in a concentration-dependent manner EphB2-EphrinB2 tetramerization, while dimerization was unaffected (Figure 6A). This indicates salt bridges are needed for tetramerization but not dimerization. In a similar experiment, individual amino acids were added to BLI experiments testing EphrinB1 and EphrinB2 binding to EphB2. Neutral amino acids (at pH 7.4) like proline or alanine were predicted to have no effect on EphB-EphrinB dimer/tetramer kinetics, while we thought charged amino acids like arginine or glutamic acid might mask the electrostatic interactions and affect tetramer binding. Indeed, proline and alanine had no effect on the EphB-EphrinB interactions, while Arg and Glu both decreased tetramer binding in a concentration-dependent manner (Figures 6B and S5A). Dimer interactions were not inhibited, indicating salt bridges are not needed for dimer formation. At very high concentrations of arginine, dimerization was enhanced (Figure 6C), similar to a high copper concentration or acidic buffer condition. High arginine may disengage Asp69 and the C-D loop from the Eph tetramer interface, as copper or protons may disrupt the C-D loop and the tandem electrostatic interactions that cement two dimers into a tetramer.

**Figure 6.**
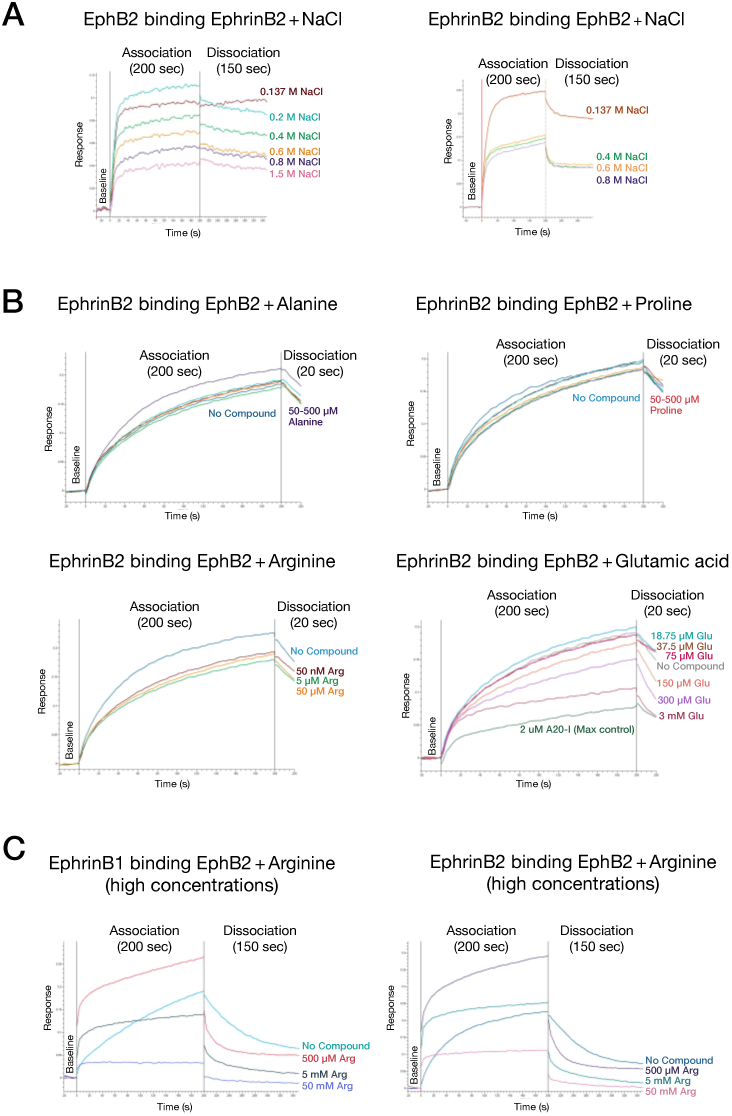
EphB2-EphrinB2 tetramerization is disrupted by high salt and charged amino acids. (A) BLI of immobilized EphB2-Fc and EphrinB2-Fc binding to soluble EphrinB2-His and EphB2-His shows increasing salt reduces tetramer formation, indicating salt bridges are important for tetramerization but not dimerization. (B) BLI of human EphrinB2-Fc binding to 50 nM human EphB2-His with excess L-alanine, L-proline, L-arginine, or L-glutamate. Alanine and proline had little effect, whereas arginine and glutamic acid reduced tetramer formation without affecting dimer kinetics. (C) BLI of human EphrinB1-Fc or EphrinB2-Fc binding to 50 nM human EphB2-His shows high L-arginine reduced tetramer formation while enhancing the dimer, suggesting selective disruption of tetramer interfaces.

### Arg87 in the F’-G loop of EphB2 is involved in Ephrin binding

In the above experiments and in (22) we used full ectodomain proteins of Eph receptors, which are ∼550 amino acids in length, to study Ephrin binding. As the protein used to determine the crystal structure of the ligand-binding region of EphB2 was much smaller and restricted to the first 210 amino acids of the mouse protein (24, 26), we studied a similarly sized fragment to determine if the dimer-tetramer dynamics documented with the full ectodomains are recapitulated using the smaller fragment. To do this, the N-terminal sequence of human EphB2 was expressed in mammalian cells and the EphB2-WT (19–210) protein purified. A mutant version, EphB2-R79P, was also expressed and purified that contains an arginine-to-proline substitution in residue 79 of the human protein (Figure S6A). As human Arg79 corresponds to Arg87 in mouse EphB2 loop F’-G discussed above to form the salt bridge with Asp69 of EphrinB2, we anticipated changing this residue to a proline, as in EphB4, would disrupt dimer-tetramer dynamics. Following expression and purification of the custom proteins (by Sino Biological), their sequence was confirmed at the UT Southwestern Proteomics Core facility (Figure S6B), and silver stained gels and anti-EphB2 immunoblots were used to determine concentrations of the proteins so that equal amounts were used in BLI experiments (Figures S6C and S6D).

EphB2-WT and EphB2-R79P were tested in Octet at 3, 9, 27, and 81 nM concentrations for binding to human EphrinB3 (Figures 7A and S6E), EphrinB1 (Figure S6F), and EphrinB2 (Figure S6G) for 800 sec association and 2,400 sec dissociation steps. Qualitatively, both proteins exhibited similar binding responses as observed above using the larger EphB2-His protein, demonstrating that the 19-210 fragment of the EphB2 ectodomain is sufficient to mediate dimerization and tetramerization with its cognate Ephrin ligands. The data indicate the arginine residue targeted in loop F’-G of the EphB2-R79P protein is not absolutely essential for binding, though it exhibited a weaker binding response in the sensorgrams compared to EphB2-WT, indicating the electropositive residue does contribute to the interaction. Additionally, like the WT protein, binding of EphB2-R79P to the EphrinB ligands was strongly reduced by tetramer inhibitor A20-I (Figures S6E-S6G). This suggests other electropositive amino acids in the F’-G and K-L loops of the receptor are also involved in Ephrin binding and stability of the tetramer. Nevertheless, as the R79P mutant exhibited reduced binding compared to the WT, we quantified this difference by using the sensorgram data to extract kinetic information. In testing multiple analyte concentrations in the BLI experiments, global fit analysis can yield accurate kinetic rate constants for association and dissociation (k_a o_r k_on a_nd k_dis o_r k_off)_ and provide an equilibrium dissociation constant to describe binding affinity (K_D)_. However, the choice of binding model is crucial. As neither the standard 1:1 (Langmuir) or 2:1 (heterogeneous ligand) binding models accurately capture the complexities of Eph-Ephrin dimer-tetramer dynamics, we used our newly developed model, dimer:tetramer (22), to better evaluate the sensorgram data.

**Figure 7.**
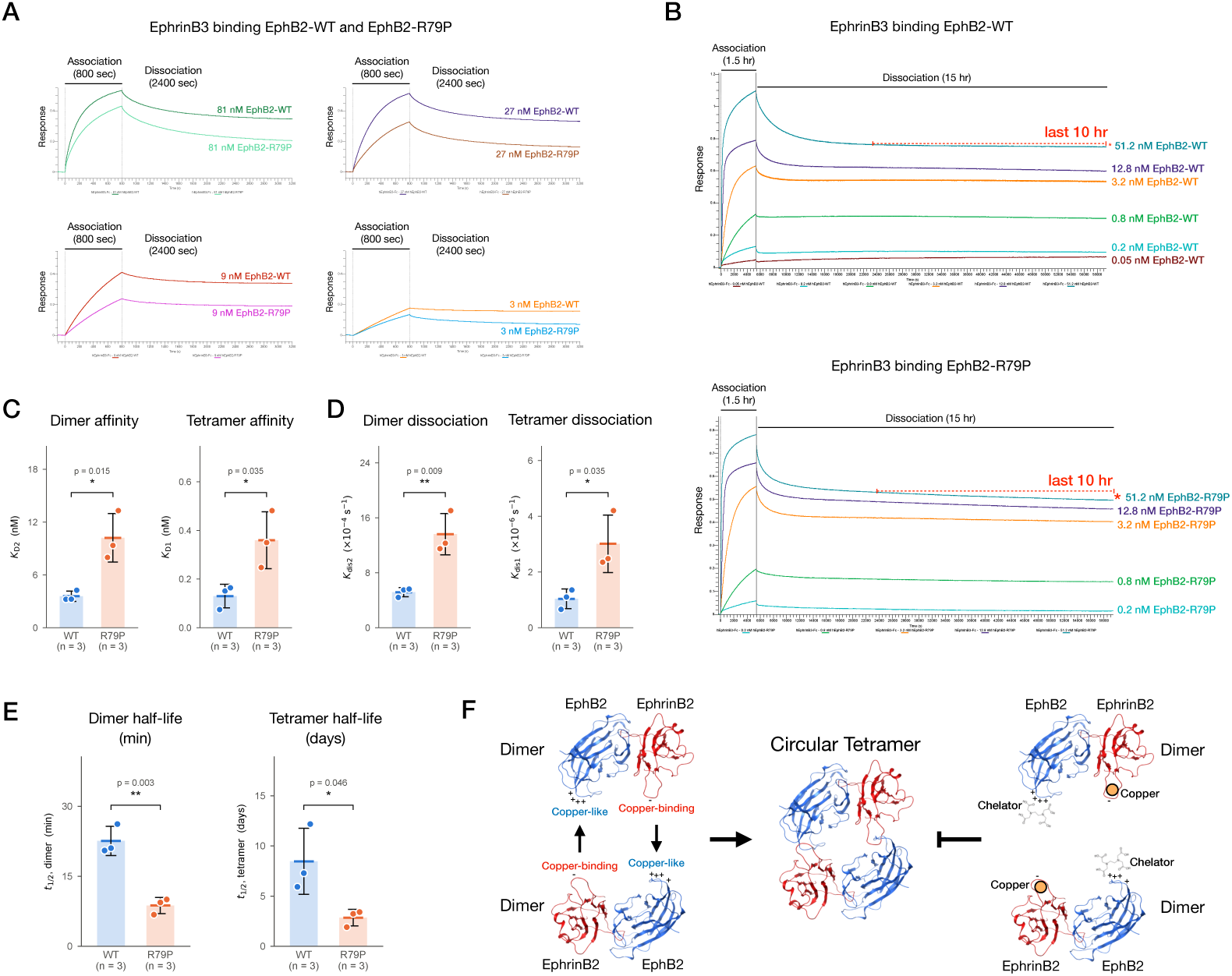
Substituting an arginine in the EphB2 tetramerization F’-G loop with an uncharged amino acid reduces binding to EphrinB3. (A) BLI of immobilized human EphrinB3-Fc binding to 3, 9, 27, and 81 nM of EphB2-WT and EphB2-R79P. R79P targets the residue equivalent to mouse Arg87, which forms a salt bridge with EphrinB2, and showed reduced binding. (B) Extended global fit BLI of EphB2-WT and EphB2-R79P (1.5 hr association, 15 hr dissociation) captured slow tetramer dissociation. The last 10 hr of dissociation at 51.2 nM (red dashed line and asterisk) shows faster tetramer melting for R79P. (C-E) Quantification of global fit sensorgrams from three independent experiments shows WT and R79P differ significantly in binding affinities (C), rates of dissociation (D), and complex stability (E) of both dimer and tetramer. (F) Model of tetramer formation by two EphB2-EphrinB2 heterodimers. Metal chelators are competitive, reversible inhibitors that bind the electropositive F’-G and K-L loops of Eph, whereas copper acts on the electronegative C-D loop of Ephrin. High copper disrupts tetramer formation, but lower concentrations enhance it.

The dimer:tetramer model analyzed the above sensorgrams of EphB2-WT and EphB2-R79P binding to three EphrinB ligands at 3, 9, 27, and 81 nM. In all cases, the tetramer K_D1 i_ndicated the WT protein bound with 2-10 fold higher affinity compared to the R79P protein, with no differences in K_D2 f_or the dimer (Figures S6E-S6G). EphB2-WT tetramer K_D1 v_alues were low or subnanomolar for EphrinB1 (0.19 nM), EphrinB2 (2.16 nM), and EphrinB3 (0.7 nM). The R^2^ values and curve fits for all three interactions using the dimer:tetramer model indicated the model correlated tightly with the sensorgrams, better than if the 1:1 or 2:1 models were employed. Nevertheless, as the 2,400 sec (40 min) dissociation step used in the above studies was too short to accurately capture the off rate of the slowly melting tetramer, additional global fit studies were run that extended the dissociation to 15 hr. Immobilized EphrinB3-Fc was exposed to a wide concentration range of the EphB2-WT and EphB2-R79P proteins from 51.2 nM to as low as 0.05 nM for a 1.5 hr association step and 15 hr dissociation step (Figure 7B). Both proteins showed similar binding responses in the association step with high concentrations of analyte indicating a rapid and steep initial slope of dimer binding with the response bending to plateau as tetramers accumulate. The EphB2-WT protein showed appreciable binding to immobilized EphrinB3 even at as low a concentration as 50 pM in the association step, while EphB2-R79P binding was only detected at 200 pM or above. After sensors were placed into dissociation step, the response initially reversed to indicate rapid melting of dimers, with signals then stabilizing to become a flat line well above baseline. Upon visual inspection of the sensorgrams, a slight but detectable decline occurred throughout the dissociation step for both EphB2 proteins, and isolating the last 10 hr of dissociation indicated the slope of the R79P mutant protein tetramer decline was steeper, as indicated with the red stippled line and asterisk on the runs using 51.2 nM of protein (Figure 7B). Three independent extended global fit runs testing EphB2-WT and EphB2-R79P binding to immobilized EphrinB3 for 1.5 hr association and 15 hr dissociation steps were conducted, kinetic information was obtained using the dimer:tetramer model, and results analyzed as a group. The data indicate the affinities of both the dimer and tetramer are significantly weakened about 3-fold by the R79P mutation, with the tetramer K_D1 g_oing from 0.13 nM to 0.36 nM and dimer K_D2 g_oing from 3.58 nM to 10.2 nM (Figure 7C)(Figure S6H). The reduced affinities of the R79P mutant can be attributed to significantly faster dissociation rates (Figure 7D), and this resulted in a significantly shorter dimer and tetramer half-life (Figure 7E). The tetramer half-life shifted from over 8 days in the WT to less than 3 days in the mutant. Additional extended global fit kinetic studies were also carried out to investigate the effect of the R79P mutant on binding of EphB2 to the cross-subclass EphrinA5 ligand (31), finding here both dimer and tetramer affinities are greater than 4-fold weaker compared to the WT protein (Figure S6I).

Altogether, the data indicate Arg79 in human EphB2 (Arg87 in mouse), one of four electropositive residues in the F’-G and K-L loops (Figure 2A), is not essential but is needed for full binding affinity. Although the crystal structure suggested this arginine forms a salt bridge with Asp69 of EphrinB2, other electropositive residues in the flexible loops can likely do so too. Indeed, the other two arginines present in the F’-G loop are conserved in all the EphB receptors, so we predict those as well as the electropositive lysines and arginines present on the K-L loop of all 14 Eph molecules likely participate in tetramerization. How the R79P mutation affects dimer kinetics is less clear because this residue is not close to the dimerization interface. It is possible a reduced tetramer affinity may indirectly affect dimer stability.

## DISCUSSION

Formation of the Eph-Ephrin circular tetramer is the key step to activate bidirectional signaling when two cells contact. Assembled tetramers also migrate laterally in the plasma membrane and gather into higher-order tetramer superclusters, which amplifies signaling (24, 27, 28, 32–34). Excessive tetramer formation is likely a problem in chronic pathological conditions, as Eph and Ephrin often become overexpressed following tissue insults, as observed for EphB1 in spinal pain neurons following nerve injury (11, 35) or EphB2 in the liver of animals fed a high fat diet (21, 36–38). Rather than searching for traditional tyrosine kinase inhibitors, which only target forward signaling and must counteract an already activated receptor (39–41), we searched for compounds that target the Eph-Ephrin receptor-ligand interaction. The compounds we identified specifically target the Eph-Ephrin tetramer and effectively reduce both forward and reverse signals, appearing more potent than kinase inhibitors (22). Other peptide/small molecule inhibitors target the Eph-Ephrin dimer (42, 43), but require higher concentrations (22). We believe tetramer inhibitors shut down bidirectional signaling so effectively because they target a high-affinity, electrostatically-driven association.

We show Ephrin ligands exhibit a proteinaceous metal-binding like activity that is needed for Eph-Ephrin tetramerization and that metals and chelators affect this interaction. We report here that EphrinB2 binds copper(II) with a K_d o_f 1.5 μM, comparable to or stronger than many Cupredoxins and other copper-binding proteins (64–68). Copper is not needed for tetramer formation, as binding assays are done in metal-free buffers, but it modulates dimer/tetramer dynamics in a biphasic way: at 1-20 μM, around the EphrinB2-copper K_d,_ tetramers and cell signaling increase, whereas at 100 μM the dimer is greatly favored and downstream signaling is reduced. All 14 Eph receptor tetramer interfaces (loops F’-G and K-L) are electropositive clusters of arginine and/or lysine (Figures S2A and S2B), which we propose resemble copper. Eph and Ephrin may act as a molecular rheostat that senses localized copper concentrations at sites of cell-cell contact, increasing signaling (more tetramers) when copper is low and decreasing it (more dimers) when copper is high. The estimated free copper concentration in the body (∼1 μM) is close to the EphrinB2-copper K_d,_ which would suit this hypothetical sensing role.

Consistent with functions in homeostasis, we previously described EphB2-EphrinB2 regulation of the volume, ionic composition, and membrane potential of the inner ear endolymph fluid (44, 45). In the vestibule, EphB2 is expressed in fluid-producing dark cells and EphrinB2 in adjacent transitional cells, and mutating either in mice affected endolymph production and ionic composition, causing severe vestibular dysfunction. To understand this role, we identified protein-protein interactions between the PDZ domain-containing proteins, Pick-1 and Syntenin, that bridge the EphB2 and EphrinB2 proteins with multi-membrane spanning Aquaporin water channels and anion exchangers (44). Water channels facilitate the movement of water molecules, while anion exchangers actively remove or exchange negatively charged ions. Together, this combined action helps drive osmotic flow, which is essential for maintaining cellular hydration and regulating physiological concentrations of ions and pH by coupling the movement of water and ions across cell membranes. We think it possible that our old and new findings together point to a function for Eph-Ephrin interactions in monitoring the extracellular environment and regulating bidirectional signaling strength in response to localized zones of ionic composition at sites of cell-cell contact, where these receptors and ligands are most active, to help maintain homeostasis.

Our findings demonstrate that small molecule chelators can also act as protein-protein interaction inhibitors by interfering with Eph-Ephrin tetramer formation. Chelators typically sequestering the metal ion from a protein, such as using chelation therapy to strip copper from Amyloid-*β* (46). Our data suggest a different activity of chelators, not to sequester a metal needed for a protein’s activity, but to directly dock to the electropositive Eph tetramer interface and compete with Ephrin binding to form the circular tetramer. Thus, without any metal involvement, a chelator affects the ability of one protein (the Eph receptor) to form a protein-protein interaction with another protein (the Ephrin ligand). Our ideas are supported by studies showing EDTA can directly bind within the active site of Farnesyltransferase to inhibit its activity, not to sequester a metal (47). Similarly, EDTA binds the nucleotide-binding pocket of dUTPases and interferes with substrate dUTP binding (48). The K_d v_alues described for EDTA binding to dUTPases ranged from 47-199 nM and are similar to what we determined here for the relatively weak 8-HQ-based primary amine tetramer inhibitor (K_d =_ 31.2 nM / K_i =_ 23.3 nM). The K_i o_f EDTA for disrupting EphB2-EphrinB2, and thus its affinity for EphB2, is below 1 nM, >50-fold stronger than these other chelator-protein interactions.

Summarizing, our data indicate the EphB2-EphrinB2 dimer primarily relies on hydrophobic contacts, while the much higher affinity circular tetramer assembles by forming tandem electrostatic interactions between two juxtaposed Eph-Ephrin dimers. This snaps the two electronegative Ephrin copper-binding tetramer interfaces with the two precisely aligned electropositive copper-like Eph tetramer interfaces (Figure 7F; see Video S22). Once formed, tetramers turn on bidirectional signaling and can gather into tetramer superclusters through lateral migration, amplifying the signals. We believe targeting this electrostatic interaction may be a powerful method to counteract excessive Eph-Ephrin expression and signaling that is now linked to numerous chronic pathological conditions. It has not escaped our attention that chelation therapy has been shown to provide benefit in numerous chronic conditions. As far back as the 1950’s, EDTA was reported to improve chronic conditions in patients suffering from systemic sclerosis (scleroderma). EDTA apparently reduced skin thickening and fibrosis and associated visceral fibrosis of the lung, kidney, and heart (49–51). Chelation therapy is also thought to benefit other chronic fibrotic conditions (52, 53), and recent studies have shown Clioquinol is effective at blunting animal models of pulmonary and renal fibrosis (54, 55). Previously available for decades as an OTC, Clioquinol is one of the 8-HQ compounds we show is a strong Eph-Ephrin tetramer inhibitor (Figure 1C). It is generally thought that the beneficial effects of chelators on chronic conditions are due to removal of excessive, pathological levels of metals. However, in light of our findings, we propose chelators may actually combat these chronic pathologies by targeting Eph-Ephrin tetramers and signaling. Genetic studies show EphB-EphrinB signaling directly participates in chronic inflammation and fibrosis in the skin, kidney, liver, heart, lung, and elsewhere (17–21, 56); knockout/knockdown of EphB2 or EphrinB2 and tetramer inhibitors strongly mitigate fibrosis, whereas gain-of-function signaling enhances it (21). We therefore conclude that chelation therapy benefits in diverse chronic conditions may be attributable to chelators targeting the Eph-Ephrin tetramer. Long-term chelation therapy is not considered viable for chronic conditions, except with excessive metal as in Wilson’s disease or thalassemia, because indiscriminate metal depletion may cause toxicity. While all current Eph-Ephrin tetramer inhibitors are chelators that also bind metal, we believe tetramer-inhibiting activity can be disentangled from the metal-binding activity. Current work will explore these non-metal binding small molecules in anticipation that they will show greater activity and specificity toward the Eph-Ephrin tetramer.

## MATERIALS & METHODS SUMMARY

Eph-Ephrin binding was measured by biolayer interferometry using an Octet RED384 or R8e with Fc- or His-tagged recombinant ectodomains captured on AHC, AHC2, or AR2G biosensors in PBS containing 0.05% Tween-20 at 25°C. Kinetics were analyzed using Octet Studio Analysis 13.1, a custom Python dimer:tetramer model for global fits, and Find50.py for AUC-based IC_50 a_nd Cheng-Prusoff K_i v_alues. Structural modeling used MOE (SiteFinder, induced-fit docking, backbone RMSD, BLOSUM alignment) and 30 ns NAMD simulations with CHARMM36 and MM/PBSA free energies. Copper binding to Alexa Fluor 488-labeled EphrinB2 was measured by microscale thermophoresis, and chelation activity was measured with a pyrocatechol violet colorimetric assay at 632 nm. Cell signaling was tested in Cos1 cells stimulated with pre-clustered EphrinB2-Fc across a copper range (0-160 μM), followed by phosphotyrosine immunoprecipitation and immunoblotting, with all recombinant proteins and antibodies from commercial suppliers.

## ACKNOWLEDGEMENTS

We thank Chad Brautigam for help with MST experiments at the UT Southwestern Macromolecular Biophysics Resource Core, Marc Diamond for use of the Octet RED384 instrument in his laboratory. Patrice Mimche, Dean Sherry, Nick Carruthers, Tracy Saxton, Chad Cowan, and Derrick Rossi provided valuable discussions. A.K. was supported in part by a Herchel Smith summer fellowship. This research was made possible by the U.S. Department of Defense (DOD) / U.S. Army Medical Research and Development Command (USAMRDC) / Congressionally Directed Medical Research Program (CDMRP) awards W81XWH-14-1-0220 and W81XWH-21-1-0949 (to M.H), U.S. Department of Defense (DOD) / Defense Health Agency Contracting Activity (DHACA) award HT9425-26-1-E064 (to M.H.), NIH award R01DK128819 (to M.H.), and an award from the Lyda Hill Philanthropies (to M.H.).

## AUTHOR CONTRIBUTIONS

A.K. designed experiments, conducted BLI assays, collected MST data, and performed NAMD simulations. H.W. made chemicals, conducted MST experiments, helped with experimental design, and assisted in the interpretation of simulation results. M.H. conducted BLI experiments. M.H. conducted BLI studies, helped design and interpret experiments, and provided guidance and insight throughout the study. All authors worked together to write the manuscript.

