## Supplementary Information Appendix for "Proteinaceous Metal-Binding Eph-Ephrin Tetramerization is Modulated by Copper and Chelators"

PNAS

**This PDF file includes:**

Supplementary Materials & Methods (pages 2 - 10)
Supplementary Figures S1 to S6 (pages 11 - 84)
Supplementary Table S1 (page 85)
Supplementary Chemistry Information (pages 86 - 90)
Supplementary Video Legends S1 to S22 (pages 91 - 93)

**Other supporting materials for this manuscript include the following:**

Supplementary Videos S1 to S22

### Materials & Methods

**Biolayer Interferometry**

Eph-Ephrin binding kinetics were analyzed using BLI and either an Octet RED384 or R8e instrument in 384 tilted well plates (Sartorius) and various commercially available recombinant Eph and Ephrin ectodomain proteins that contained C-terminal Fc and/or His tagged epitopes (sources indicated below). Unconjugated human Fc only protein was also utilized in BLI as negative control and showed no binding in the experiments. Reconstituted ectodomain proteins were routinely passed through Zeba desalting columns before use (Thermo Scientific, 89890). The buffer conditions used were PBS pH 7.4 containing 0.05% Tween 20 (PBST), and plates incubated at 25$^{o}$C with shaking at 1,000 RPM. In most experiments, Fc-tagged Eph or Ephrin ectodomains were first sparsely loaded onto Octet Anti-Human IgG Fc Capture biosensor chips (AHC and AHC2, Sartorius, 18-5060 and 18-5143) using 0.25 - 2.0 µg/ml of the Fc protein, and then after washing and obtaining baseline measurements, the biosensors were placed into wells that contained the desired concentration of soluble interacting partner for the association step for the desired period of time, after which the biosensors were returned to the baseline well for dissociation step. In some experiments compounds were sparsely loaded onto Octet Amine Reactive Second-Generation (AR2G) Biosensor Chips (Sartorius, 18-5092) using Octet AR2G Reagent Kit (Sartorius, 18-5095), and then after washing and obtaining baseline measurements, the biosensors were placed into wells that contained the desired concentration of soluble interacting proteins for the association step for the desired period of time, after which the biosensors were returned to the baseline well for dissociation step. In experiments that assessed a tetramer inhibitor compounds, the baseline and association wells also contained the diluted compound in 1% DMSO final as compounds were dissolved and diluted to 100X final concentrations in DMSO prior to being diluted 1:100 in PBST. In experiments where pH was changed to 3.6, biosensors behaved normally and were not damaged. For global fit kinetic studies using multiple concentrations of analyte and a long, extended dissociation step, 96 well flat bottom plates were used with an Octet AE Evaporation Cover (Sartorius, 18-5152). Data acquisition was performed using Octet BLI Discovery 13.1 software to automate the process and generate output files that were then analyzed using Octet Studio Analysis 13.1 software to visualize the sensorgram data.

#### Computer programs

**Find50.py** Was developed to analyze biophysical kinetic data generated by the Octet instruments and calculate the % of a protein-protein interaction that is inhibited at a given concentration of an inhibitor and the $IC_{50}$ value of an inhibitor using area under the curve (AUC) analysis. Find50.py can also be utilized with other types of dose/concentration-dependent data that generates curves needing AUC analysis and $IC_{50}$ calculations. FIND50.py will further use the determined $IC_{50}$ value to calculate the $K_i$ (inhibition constant) using the Cheng-Prussof equation: $K_i = IC_{50}/(1+ [L]/K_D)$ where [L] is the concentration of the soluble analyte used in the assay and $K_D$ is the equilibrium dissociation constant of the binding interaction. The $K_i$ is a dissociation constant specific for the protein-inhibitor complex; a lower value indicates a stronger binding affinity as it indicates a lower concentration of the competitive inhibitor is needed to occupy 50% of its target protein. This equation defines the theoretical relationship between the $IC_{50}$ and $K_i$ of an inhibitory compound based on the $K_D$ of the interaction it is inhibiting and provides a useful constant to describe a specific inhibitor's activity towards its target.

#### Dimer:Tetramer kinetic binding model

Due to the very slow dissociation of the circular tetramer and limitations of the Octet Studio Analysis 13.1 software, a new mathematical model was developed in attempts to better analyze dimer:tetramer binding dynamics from global fit BLI sensorgram traces. A multi-step model was developed to account for the binding situation in which a biosensor immobilized Fc-dimerized Ephrin-Fc protein complex when placed into the association step (1) first binds one soluble Eph interacting molecule to form a single fast-on/fast-off heterodimer to produce an increase in response signal, and then (2) subsequently if a second Eph molecule binds the other tethered immobilized Ephrin-Fc molecule to make a second fast-on/fast-off heterodimer, that also leads to an equal sized increase in response signal, and then (3) if the two closely juxtaposed Eph–Ephrin heterodimers close into a 2:2 circular heterotetramer; closure conserves bound mass and produces no change in response. Because the circular tetramer is extremely stable and dissociates far more slowly than heterodimers, the vast majority of the response contributed by tetramer formation persists through to the end of the dissociation.

$$\mathrm{L} + \mathrm{E} \underset{k_{\mathrm{dis2}}}{\overset{k_{\mathrm{a2}}}{\rightleftharpoons}} \mathrm{LE} \tag{1}$$

$$\mathrm{LE} + \mathrm{E} \underset{k_{\mathrm{dis2}}}{\overset{k_{\mathrm{a2}}}{\rightleftharpoons}} \mathrm{LE}_2 \tag{2}$$

$$\mathrm{LE}_2 \underset{k_{\mathrm{dis1}}}{\overset{k_{\mathrm{a1}}}{\rightleftharpoons}} \mathrm{T} \tag{3}$$

where L is the immobilized Fc-dimerized Ephrin-Fc, E is the soluble Eph, LE and $LE_2$ are the Ephrin-Fc with one and two Eph bound, and T is the closed 2:2 heterotetramer.

Accordingly, the binding events were treated as biphasic, with the faster process always assigned to the heterodimer and the slower to the tetramer (assignment by rate, not amplitude). Reference-subtracted sensorgram data was exported from Octet Data Analysis; with individual sensor data excluded if the response signal was low (peaked at ≤ 0.02 nm) or drifted upward during dissociation. Dissociation phases from all retained sensors were fitted globally to a bi-exponential decay (Eq. 4) with shared rates and per-sensor amplitudes, giving $k_{dis2}$, dimer (fast) and $k_{dis1}$, tetramer (slow).

$$R_i(t) = A_{\mathrm{f},i}\mathrm{e}^{-k_{\mathrm{dis2}}t} + A_{\mathrm{s},i}\mathrm{e}^{-k_{\mathrm{dis1}}t} \tag{4}$$

Association phases were fitted globally to a two-phase model (Eq. 5) with observed rates linear in analyte concentration (Eq. 6) and amplitudes free per sensor; $k_{a2}$, dimer (fast) was taken as the slope of the fast observed rate against analyte concentration.

$$R_i(t) = A_{\mathrm{f},i}\left(1 - \mathrm{e}^{-k_{\mathrm{f,i}}t}\right) + A_{\mathrm{s},i}\left(1 - \mathrm{e}^{-k_{\mathrm{s,i}}t}\right) \tag{5}$$

$$k_{\mathrm{f,i}} = k_{\mathrm{a2}}C_i + k_{\mathrm{f0}}, \qquad k_{\mathrm{s,i}} = k_{\mathrm{a1}}C_i + k_{\mathrm{s0}} \tag{6}$$

where $C_i$ is the analyte concentration of sensor i (in M), $k_{f0}$ and $k_{s0}$ are the rate intercepts, and $k_a$ and $k_{dis}$ are in $M^{-1}$ $s^{-1}$ and $s^{-1}$, respectively.

This slope was reported only if it was bounded on both sides by a 5% sum-of-squares profile and at least three sensors carried a fast phase making up ≥ 10% of their signal. Otherwise $k_{a2}$, dimer (fast) was estimated from the top concentration (Eq. 7) and reported as approximate.

$$k_{\mathrm{a2}} = \frac{k_{\mathrm{fast}} - k_{\mathrm{dis2}}}{C_{\mathrm{top}}} \tag{7}$$

Because the slow observed rate depended only weakly on concentration, $k_{a1}$, tetramer was obtained from a two-exponential fit to the highest-concentration trace (Eq. 8), and was reported only when that fit had $R^2 \geq 0.95$.

$$k_{a1} = \frac{k_{slow} - k_{dis1}}{C_{top}} \tag{8}$$

Equilibrium constants were calculated as in Eq. 9 (both in nM), and half-lives were calculated from the dissociation rate constants as in Eq. 10 (dimer in minutes, tetramer in days).

$$K_{D2} = \frac{k_{dis2}}{k_{a2}} \text{ (dimer)}, \qquad K_{D1} = \frac{k_{dis1}}{k_{a1}} \text{ (tetramer)} \tag{9}$$

$$t_{½,dimer} = \frac{\ln 2}{k_{dis2}}, \qquad t_{½,tetramer} = \frac{\ln 2}{k_{dis1}} \tag{10}$$

All fitting used bounded trust-region least squares in custom Python code (Python 3.13, NumPy 2.3, SciPy 1.16).

**MOE SiteFinder**

All protein structures were first protonated and energy-minimized using the QuickPrep algorithm in Molecular Operating Environment (MOE, Chemical Computing Group ULC). Site prediction was performed using MOE SiteFinder, which employs geometric and empirical algorithms to detect potential ligand-binding pockets across the entire protein surface. Each pocket was evaluated based on the Propensity for Ligand Binding (PLB) score, which quantifies the likelihood and strength of ligand accommodation at a given site.

**MOE Docking**

Ligand structures were constructed in MOE's Builder module and subjected to protonation and minimization using MOE QuickPrep. EphB2 monomer (PDB code: 1NUK) was also prepared using MOE QuickPrep. Docking simulations were performed using the MOE Docking with the GBVI/WSA dG scoring function to estimate the binding free energy (ΔG) as the docking score. For each protein–ligand pair, five simulations were performed: each simulation contains 30 poses per ligand generated under induced-fit conditions to account for receptor flexibility. The best-ranked poses were retained (default MOE Docking Settings were used). The docking results were outputted as interaction tables denoting all amino acid and ligand interactions as well as docking scores.

**MOE RMSD Backbone Superimposition**

All proteins were protonated and minimized using QuickPrep prior to alignment. Structural superimposition was carried out in MOE using the Superimposition tool to assess conformational similarity between various proteins or between unbound and ligand-bound configurations. Root-Mean-Square Deviation (RMSD) values were calculated over backbone atoms, and resulting superimpositions were visualized and analyzed to quantify structural conservation and conformational shifts. A separate MOE script was utilized to quantify specific amino acid side chain conformational changes (RMSD).

**BLOSUM Sequence Similarity**

Amino acid sequence similarity among protein variants was quantified using MOE's BLOSUM (Blocks Substitution Matrix) alignment algorithm. All structures were first preprocessed through QuickPrep to ensure residue consistency with the modeled structures. Pairwise sequence alignments were generated, and similarity scores were computed to assess evolutionary conservation and residue-level correspondence across the analyzed proteins.

**Nanoscale Molecular Dynamics (NAMD)**

Protein structures were extracted and prepared using MOE QuickPrep algorithm for protonation and pH adjustment at 7.4, adding solvent water and 0.137 M NaCl to the system to mimic physiological salt concentrations. The system was energy-minimized using MOE Minimization and then batch instructions were created and transferred to the Southwestern BioHPC P100GPU cluster. NAMD simulations were run using the CHARMM36 force field and explicit solvent models for up to 5 days. Simulations included 200 ps minimization, 100 ps heating, 100 ps NVT (canonical ensemble) phase, 100 ps NPT (isothermal-isobaric ensemble) phase, followed by the 30 ns simulation. Once complete, trajectory files were exported and analyzed with the topology files in MOE MD Analysis to determine RMSD values, distance apart from atoms/centroids, and Molecular Mechanics/Poisson-Boltzmann Surface Area (MM/PBSA) binding free energy ($\Delta G$) estimates of solvation energy in kcal/mol.

**Microscale Thermophoresis**

EphrinB2-metal interactions were analyzed using a Monolith NT.115 Microscale Thermophoresis system (Nano Temper Technology). An untagged version of human EphrinB2 ectodomain protein (Sino Biologicals, 10881-HCCH) was reconstituted in PBS and passed through Zeba desalting columns (Thermo Scientific, 89890) into a buffer condition of 50 mM HEPES buffer with 150 mM NaCl. The protein was labeled with Alex Fluor 488 dye (Succinimidyl Ester, from Invitrogen, #A2000) that efficiently coupled to the ε-amino groups of lysines under slightly basic conditions (pH ~8.3). After incubation, unreacted dye was removed by size-exclusion column. The labeling ratio of dye to protein was determined using Cary 60 UV-Vis spectrophotometer (Agilent Technologies). In the standard MST assay, 10 μl of increasing amounts of an experimental metal ($CuCl_2$, $CoCl_2$, or $NiCl_2$) were combined with 10 μl of dye labeled protein. After incubation in dark for 30 min, the samples were transferred to capillary tubes and subjected to MST system for fluorescence scan and test. The final concentration of an experimental metal ranged from 0 to 312.5 μM, and the final concentration of dye labeled protein was 57 nM. Data acquisition is performed using the Monolith NT.115 Microscale Thermophoresis instrument and its associated customizable software to automate the process and generate output files for analysis using PALMIST software and GUSSI software to visualize the thermophoresis data and calculate the important kinetic parameters of the molecular interaction under study. To supplement MST data, the SD-Test (SDS denaturation test) is a control assay used in binding studies to confirm that fluorescence changes are due to specific ligand–protein interactions rather than non-specific artifacts such as aggregation or adsorption. In this test, samples of free protein and protein–metal complexes are mixed with an SDS/DTT solution and heated at 95°C to fully denature the proteins. If the fluorescence levels of all samples equalize after denaturation, the original changes can be attributed to specific binding; if differences persist, they likely result from non-specific effects. This test is recommended when a ligand causes a fluorescence shift greater than ±20% and provides a straightforward way to validate interaction data. In the case of copper binding to EphrinB2, SD tests confirm that fluorescence quenching was due to copper interacting with EphrinB2 as fluorescence was recovered after SD test in the 3rd highest concentration consistently.

***In vitro* copper chelation assay**

In the standard chelation assay, 30 µl of increasing amounts of an experimental chemical compound are combined with 30 µl of 0.1µg/µl $CuSO_4$, 50 µl of 0.34 mM pyrocatechol violet (PV), and DMSO (if necessary depending on compound solubility) in 155 µl 1X AOX buffer (ZenBio, AOX-16; or Henkemeyer laboratory made, 5 mM potassium phosphate, pH 6.1 with 0.9% sodium chloride) in triplicate wells of an assay plate. This colorimetric assay is based on the metal chelator activity of PV that when complexed with copper is bluish-violet in color, and that addition of a competing chelator will remove metal from PV and lead to loss of blue-violet color. As the assay is only accurate if the compound under study dissolves well, DMSO may be included to increase solubility and achieve clear and transparent solutions. Negative control wells contain PV without $CuSO_4$, positive control wells have PV+$CuSO_4$ but no added experimental chemical compound. Absorbance was measured at 632 nm.

**Cell stimulations, protein lysates, and immunoprecipitations**

Cos1 cells were grown in 6 well culture plates in DMEM medium containing 10% fetal bovine serum, 2 mM L-Glutamine and antibiotics (1% penicillin and streptomycin). Serum-starved cells were stimulated with 1.5 µg/ml (30 nM) pre-clustered mouse EphrinB2-Fc (to stimulate forward signaling) for 32 minutes in protein free DMEM supplemented with 10 mM HEPES and containing increasing concentrations of $CuSO_4$ (0-160 uM). For pre-clustering, EphrinB2-Fc protein was combined with goat anti-human IgG at a 2:1 ratio for at least 1 hr prior to cell stimulations. Following stimulations, plates were placed on ice, cells washed with cold PBS, and then lysed using 1 ml/well ice cold PLC Lysis Buffer (50 mM HEPES (pH 7.5), 150 mM NaCl, 1.5 mM $MgCl_2$, 10% Glycerol, 1% Triton-X100, 1 mM EGTA, 10 mM $Na_4P_2O_7 \cdot 10H_2O$, 10 mM NaF, 1 mM $Na_3VO_4$, 1X Halt Protease Inhibitor Cocktail, 1 mM PMSF). After nutating the lysate for 30 min in a cold room, the insoluble material was pelleted, and cleared supernatants frozen on dry ice and placed in -80 $^{o}$C freezer for long-term storage.

For immunoprecipitations, 1 ml protein lysates prepared in advance from desired cell stimulations were thawed on ice and after removing a small sample for the immunoblot lysate lanes 20 µl of bead-conjugated pTyr1000 antibody was added and allowed to Nutate in a cold room for 2 hr to precipitate all phosphotyrosine-containing proteins. Beads were washed 3X in ice cold low salt HNTGV (20 mM HEPES, pH 7.5, 50 mM NaCl, 0.02% Triton-X100, 10% Glycerol, 1 mM $Na_3VO_4$), and proteins eluted by adding 13 µl 4X SDS Sample Buffer containing DTT and heating at 65 $^{o}$C for 5 minutes, resolved on an SDS-PAGE gel, transferred to membrane, and immunoblotted with indicated antibodies.

**Proteins**

**Recombinant Eph/Ephrin ectodomain proteins:**

Human Fc protein (R&D Systems, 110-HG-100). Is a 26.6 kDa protein (with human Fc Pro100-Lys330) that migrates as a 30-35 kDa protein in SDS-PAGE under reducing conditions.

Human EphrinB2-Fc (R&D Systems, 7397-EB-050). Is a 48.8 kDa Fc fusion protein (with human EphrinB2: Met1-Ala229) that migrates as a 58-66 kDa protein in SDS-PAGE under reducing conditions.

Rat EphB1-Fc protein (R&D Systems, 1596-B1-200). Is an 85 kDa Fc fusion protein (with rat EphB1: Met18-Gln538) that migrates as a 102 kDa protein in SDS-PAGE under reducing conditions.

Mouse EphB2-Fc-His protein (R&D Systems, 467-B2-200). Is an 85 kDa Fc fusion protein (with mouse EphB2: Val27-Lys548) that migrates as a 100-110 kDa protein in SDS-PAGE under reducing conditions.

Human EphB3-Fc (R&D Systems, 5667-B3). Is an 82.7 kDa Fc fusion protein (with human EphB43: Leu38-Ala550) that migrates as a 90-100 kDa protein in SDS-PAGE under reducing conditions.

Human EphB4-Fc (R&D Systems, 11307-B4). Is an 85.2 kDa Fc fusion protein (with human EphB4: Leu16-Arg539) that migrates as a 95-108 kDa protein in SDS-PAGE under reducing conditions.

Human EphrinB2-His protein (Sino Biologicals, 10881-H08H). Is a 23.5 kDa His fusion protein (with human EphrinB2: Met1-Ala229) that migrates as a 35-40 kDa protein in SDS-PAGE under reducing conditions.

Mouse EphrinB2-His protein (Sino Biologicals, 50598-M08H). Is a 23.5 kDa His fusion protein (with mouse EphrinB2: Met1-Ala232) that migrates as a 30-40 kDa protein in SDS-PAGE under reducing conditions.

Human EphB1-His (Sino Biologicals, 11963-H08H). Is a 60 kDa His fusion protein (human EphB1: Met1-Pro540) that migrates as a 66 kDa protein in SDS-PAGE under reducing conditions.

Human EphB2-His (Sino Biologicals, 10762-H08H). Is a 59.7 kDa His fusion protein (human EphB2: Met1-Leu543) that migrates as a 70 kDa protein in SDS-PAGE under reducing conditions.

Human EphB4-His (Sino Biologicals, 10235-H08H). Is a 58.5 kDa His fusion protein (human EphB4: Met1-Ala539) that migrates as a 72 kDa protein in SDS-PAGE under reducing conditions.

**Primary Antibodies:**
Rabbit anti-phosphotyrosine MultiMab™ Rabbit mAb mix (pTyr1000) (Cell Signaling, 8954). A mixture of rabbit monoclonal antibodies that recognizes a broad range of tyrosine-phosphorylated proteins and peptides, and does not cross-react with proteins or peptides containing phospho-Ser or phospho-Thr residues.

**Secondary Antibodies and detection reagents:**
Goat anti-Rabbit HRP secondary (Jackson ImmunoResearch, 111-035-144). Peroxidase AffiniPure Goat Anti-Rabbit IgG (H+L) (min X Hu, Ms, Rat Sr Prot).

ELISA Pico Chemiluminescent Substrate (Thermo Scientific, 37069). ELISA Pico Chemiluminescent Substrate is optimized to generate an intense light signal and provide exceptional performance in luminometer-based assays.

SuperSignal West Dura Extended Duration Substrate (Thermo Scientific, 34076). A luminol-based enhanced chemiluminescence (ECL) horseradish peroxidase (HRP) substrate ideal for quantitative western blotting with stable light output for mid-femtogram level detection for western blot analysis.

### Supplementary Figures

8-Hydroxyquinoline based small molecules and EDTA are copper chelators

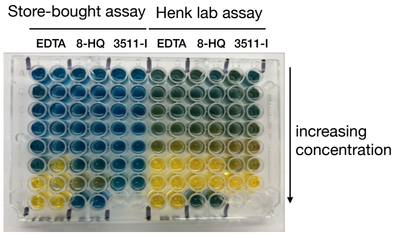


| Store-bought assay | | | | | | Henk lab assay | | | | | |
|---|---|---|---|---|---|---|---|---|---|---|---|
| EDTA | | 8-HQ | | 3511-I•.2xHCl | | EDTA | | 8-HQ | | 3511-I•2xHCl | |
| 1 | 2 | 3 | 4 | 5 | 6 | 7 | 8 | 9 | 10 | 11 | 12 |
| 1.423 | 1.286 | 1.37 | 1.213 | 1.363 | 1.227 | 0.819 | 0.838 | 0.828 | 0.915 | 0.86 | 1.001 |
| 1.121 | 1.425 | 1.342 | 1.273 | 1.298 | 1.283 | 0.867 | 0.811 | 0.867 | 0.88 | 0.869 | 0.844 |
| 1.339 | 1.363 | 1.215 | 1.368 | 1.306 | 1.337 | 0.66 | 0.829 | 0.808 | 0.872 | 0.807 | 0.874 |
| 1.14 | 1.187 | 1.359 | 1.275 | 1.385 | 1.264 | 0.608 | 0.634 | 0.721 | 0.615 | 0.833 | 0.693 |
| 1.143 | 1.105 | 1.219 | 1.174 | 1.34 | 1.296 | 0.482 | 0.326 | 0.425 | 0.457 | 0.48 | 0.458 |
| 0.906 | 0.154 | 0.983 | 0.944 | 1.377 | 1.346 | 0.121 | 0.142 | 0.289 | 0.244 | 0.515 | 0.537 |
| 0.132 | 0.169 | 0.587 | 0.579 | 1.431 | 1.427 | 0.116 | 0.123 | 0.204 | 0.213 | 0.131 | 0.13 |

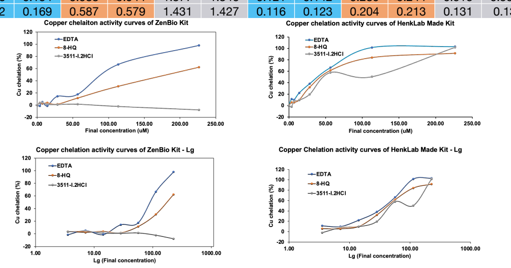


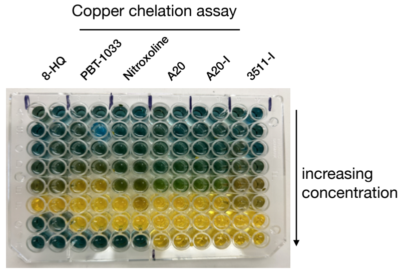


| 8-HQ | | PBT-1033 | | Nitroxoline | | A20•2xHCl | | A20-I•2xHCl | | 3511-I•2xHCl | |
|---|---|---|---|---|---|---|---|---|---|---|---|
| 1 | 2 | 3 | 4 | 5 | 6 | 7 | 8 | 9 | 10 | 11 | 12 |
| 0.712 | 0.918 | 0.829 | 0.833 | 0.894 | 0.981 | 0.817 | 0.87 | 1.024 | 0.868 | 0.815 | 1.012 |
| 0.979 | 0.812 | 0.675 | 0.889 | 0.866 | 0.909 | 0.769 | 0.964 | 0.87 | 0.825 | 0.875 | 0.899 |
| 0.809 | 0.695 | 0.83 | 0.728 | 0.788 | 0.812 | 0.749 | 0.849 | 0.885 | 0.869 | 0.721 | 0.992 |
| 0.711 | 0.725 | 0.553 | 0.56 | 0.708 | 0.707 | 0.797 | 0.731 | 0.772 | 0.821 | 0.67 | 0.731 |
| 0.479 | 0.466 | 0.207 | 0.194 | 0.519 | 0.513 | 0.565 | 0.605 | 0.586 | 0.584 | 0.459 | 0.466 |
| 0.185 | 0.207 | 0.12 | 0.115 | 0.118 | 0.226 | 0.106 | 0.121 | 0.128 | 0.13 | 0.475 | 0.498 |
| | | 0.129 | 0.135 | 0.109 | 0.117 | 0.089 | 0.097 | 0.102 | 0.102 | 0.113 | 0.116 |
| 0.888 | 0.885 | 0.903 | 0.89 | 0.886 | 0.897 | 0.107 | 0.118 | 0.11 | 0.108 | 0.107 | 0.108 |
| Positive control (No compounds, include $CuSO_4$ and PV) Maximum absorbance value | | | | | | Negative control (No compounds, no $CuSO_4$, include PV) Background absorbance value | | | | | |

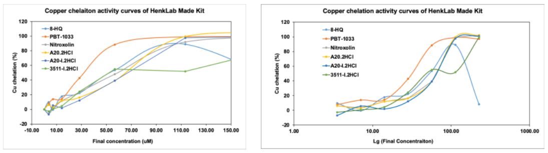


**Figure S1A.** *In vitro* copper chelation assays indicate 8-hydroxyquinoline based small molecules including 8-hydroxyquinoline (8-HQ), PBT-1033, Nitroxoline, A20, A20-I, and 3511-I as well as commonly used metal chelators EDTA and EGTA are copper chelators. (Left) A comparison between a store-bought copper chelation assay kit with one made in Henkemeyer laboratory demonstrates the home-made assay that uses AOX buffer containing 30% DMSO is more robust. A photograph of the *in vitro* copper chelation assay plate showing effect of increasing amounts of EDTA, 8-HQ, and 3511-I·2xHCl salt, the actual numerical data after scanning the plate for absorbance at 632 nm, and the resulting graphs of the data are shown. Copper chelation activity for all three compounds (EDTA, 8-HQ, and 3511-I) was detected with 3511-I showing less activity than EDTA or 8-HQ. (Right) Using the home-made copper chelation kit, 8-HQ based small molecules were further studied to show the effect of increasing amounts of 8-HQ, PBT-1033, Nitroxoline, A20·2xHCl, A20-I·2xHCl, and 3511-I·2xHCl. All six compounds showed copper chelation activity, with PBT-1033 exhibiting greater activity and 3511-I showing slightly less activity.

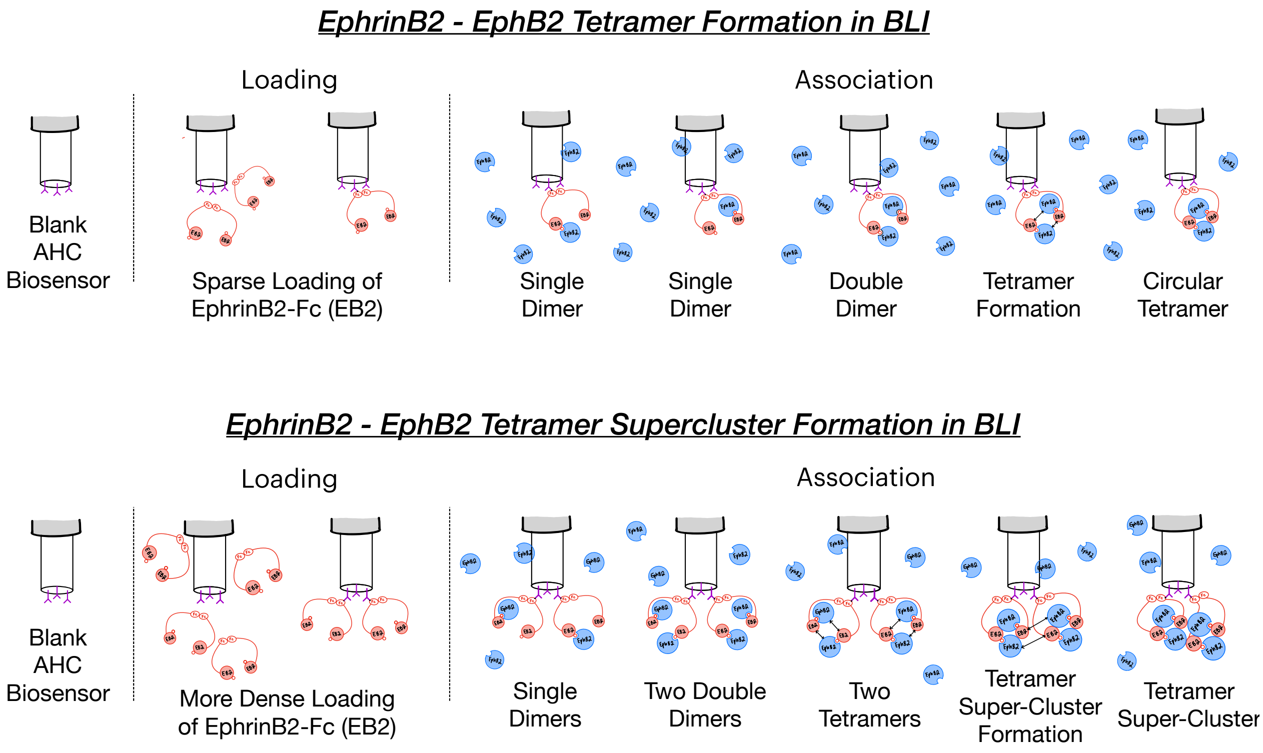


**Figure S1B.** Cartoons visualizing how biolayer interferometry (BLI) can be used to study the complex Eph-Ephrin receptor-ligand protein-protein binding interactions that lead to formation of dimers, tetramers, and tetramer superclusters. Here, on the **top drawing** as example, low concentrations of soluble EphrinB2 ectodomain available as a fusion protein with the Fc region of human IgG1 attached to the C-terminus (EphrinB2-Fc or simply EB2 in the cartoon, red) is first sparsely loaded in buffer (PBST: PBS containing 0.05% Tween-20) onto AHC or AHC2 Anti-Human Fc Capture sensorchips to immobilize the "bait" protein using an Octet RED384 (RH16) or R8e instrument (Sartorius). As the Fc domain forms a strong Fc-Fc homodimer in solution, sparse loading leads to an isolated island of two closely positioned EphrinB2-Fc ectodomains. After washing the EphrinB2-Fc loaded biosensors and obtaining baseline measurements in buffer, each biosensor is then placed in a well containing the desired soluble "prey" target protein (e.g. the EphB2 ectodomain protein fused to a poly-Histidine tag at the C-terminus, EphB2-His, blue) which is diluted to a specific concentration in buffer for a timed association step. As illustrated, each of the two available EphrinB2 ectodomain "baits" can independently bind to an EphB2 ectodomain "prey" using the dimerization interface, which leads to an increase in sensorgram response signal. At some point, both of the immobilized EphrinB2 proteins may each form a dimer with available EphB2 molecules, forming two adjacent dimers, and when that happens it now becomes possible to form the circular tetramer through the bringing together of the two distinct juxtaposed Eph-Ephrin tetramerization interfaces. After the association step, biosensors are then placed back into buffer only for a timed dissociation step. Here, the dimers, which exhibit fast-on/fast-off binding kinetics, rapidly dissociate/melt with EphB2 being released from the immobilized EphrinB2, reducing biosensor response signal and leaving behind the stable slow-on/slow-off circular tetramer macromolecular structure. As the circular tetramer is stable and can persist through to the end of the dissociation step, the sensorgram response signal remains well above that of baseline at the end of the experiment. On the **bottom drawing** is an illustration of what may occur when EphrinB2-Fc "bait" protein is loaded at higher density on the biosensor surface where two Fc-Fc homodimers become immobilized to occupy nearby adjacent regions of the chip. Here, dimer and circular tetramer formation proceed as above, but now with two adjacent circular tetramers nearby, the protein complexes can interact and assemble through other Eph-Ephrin contact interfaces to form tetramer superclusters, which are extremely stable macromolecular complexes less prone to be targeted by tetramer inhibitors (insurmountable). The cartoons shown are simplified and do not take into account that soluble EphB2 and EphrinB2 ectodomains can also each assemble into homodimers. Reference biosensors loaded with the human Fc only protein or blank loaded with no protein were included in the BLI studies to control for any non-specific binding and typically showed no response signal (or only slight signals) when exposed to an Eph-His protein during the association step, compared to sensors loaded with an EphrinB-Fc protein which exhibited obvious strong response signals. Biosensors loaded with EphrinB-Fc protein but only exposed to buffer only during the association step were used to control for and subtract baseline drift. These control EphrinB-Fc loaded sensors typically exhibited little if any response drift, indicating protein remained stably immobilized onto the AHC/AHC2 chips throughout the baseline, association, and dissociation steps.

Copper chelators A20-I, Clioquinol, EDTA, & EGTA disrupt EphB-EphrinB2 tetramerization

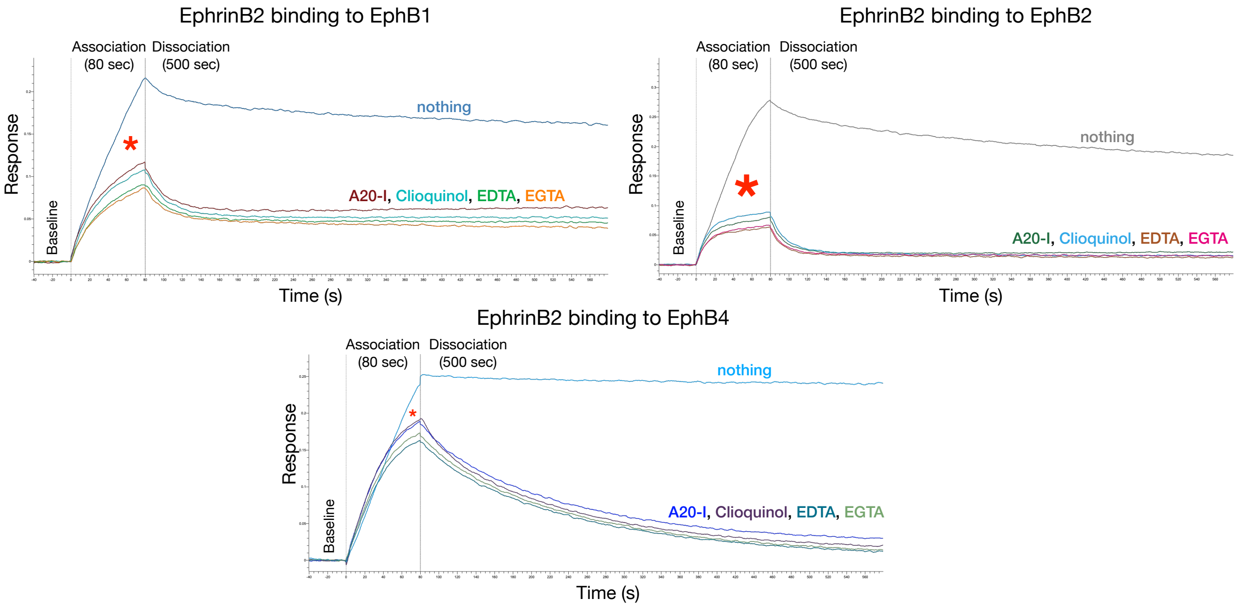


**Figure S1C.** Shown are sensorgrams of Biolayer Interferometry (BLI) studies used to characterize the effects of 8-HQ, Clioquinol, EDTA, and EGTA chelators on EphB1-EphrinB2, EphB2-EphrinB2, and EphB4-EphrinB2 binding kinetics and tetramer formation. The human EphrinB2-Fc ectodomain was sparsely immobilized onto AHC biosensors and then after baseline measurements was exposed to 25 nM of soluble human EphB1-His, EphB2-His, or EphB4-His ectodomain for an 80 sec association step followed by a 500 sec dissociation step, either without any tetramer inhibitor added (nothing), or testing the effect of adding a compound of interest, here testing 3.2 μM of the copper chelators A20-I, Clioquinol, EDTA, and EGTA. The data indicates all four of the compounds tested were able to specifically inhibit formation of the tetramer during the association step (red asterisks), with the typical strongest effect on the tetramer-driven EphB2-EphrinB2 interaction and the weakest effect on the dimer-driven EphB4-EphrinB2 interaction. Note that presence of tetramer inhibitor does not affect the initial slope of response at the start of association that represents the fast-on/fast-off kinetics of dimer binding, but rather severely curtails the assembly of tetramers, which are super stable and persist throughout the dissociation in the absence of any added metal chelator.

In all BLI studies, for each biosensor, the indicated concentration of compound was included throughout the run (in baseline, association, and dissociation wells/steps) to avoid potential changes in response during step changes, and the buffer condition used was always PBST (PBS + 0.05% Tween-20) with 1% DMSO (used for compound dilutions), unless otherwise noted. BLI experiments typically include a biosensor with 2 μM A20-I compound to serve as the maximum (Max) tetramer inhibitor control to compare with the biosensor run without any compound (nothing, DMSO only). BLI experiments using a concentration-series of a compound of interest are used to determine the relative tetramer-specific half maximal inhibition concentration ($IC_{50}$) of a tested compound as calculated by area under the curve (AUC) analysis of each of the different concentration runs as fully described in (Wang, Khambete, et al., submitted). The inhibition constant ($K_i$) can further calculated using a modified Cheng-Prusoff equation that describes the relationship between the $IC_{50}$ value of a competitive inhibitor compound, the concentration the soluble protein it is competing with, and the dissociation constant ($K_D$) of the protein-protein interaction that is disrupted. Only $K_i$ values determined using the EphB2-EphrinB2 interaction are provided. Without any compound, the kinetics of immobilized EphrinB2 binding to EphB2 exhibited a complex pattern of dimer and tetramer binding, while BLI runs with sufficient concentrations of a tetramer inhibitor, as exemplified by the Max control runs, show mainly dimer binding kinetics with greatly reduced formation and accumulation of the high-affinity and very stable tetramer complex.

Listed FDA approved drugs do not disrupt Eph-Ephrin tetramerization

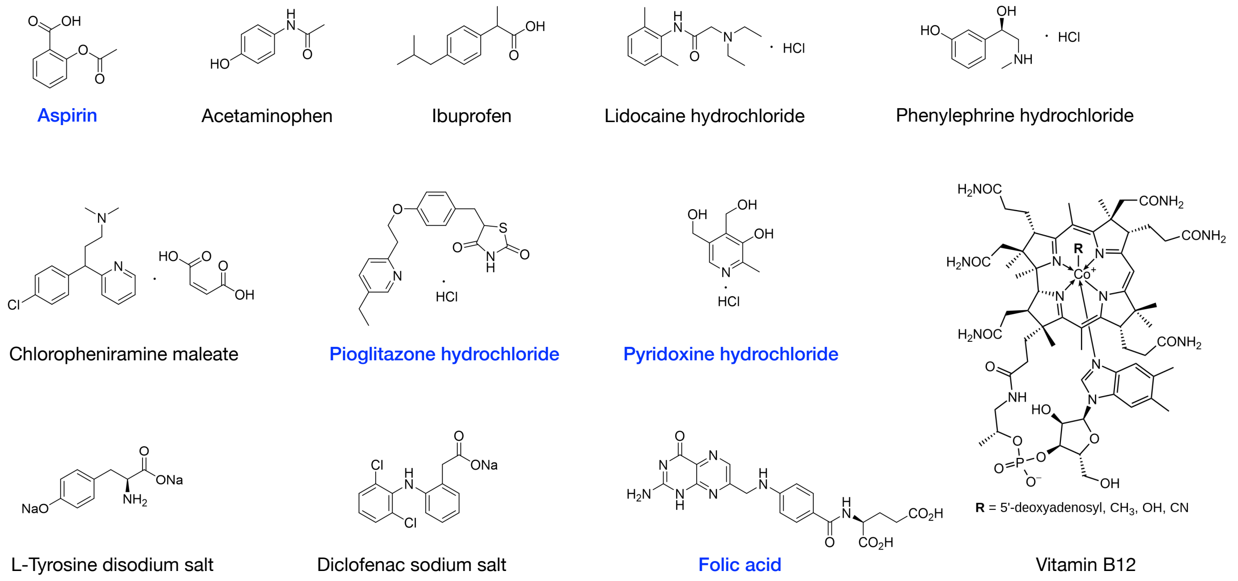


**Figure S1D.** *In vitro* copper chelation assay indicates aspirin, pioglitazone hydrochloride, pyridoxine hydrochloride, and folic acid are copper chelators (in blue). BLI studies of EphB2-EphrinB2 interactions indicates that none of the purchased FDA approved drugs target the interaction or tetramer formation at concentrations up to 100 μM.

#### Common metal chelators disrupt Eph-Ephrin tetramerization

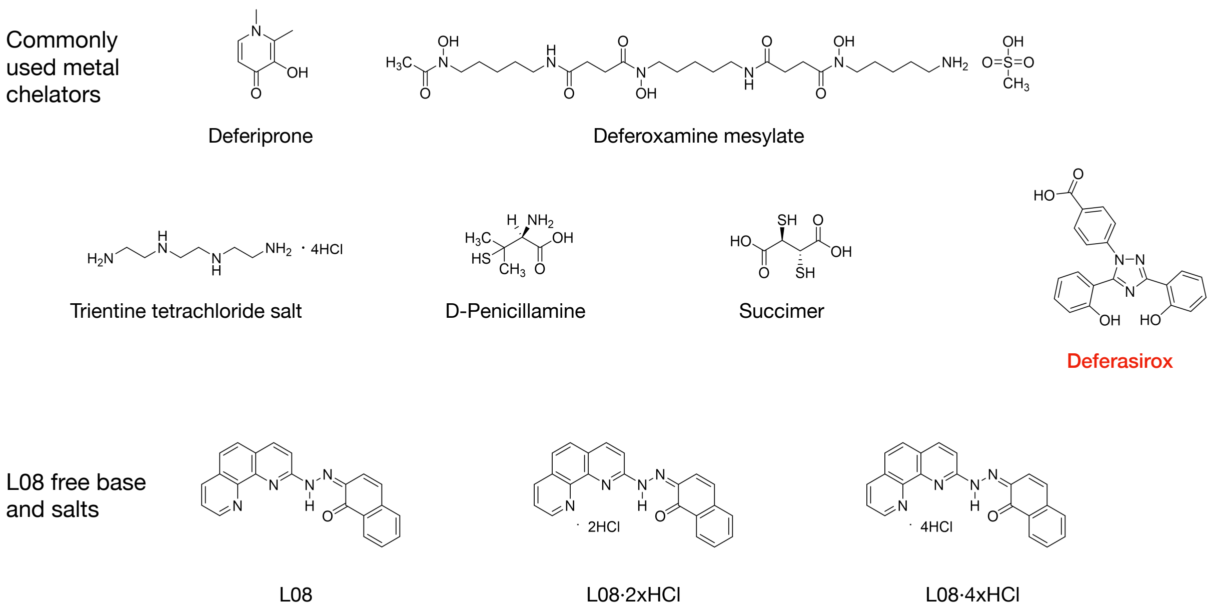


**Figure S1E.** *In vitro* copper chelation assay indicates that commonly used metal chelators and L08 compounds are copper chelators. BLI studies of EphB2-EphrinB2 interactions indicates that these compounds can target tetramers, except deferasirox (in red).

### Copper disrupts EphB-EphrinB tetramerization

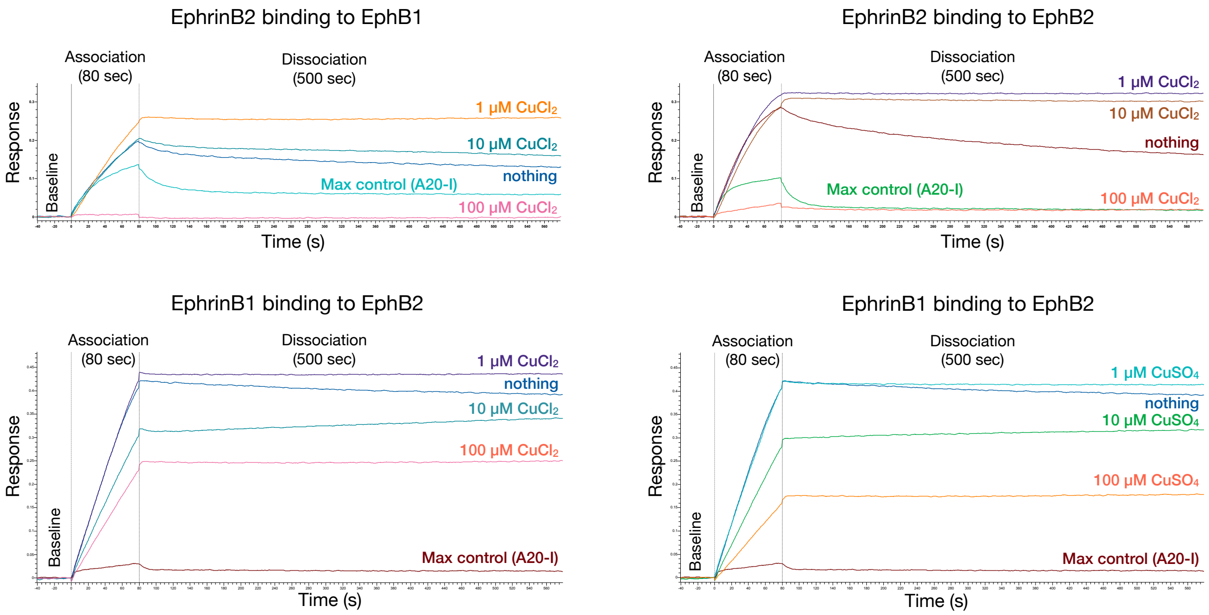


**Figure S1F.** Additional BLI studies of EphB1-EphrinB2, EphB2-EphrinB2, and EphB2-EphrinB1 indicate copper affects dimer/tetramer dynamics. AHC immobilized human EphrinB2-Fc ectodomain binding to 25 nM soluble human EphB1-His and EphB2-His ectodomains (top), as well as AHC immobilized human EphrinB1-Fc ectodomain binding to 25 nM soluble human EphB2-His ectodomain (bottom). Compared to the striking biphasic effect of copper on EphrinB2 dimer/tetramer dynamics, the effect of copper on EphrinB1 binding was more muted, though still clearly causing a concentration-dependent loss of tetramer accumulation. The data further shows copper disrupts tetramers irrespective of whether copper chloride or copper sulfate was used in the binding experiment.

Cobalt and nickel do not disrupt EphB-EphrinB2 tetramerization

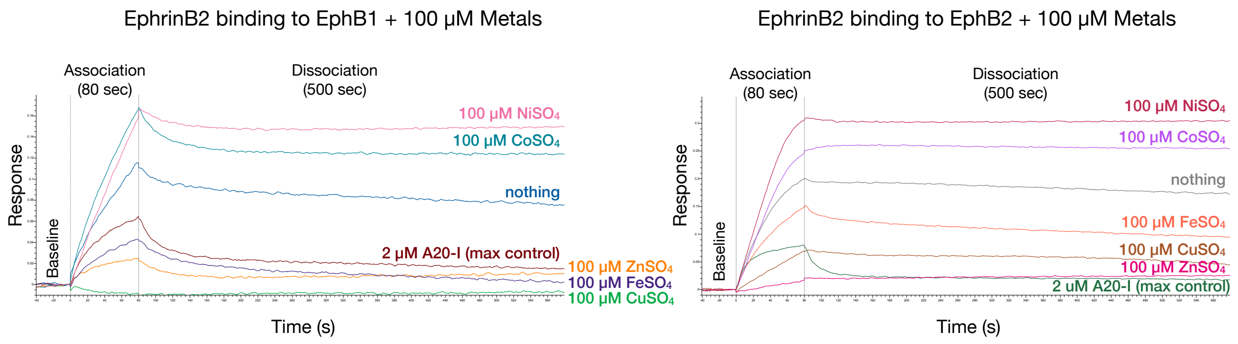


**Figure S1G.** Additional BLI studies of AHC immobilized human EphrinB2-Fc ectodomain binding to 25 nM human EphB1-His and EphB2-His ectodomains show 100 µM concentrations of cobalt or nickel do not reduce binding interactions or formation of stable tetramers, if anything they enhance the interactions, while copper, zinc, and iron all effectively target the interactions and reduce formation of tetramers.

#### EphrinB2 stimulated EphB2 forward signaling in Cos1 - testing copper

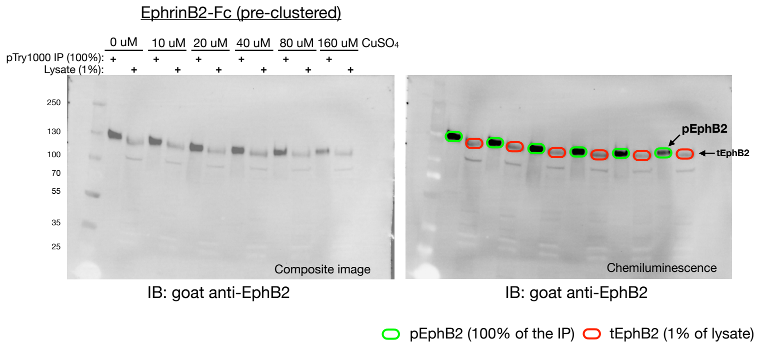


Gels 245E - Band analysis with % pEphB2 - CuSO4

| Label | Volume (Int) | Adj. Vol. (Int) | %Phosphorylated |
|---|---|---|---|
| pEphB2 - No Copper | 86108740 | 54904259 | 4.221670 |
| tEphB2 - No Copper | 52460246 | 12456300 | |
| pEphB2 - 10 uM Copper | 79386262 | 50899363 | 3.782184 |
| tEphB2 - 10 uM Copper | 51841924 | 12948671 | |
| pEphB2 - 20 uM Copper | 97298643 | 64722191 | 5.496527 |
| tEphB2 - 20 uM Copper | 52124525 | 11127884 | |
| pEphB2 - 40 uM Copper | 86132737 | 50240942 | 3.058187 |
| tEphB2 - 40 uM Copper | 55237733 | 15925935 | |
| pEphB2 - 80 uM Copper | 68586142 | 35374324 | 3.248767 |
| tEphB2 - 80 uM Copper | 46834054 | 10534796 | |
| pEphB2 - 160 uM Copper | 50291400 | 17170395 | 1.484547 |
| tEphB2 - 160 uM Copper | 40657883 | 11394381 | |

$$\text{\% pEphB2} = \frac{\text{\# phospho-photons}}{\text{\# photons total}} = \frac{\text{pEphB2}}{\text{pEphB2} + (\text{tEphB2} \times 100)}$$

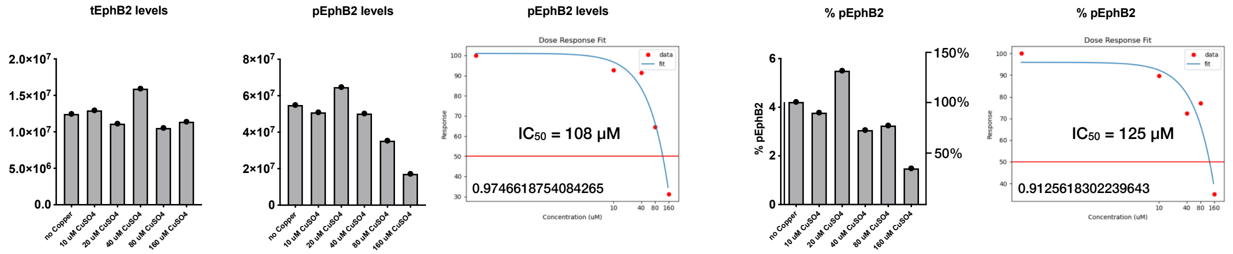


**Figure S1H.** $CuSO_4$ modulates EphB2 forward signaling in live cells. Increasing concentrations of $CuSO_4$ (10-160 μM) were added to serum-starved Cos1 cells that endogenously express EphB2 receptor and stimulated with 1.5 μg/ml (30 nM) pre-clustered EphrinB2-Fc for 32 minutes. Protein lysates were incubated with pTyr1000 beads to immunoprecipitate all phosphotyrosine-containing proteins and 100% of the eluted proteins were run in an SDS-PAGE gel next to 1% of the starting lysate, and then immunoblotted for EphB2. This revealed in the absence of copper or at low concentrations of copper a strong phospho-EphB2 signal in the IP lanes (pEphB2, green circled bands) compared to the whole cell lysate lane for total EphB2 expression (tEphB2, red circled bands). Note that the pEphB2 band at just above the 130 KDa protein standard exhibited a slightly reduced migration in the SDS-PAGE gel due to its phosphorylation compared to the tEphB2 band, which migrated just below the 130 KDa standard. Shown are both composite (to visualize the pre-stained protein standards) and chemiluminescent images of the data. Band quantification showed at 10 μM $CuSO_4$ there was perhaps a slight decrease in the level of pEphB2 (green circled bands) compared to no added copper, while at 20 μM the level of pEphB2 actually increased compared to no copper, which then reversed with more copper such that at 80 μM and 160 μM $CuSO_4$ there was an obvious reduced pEphB2 signal. The band quantification for total tEphB2 levels in the lysate lanes indicated no differences in the overall level of receptor protein between the treatment groups. Using the equation shown, the pEphB2 and adjacent tEphB2 signals can be used to precisely quantify the percentage of EphB2 in the cells from each treatment group that is in the active, tyrosine phosphorylated form (expressed as % pEphB2). This indicated without any copper, the cell stimulation with EphrinB2-Fc resulted in ~4% of the total EphB2 becoming the activated, pEphB2 form, and with 20 uM copper the % pEphB2 actually increased to ~5.5%, which then became reduced with higher concentrations of copper to only reach 1.5% in the presence of 160 uM copper. This biochemical data is consistent with the biphasic response to copper observed in BLI studies and indicate the metal at low concentrations can increase pEphB2 levels and cell signaling, while at high concentrations leads to reduced activation of the receptor.

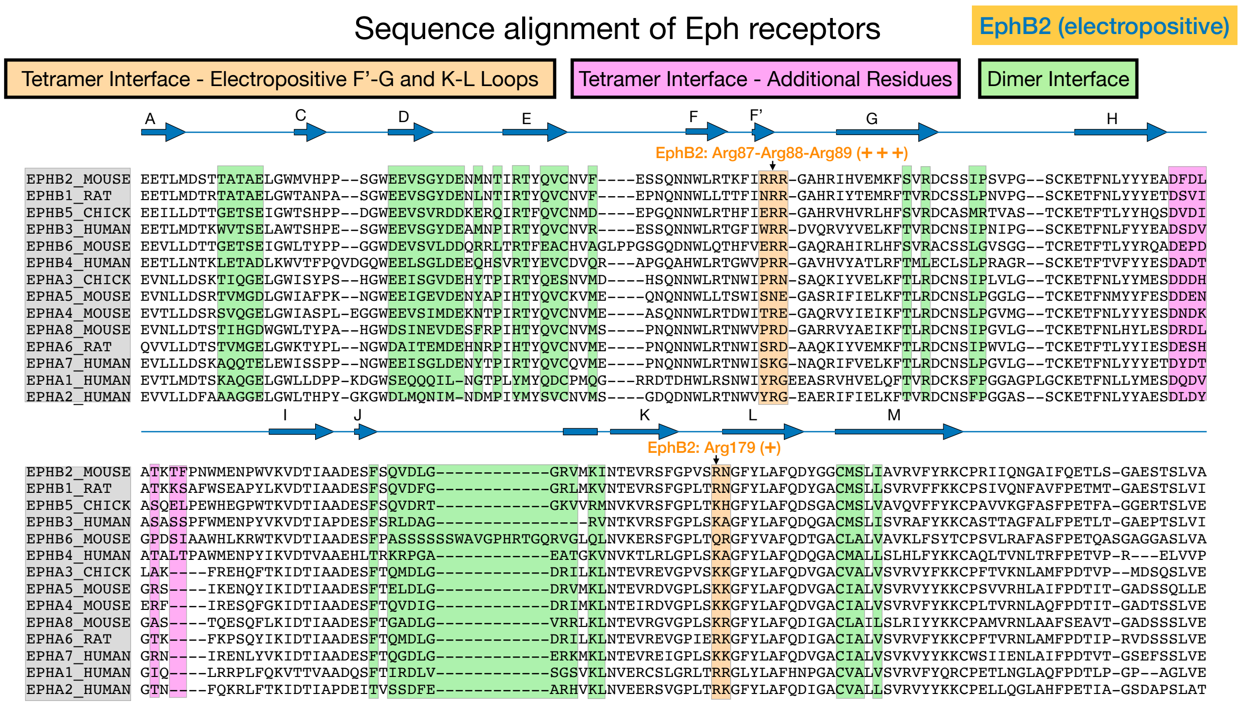


**Figure S2A.** Sequence alignment of the ligand-binding domains of Eph receptors. Based on the co-crystal structure of EphB2-EphrinB2, key residues in the tetramerization interface H-I loop that make contact with EphrinB2 are highlighted in magenta, whereas the key arginine-rich electropositive loops F'-G and K-L of the EphB2 tetramer interface that contain Arg87, Arg88, Arg89, Arg179, and Asn180 are highlighted in orange. Residues involved in the dimer interface are highlighted in green. Inspired from: Himanen, Rajashankar, Lackmann, Cowan, Henkemeyer, and Nikolov "Crystal structure of an Eph receptor-ephrin complex". Nature 414: 933–938 (2001).

Benchling [Sequence Alignment]. (2025). Retrieved from https://benchling.com

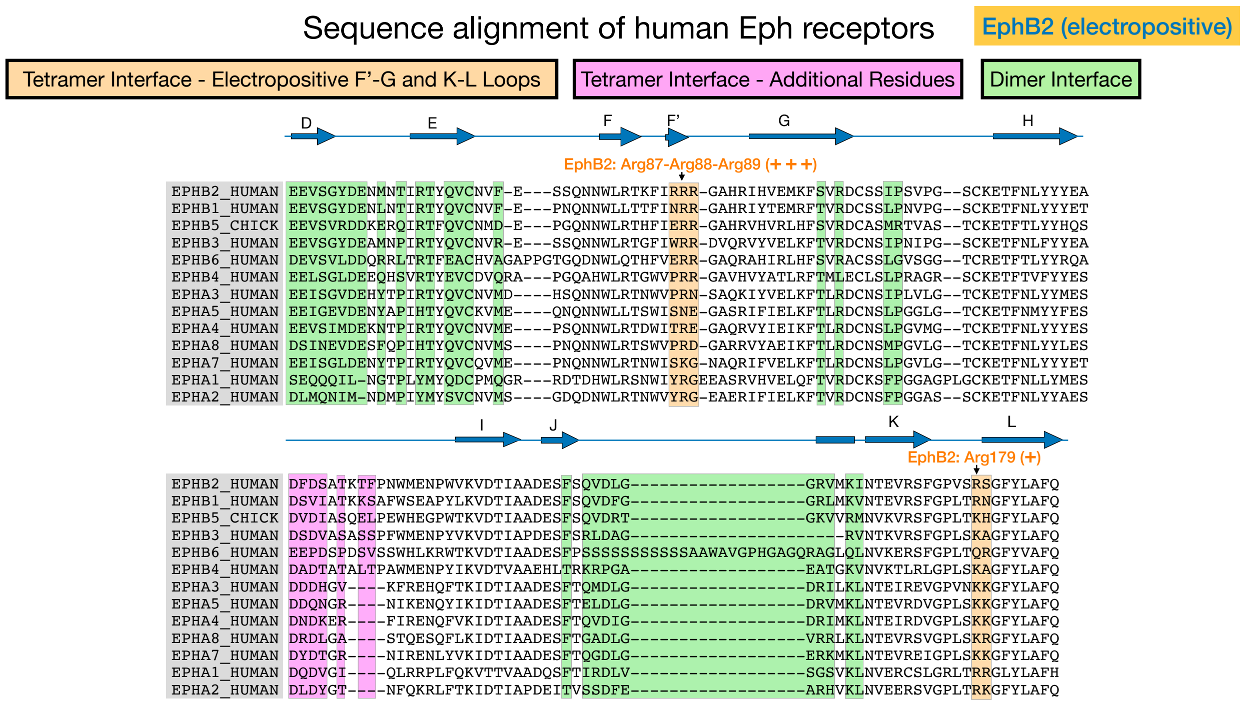


**Figure S2B.** Sequence alignment of the ligand-binding domains of human Eph receptors. Key arginine residues in EphB2 loops F'-G and K-L in the tetramerization interface are highlighted in orange, including Arg87, Arg88, Arg89, Arg179, and Asn180. Residues in the EphB2 tetramerization interface H-I loop that make contact with EphrinB2 are highlighted in magenta. Residues involved in the dimer interface are highlighted in green.

#### Sequence alignment of Ephrin ligands

EphrinB2
(copper-binding ancestry)

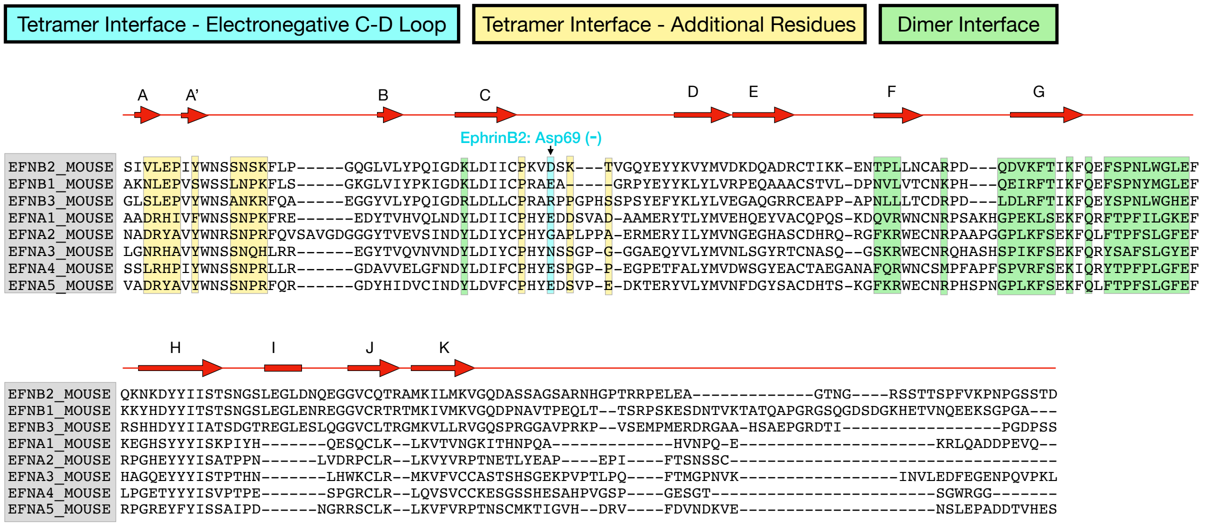


**Figure S2C.** Sequence alignment of the receptor-binding domains of Ephrin ligands. Based on the co-crystal structure of EphB2-EphrinB2, key residues in the tetramerization interface that make contact with EphB2 are highlighted yellow in loops A-A’, A’-B, and C-D, with the electronegative Asp69 of EphrinB2 highlighted in light blue. Residues involved in the dimer interface are highlighted in green. Inspired from: Himanen, Rajashankar, Lackmann, Cowan, Henkemeyer, and Nikolov “Crystal structure of an Eph receptor-ephrin complex”. Nature 414: 933–938 (2001).

List of proteins from PDB

| Eph-Ephrin Tetramer (receptor – ligand) | | Eph Monomer (receptor) | | Ephrin Monomer (ligand) | | Eph-Ephrin Dimer (receptor – ligand) | | Cupredoxin (copper binding proteins) | |
|---|---|---|---|---|---|---|---|---|---|
| mEphB2 – mEphrinB2 | 1KGY | mEphB2 | 1NUK | mEphrinB2 | 1IKO | hEphB4 – hEphrinB2 | 2HLE | Plastocyanin Cupredoxin from Cyanobacteria | 1JXD |
| hEphA4 – hEphrinB3 | 4BKF | hEphB3 | 3P1I | hEphrinA5 | 4ET7 | hEphA2 – hEphrinA1 | 3CZU | Cytochrome Cupredoxin from bacteria electron transport | 2CUA |
| | | hEphB6 | 7K7J | | | hEphA3 – hEphrinA5 | 4L0P | Red Copper Nitrosocynanin Cupredoxin from bacteria | 1IBZ |
| | | hEphA2 | 2X10 | | | hEphA4 – hEphrinA2 | 2WO3 | Azurin from Bacterial electron transport | 2CCW |
| | | hEphA4 | 4BK4 | | | | | | |
| | | hEphA7 | 3NRU | | | | | | |

| Other Proteins | |
|---|---|
| Henipavirus with mEphrinB1 | 6P7S |
| SNEW – hEphB2 | 2QBX |

**Figure S2D.** Ectodomain protein structures used for Molecular Operating Environment (MOE) simulations are from Protein Data Bank (PBD) and their PDB code are provided.

#### Similarity of Eph receptors

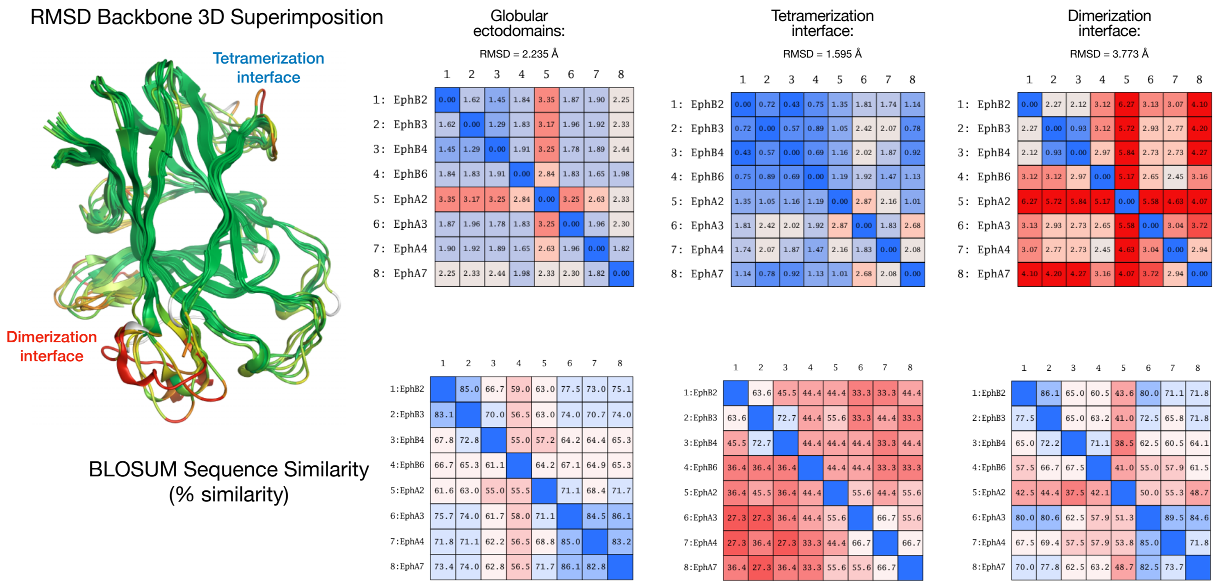


**Figure S2E.** Similarity of Eph receptor ectodomains determined by RMSD backbone superimposition test and BLOSUM sequence similarity test. RMSD superimposition results show similarity of the available Eph crystal structures throughout the entire globular ectodomains involved in Ephrin binding, with higher similarity in tetramerization interface (more blue colors), and lower similarity in dimerization interface (more red colors). BLOSUM % amino acid sequence similarity results show overall high conservation of the globular ectodomains, though the isolated tetramerization interface is less conserved compared to the dimerization interface (red colors). The data indicates amino acid backbones of proteins and sequence similarity of proteins do not necessarily correlate.

Mutation analysis for EphB2 tetramer affinity to EphrinB2

87 89 179
**EphB2: RRR - - - RN**
**EphB1: NRR - - - RN**
**EphB3: WRR - - - KA**
**EphB4: PRR - - - KA**
**EphA1: YRGE - - RR**
**EphA2: YRGE - - RK**
**EphA3: PRN - - - KK**

| Target protein | “mutation" | | | | "dAffinity" | "dStability" |
|---|---|---|---|---|---|---|
| EphB2 | R87 | R89 | R179 | N180 | 0 | 0 |
| EphB1 | N | - | - | - | 0.7283 | 1.2277 |
| EphB3 | W | - | K | A | -2.8704 | 0.7369 |
| EphB4 | P | - | K | A | 5.1244 | 1.6185 |
| EphA1 | Y | G | - | R | -7.4263 | -0.2102 |
| EphA2 | Y | G | - | K | -3.5095 | 1.1121 |
| EphA3 | P | N | K | K | 1.5344 | 1.8508 |

**Figure S2F.** The EphB2-EphrinB2 dimer (taken from 1KGY) was protonated and minimized in isolation using MOE. EphB2 was taken in isolation. Amino acids at positions 87, 89, 179, and 180 of EphB2 were changed as indicated *in silico* to swap EphB2 residues with the corresponding residues of other Eph receptors, which were then analyzed using MOE to predict the relative affinity to the tetramer interface of EphrinB2. The first row represents the control trial; MOE was accurately able to predict the alignment of Arg87 in the tetramer pocket as the score of “0” represents natural affinity. ‘dAffinity’ scores greater than 0 indicate a decreased affinity of the modified EphB2 protein to EphrinB2, whereas scores lower than 0 indicate increased affinity to EphrinB2. Further, ‘dStability’ estimates the thermodynamic stability of these complexes in PBS. The data indicates that proline at position 87 strongly disrupts tetramer affinity of mutated EphB2 to EphrinB2, and is consistent with BLI studies that show EphB4 with a proline at this position exhibits a more dimer-driven interaction with EphrinB2.

#### Mutation analysis for EphrinB2 tetramer affinity to EphB2

| Target protein | “mutation" | "dAffinity" | "dStability" |
|---|---|---|---|
| EphrinB2 | Asp69 | 0 | 0 |
| | Glu | 0.587797045749198 | 0.181852171471974 |
| | Ala | 1.04905630952123 | 0.669271828089222 |
| | Arg | 1.74057574390611 | 0.280617288442727 |
| | Lys | 1.13528991197961 | 1.92722750768508 |

**Figure S2G.** The EphB2-EphrinB2 dimer (taken from 1KGY) was protonated and minimized in isolation using MOE. EphrinB2 was taken in isolation. Aspartic acid at position 69 on EphrinB2 was mutated *in silico* and MOE used to estimate the relative affinity to the tetramer interface of EphB2. The first row represents the control trial; MOE was accurately able to predict the alignment of Asp69 in the tetramer pocket as the score of “0” represents natural affinity. ‘dAffinity’ scores greater than 0 indicate decreased affinity to EphB2. ‘dAffinity’ scores lower than 0 indicate increased affinity to EphB2. Further, ‘dStability’ estimates the thermodynamic stability of these complexes in PBS. The data indicates mutation of Asp69 to another electronegative amino acid, Glu69, has little affect on the ability of EphrinB2 to tetramerize with EphB2. Changing this residue to an uncharged or positive charged amino acid lowered the affinity.

### Codes used for MOE simulations

#### Interaction Table Code

D - sidechain hydrogen bond donor
A - sidechain hydrogen bond acceptor
d - backbone hydrogen bond donor
a - backbone hydrogen bond acceptor
O - solvent hydrogen bond
I - ionic attraction
M - metal ligation
R - arene attraction
H - hydrophobic surface contact
Q - charged surface contact
P - partial hydrophobic contact
X - other surface contact
C - total surface contact

#### 2D Interaction Map Code

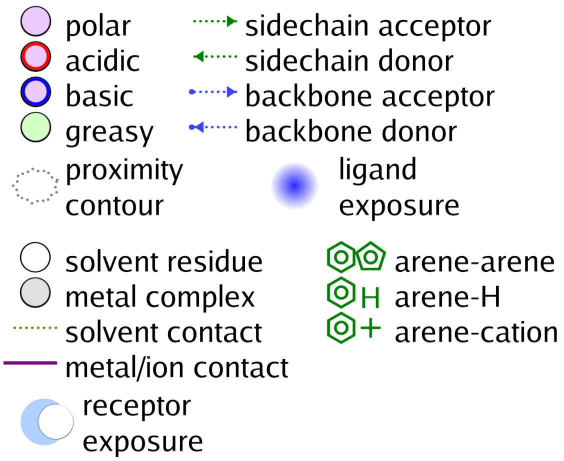


**Figure S2H.** Codes for interaction tables and 2D interaction maps resulting from MOE docking simulations. In the 2D interaction maps, maroon lines are ionic bonds, green arrows are sidechain hydrogen bonds, blue arrows are backbone hydrogen bonds, light purple are polar residues and light green are non-polar residues, with receptor exposure indicated by the blue off-set, and blue gradients indicate ligand exposure.

### MOE docking results for EGTA and EDTA binding to EphB2

Interaction Table for EGTA and EDTA docked to EphB2

| entry | molecule | Score | 28 | 29 | 30 | 31 | 32 | 33 | 36 | 84 | 85 | 87 | 89 | 90 | 125 | 127 | 129 | 180 | 202 |
|---|---|---|---|---|---|---|---|---|---|---|---|---|---|---|---|---|---|---|---|
| 1 | EGTA | -4.58868 | --a-- | d-aa-- | -- | -- | ---- | ---- | ------ | --- | -- | ------- | A---ll- | -- | ---- | ------ | ---- | - | A |
| 2 | EGTA | -4.30423 | ----- | ------ | -- | -- | a--- | ---- | ------ | --- | -- | ------- | AA--ll- | -- | ---- | ------ | ---- | - | - |
| 3 | EGTA | -3.99818 | ----- | ------ | -- | -- | ---- | ---- | ------ | --- | -- | -aal--- | AA--ll- | -- | ---- | ------ | ---- | - | - |
| 4 | EGTA | -3.98847 | ----- | ------ | -- | -- | a--- | ---- | ------ | --- | -- | ---l--- | AA--ll- | -- | ---- | ------ | ---- | - | - |
| 5 | EGTA | -3.91281 | ----- | ------ | d- | -- | aa-- | ---- | ------ | --- | -- | ------- | AA--ll- | -- | ---- | ------ | ---- | - | - |
| 1 | EDTA | -3.77528 | ---l- | --aa-- | -- | -- | ---- | ---- | ------ | --- | -- | ------- | AA--ll- | -- | ---- | ------ | ---- | - | A |
| 2 | EDTA | -3.62733 | ----- | d-aa-- | -- | -- | ---- | ---- | ------ | --- | -- | ------- | AA--ll- | -- | ---- | ------ | ---- | - | A |
| 3 | EDTA | -2.96329 | --a-- | --aa-- | -- | -- | ---- | ---- | ------ | --- | -- | ------- | AA--ll- | -- | ---- | ------ | ---- | - | - |
| 4 | EDTA | -2.94988 | ----- | ------ | -- | -- | ---- | ---- | ------ | A-- | -- | -a-ll-- | AA--ll- | -- | ---- | ------ | ---- | - | - |
| 5 | EDTA | -2.80558 | ----- | ------ | -- | -- | ---- | ---- | ------ | --- | -- | A--ll-- | ----l-- | -- | ---- | ------ | ---- | - | - |

**Figure S2H** (continued). Interaction Table for MOE analysis of EphB2 docking to EDTA and EGTA.

Interaction Table for 8-HQ and 3511 compounds with a hydrogen, bromine, or iodine at position 7 and docked to EphB2

| Rank | molecule | Score | 28 | 29 | 30 | 31 | 32 | 33 | 36 | 84 | 85 | 87 | 89 | 90 | 125 | 127 | 129 | 180 | 202 |
|---|---|---|---|---|---|---|---|---|---|---|---|---|---|---|---|---|---|---|---|
| 1 | 8-HQ series - Br at position 7 | -2.62076 | ----- | ------ | -- | RR | ---- | ---- | ------ | --- | -- | ------- | AA--II- | -- | ---- | ------ | ---- | - | - |
| 2 | 8-HQ series - Br at position 7 | -2.50643 | ----- | ------ | -- | -- | ---- | ---- | ------ | --- | -- | ------- | ------R | -- | ---- | ------ | ---- | - | - |
| 3 | 8-HQ series - Br at position 7 | -2.26498 | ----- | ------ | -- | -- | ---- | ---- | ------ | --- | -- | -----RR | ------- | -- | ---- | ------ | ---- | - | - |
| 4 | 8-HQ series - Br at position 7 | -2.2176 | ----- | ------ | -- | R- | --R- | ---- | ------ | --- | -- | ------- | AA--II- | -- | ---- | ------ | ---- | - | - |
| 5 | 8-HQ series - Br at position 7 | -2.02484 | ----- | ------ | -- | -- | ---- | ---- | D----- | --- | -- | ------- | AA--II- | -- | ---- | ------ | ---- | - | - |
| 1 | 8-HQ series - H at position 7 | -2.90725 | ----- | ------ | d- | -- | ---- | ---- | ------ | --- | -- | -----R- | ------- | -- | ---- | ------ | ---- | - | - |
| 2 | 8-HQ series - H at position 7 | -2.59065 | ----- | ------ | -- | -- | ---- | ---- | ------ | --- | -- | -----R- | AA----- | -- | ---- | ------ | ---- | - | - |
| 3 | 8-HQ series - H at position 7 | -1.97696 | ----- | ------ | -- | -- | ---- | ---- | ------ | --- | -- | ------- | ------- | -- | ---- | ------ | ---- | - | - |
| 1 | 8-HQ series - I at position 7 | -2.44338 | ----- | ------ | -- | RR | ---- | ---- | ------ | --- | -- | ------- | AA--II- | -- | ---- | ------ | ---- | - | - |
| 2 | 8-HQ series - I at position 7 | -2.28282 | ----- | ------ | -- | R- | ---- | ---- | ------ | --- | -- | ------- | AA--II- | -- | ---- | ------ | ---- | - | - |
| 3 | 8-HQ series - I at position 7 | -2.2432 | ----- | ------ | -- | -- | ---- | ---- | ------ | --- | -- | ------- | AA--II- | -- | ---- | ------ | ---- | - | - |
| 4 | 8-HQ series - I at position 7 | -2.05397 | ----- | ------ | -- | -- | ---- | ---- | ------ | --- | -- | ------- | AA--II- | -- | ---- | ------ | ---- | - | - |
| 5 | 8-HQ series - I at position 7 | -1.82613 | ----- | ------ | -- | RR | --RR | ---- | ------ | --- | -- | ------- | AA--II- | -- | ---- | ------ | ---- | - | - |
| 1 | 3511 series - Br at position 7 | -3.7544 | ----- | ------ | -- | RR | --R- | ---- | ------ | --- | -- | ------- | AA--II- | -- | ---- | ------ | ---- | - | - |
| 2 | 3511 series - Br at position 7 | -3.45038 | ----- | ------ | -- | RR | ---- | ---- | ------ | --R | -- | ------- | AA--II- | -- | ---- | ------ | ---- | - | - |
| 3 | 3511 series - Br at position 7 | -3.15217 | ----- | ------ | -- | -- | a--- | ---- | ------ | --- | -- | ------- | ------- | -- | ---- | ------ | ---- | - | - |
| 4 | 3511 series - Br at position 7 | -3.1107 | ----- | ------ | -- | RR | --R- | ---- | ------ | --- | -- | ------- | AA--II- | -- | ---- | ------ | ---- | - | - |
| 5 | 3511 series - Br at position 7 | -3.08317 | ----- | ------ | -- | RR | --R- | ---- | ------ | --- | -- | ------- | AA--II- | -- | ---- | ------ | ---- | - | - |
| 1 | 3511 series - H at position 7 | -3.97713 | ----- | ------ | -- | R- | ---- | ---- | ------ | --- | dd | ------- | ------- | -- | ---- | ------ | ---- | - | - |
| 2 | 3511 series - H at position 7 | -3.88795 | ----- | ------ | -- | -- | ---- | ---- | ------ | --- | -- | -----R- | AA----- | -- | ---- | ------ | ---- | - | - |
| 3 | 3511 series - H at position 7 | -3.28609 | ----- | ------ | -- | -- | ---- | ---- | ------ | --- | -- | ------- | ------- | -- | ---- | ------ | DD-- | - | - |
| 4 | 3511 series - H at position 7 | -3.23952 | ----- | ------ | -- | -- | ---- | ---- | ---d-- | --- | dd | -a----- | ------R | -- | ---- | ------ | ---- | - | - |
| 5 | 3511 series - H at position 7 | -2.93015 | ----- | ------ | dd | -- | ---- | ---- | ------ | --- | -- | -----R- | A------ | -- | ---- | ------ | ---- | - | - |
| 1 | 3511 series - I at position 7 | -2.97039 | ----- | ------ | -- | -- | ---- | ---- | --A--- | --- | -- | ------- | ------- | -- | ---- | ------ | ---- | - | - |
| 2 | 3511 series - I at position 7 | -2.59611 | ----- | ------ | -- | -- | ---- | ---- | ------ | --- | -- | ------- | AA--II- | -- | ---- | ------ | ---- | - | - |
| 3 | 3511 series - I at position 7 | -2.53509 | ----- | ------ | -- | -- | ---- | ---- | ------ | --- | -- | ------- | AA--II- | -- | ---- | ------ | ---- | - | - |
| 4 | 3511 series - I at position 7 | -2.2718 | ----- | ------ | -- | -- | ---- | ---- | ------ | --- | -- | ------- | AA--II- | -- | ---- | ------ | ---- | - | - |
| 5 | 3511 series - I at position 7 | -2.24569 | ----- | ------ | -- | -- | ---- | ---- | ------ | --- | -- | -aa---- | ------- | -- | ---- | ------ | ---- | - | - |

**Figure S2H** (continued). Interaction Table for MOE analysis of EphB2 docking to 8-HQ and 3511 compounds with a hydrogen, bromine, or iodine at position 7.

Interaction Table for A20 and BQPB4 compounds with a hydrogen, bromine, or iodine at position 7 and docked to EphB2

| Rank | molecule | Score | 28 | 29 | 30 | 31 | 32 | 33 | 36 | 84 | 85 | 87 | 89 | 90 | 125 | 127 | 129 | 180 | 202 |
|---|---|---|---|---|---|---|---|---|---|---|---|---|---|---|---|---|---|---|---|
| 1 | A20 series - Br at position 7 | -2.7653 | ----- | ------ | -- | -- | ---- | ---- | ------ | --- | -- | -----R- | AA--IIR | -- | ---- | ------ | ---- | - | - |
| 2 | A20 series - Br at position 7 | -2.10786 | ----- | ------ | -- | -- | ---- | ---- | ------ | --- | dd | ------- | AA--II- | -- | ---- | ------ | ---- | - | - |
| 3 | A20 series - Br at position 7 | -2.10384 | DD-II | ----R- | -- | -- | ---- | ---- | ------ | --- | -- | ------- | AA--II- | -- | ---- | ------ | ---- | - | - |
| 4 | A20 series - Br at position 7 | -2.05846 | ----- | ------ | -- | -- | ---- | DDII | ------ | --- | -- | ------- | AA--II- | -- | ---- | ------ | ---- | - | - |
| 5 | A20 series - Br at position 7 | -2.00155 | ----- | ------ | -- | -- | ---- | ---- | ------ | --- | -- | ------- | AA--II- | -- | ---- | ------ | ---- | - | - |
| 1 | A20 series - H at position 7 | -3.63755 | ----- | ------ | -- | -- | ---- | ---- | ------ | --- | -- | -----R- | ------- | -- | ---- | ------ | ---- | - | - |
| 2 | A20 series - H at position 7 | -3.15724 | ----- | ------ | dd | -- | ---- | ---- | ------ | AA- | -- | ------- | AA----- | -- | ---- | ------ | ---- | - | - |
| 3 | A20 series - H at position 7 | -3.13758 | ----- | ------ | dd | -- | ---- | ---- | ------ | --- | -- | ------- | A------ | -- | ---- | ------ | ---- | - | - |
| 4 | A20 series - H at position 7 | -2.91859 | ----- | ------ | -- | -- | ---- | ---- | ------ | --- | -- | -----R- | ------- | -- | ---- | ------ | ---- | - | - |
| 5 | A20 series - H at position 7 | -2.91239 | ----- | ------ | -- | -- | ---- | ---- | ------ | --- | -- | ------- | A-----R | -- | ---- | ------ | ---- | - | - |
| 1 | A20 series - I at position 7 | -2.88006 | ----- | ------ | -- | RR | ---- | ---- | ---dd- | --- | -- | ------- | AA--II- | -- | ---- | ------ | ---- | - | - |
| 2 | A20 series - I at position 7 | -2.77622 | ----- | ------ | -- | -- | ---- | ---- | ------ | --- | -- | ------- | AA--IIR | -- | ---- | ------ | ---- | - | - |
| 3 | A20 series - I at position 7 | -2.24822 | DD-II | ----RR | -- | -- | ---- | ---- | ------ | --- | -- | ------- | AA--II- | -- | ---- | ------ | ---- | - | - |
| 4 | A20 series - I at position 7 | -2.11957 | ----- | ------ | -- | -- | ---- | ---- | ------ | --- | -- | -----R- | AA--II- | -- | ---- | ------ | ---- | - | - |
| 5 | A20 series - I at position 7 | -2.07171 | D--I- | ----R- | -- | -- | ---- | ---- | ------ | --- | -- | ------- | AA--II- | -- | ---- | ------ | ---- | - | - |
| 1 | BQPB4 series - Br at position 7 | -2.83837 | ----- | ------ | -- | -- | --RR | ---- | ------ | --- | -- | ------- | AA--II- | -- | ---- | ------ | ---- | - | - |
| 2 | BQPB4 series - Br at position 7 | -2.42857 | ----- | ------ | -- | -- | ---- | ---- | ------ | --- | -- | ------- | AA--II- | -- | ---- | ------ | ---- | - | - |
| 3 | BQPB4 series - Br at position 7 | -2.40508 | ----- | ------ | -- | -- | --R- | ---- | ------ | --- | -- | ---I--- | ------- | -- | ---- | ------ | ---- | - | - |
| 4 | BQPB4 series - Br at position 7 | -2.36113 | ----- | ------ | -- | -- | ---- | ---- | -----R | --- | -- | -----R- | AA--II- | -- | ---- | ------ | ---- | - | - |
| 5 | BQPB4 series - Br at position 7 | -2.5632 | ----- | ------ | -- | -- | ---- | ---- | -----R | --- | -- | -aa---- | ------R | -- | ---- | ------ | ---- | - | - |
| 1 | BQPB4 series - H at position 7 | -4.51074 | ----- | ------ | -- | -- | ---- | ---- | -----R | --- | -- | ------- | ------- | -- | ---- | ------ | ---- | - | - |
| 2 | BQPB4 series - H at position 7 | -3.49637 | ----- | ------ | -- | -- | ---- | ---- | ------ | --- | -- | -----R- | AA----- | -- | ---- | ------ | ---- | - | - |
| 3 | BQPB4 series - H at position 7 | -3.16189 | ----- | ------ | -- | -- | --RR | ---- | ------ | --- | dd | -----R- | ------- | -- | ---- | ------ | ---- | - | - |
| 1 | BQPB4 series - I at position 7 | -2.46621 | ----- | ------ | -- | -- | ---- | ---- | ------ | --- | -- | ------- | AA--IIR | -- | ---- | ------ | ---- | - | - |
| 2 | BQPB4 series - I at position 7 | -2.41905 | ----- | ------ | -- | -- | ---- | ---- | ------ | --- | -- | -----R- | AA--II- | -- | ---- | ------ | ---- | - | - |
| 3 | BQPB4 series - I at position 7 | -2.28499 | ----- | ------ | -- | -- | ---- | ---- | -----R | --- | -- | A------ | AA--II- | -- | ---- | ------ | ---- | - | - |
| 4 | BQPB4 series - I at position 7 | -2.18215 | ----- | ------ | -- | -- | ---- | ---- | ------ | --- | -- | -aa---- | ------- | -- | ---- | ------ | ---- | - | - |
| 5 | BQPB4 series - I at position 7 | -2.11262 | ----- | ------ | -- | -- | ---- | ---- | ---dd- | --R | -- | ---I--- | ------- | -- | ---- | ------ | ---- | - | - |

**Figure S2H** (continued). Interaction Table for MOE analysis of EphB2 docking to A20 and BQPB4 compounds with a hydrogen, bromine, or iodine at position 7.

Interaction Table for G15 and primary amine compounds with a hydrogen, bromine, or iodine at position 7 and docked to

| entry | molecule | Score | 28 | 29 | 30 | 31 | 32 | 33 | 36 | 84 | 85 | 87 | 89 | 90 | 125 | 127 | 129 | 180 | 202 |
|---|---|---|---|---|---|---|---|---|---|---|---|---|---|---|---|---|---|---|---|
| 1 | G15 series - Br at position 7 | -3.65549 | ----- | ------ | -- | -- | ---- | ---- | ------ | --- | -- | -aa---- | ------- | -- | ---- | ------ | ---- | - | - |
| 2 | G15 series - Br at position 7 | -3.33544 | ----- | ------ | -- | -- | ---- | ---- | ------ | --- | d- | ------- | AA--II- | -- | ---- | ------ | ---- | - | - |
| 3 | G15 series - Br at position 7 | -2.63673 | ----- | ------ | -- | -- | ---- | ---- | ------ | --- | -- | -----R- | AA--II- | -- | ---- | ------ | ---- | - | - |
| 4 | G15 series - Br at position 7 | -2.63267 | ----- | ------ | -- | -- | ---- | ---- | ------ | --- | -- | -----R- | AA--II- | -- | ---- | ------ | ---- | - | - |
| 1 | G15 series - H at position 7 | -4.21624 | ----- | ------ | dd | -- | ---- | ---- | ------ | --- | -- | ------- | A------ | -- | ---- | ------ | ---- | - | - |
| 2 | G15 series - H at position 7 | -4.15363 | ----- | ------ | -- | -- | ---- | ---- | ------ | --- | -- | -----R- | ------- | -- | ---- | ------ | ---- | - | - |
| 3 | G15 series - H at position 7 | -3.90407 | ----- | ------ | d- | -- | ---- | ---- | ------ | --R | -- | ------- | AA----- | -- | ---- | ------ | ---- | - | - |
| 4 | G15 series - H at position 7 | -3.88957 | ----- | ------ | dd | -- | ---- | ---- | ------ | --- | -- | -----R- | A------ | -- | ---- | ------ | ---- | - | - |
| 1 | G15 series - I at position 7 | -3.42171 | ----- | ------ | -- | -- | ---- | ---- | ------ | --- | -- | ------- | AA--IIR | -- | ---- | ------ | ---- | - | - |
| 2 | G15 series - I at position 7 | -3.09756 | ----- | ------ | -- | -- | ---- | ---- | ------ | --- | -- | -----R- | AA--II- | -- | ---- | ------ | ---- | - | - |
| 3 | G15 series - I at position 7 | -2.9022 | ----- | ------ | -- | -- | ---- | ---- | ------ | --- | -- | ------- | AA--IIR | -- | ---- | ------ | ---- | - | - |
| 4 | G15 series - I at position 7 | -2.70123 | ----- | ------ | -- | -- | ---- | ---- | ------ | --- | dd | ------- | AA--II- | -- | ---- | ------ | ---- | - | - |
| 5 | G15 series - I at position 7 | -2.54628 | ----- | ------ | -- | -- | ---- | ---- | ------ | --- | -- | ------- | ------R | -- | ---- | ------ | ---- | - | - |
| 1 | Primary amine - Br at position 7 | -3.10422 | ----- | ------ | -- | -- | ---- | DDI- | DD---- | --- | d- | ------- | AA--II- | -- | ---- | ------ | ---- | - | - |
| 2 | Primary amine - Br at position 7 | -3.04672 | ----- | ------ | -- | -- | ---- | DDII | DD---- | --- | -- | ------- | AA--II- | -- | ---- | ------ | ---- | - | - |
| 3 | Primary amine - Br at position 7 | -2.25096 | ----- | ------ | -- | -- | ---- | --I- | ------ | --- | -- | ------- | AA--II- | -- | ---- | ------ | ---- | - | - |
| 4 | Primary amine - Br at position 7 | -2.15608 | ----- | ------ | -- | -- | ---- | DDII | ------ | --- | -- | -aa---- | ------- | -- | ---- | ------ | ---- | - | - |
| 5 | Primary amine - Br at position 7 | -2.11851 | ----- | ------ | -- | -- | ---- | DDII | ------ | --- | -- | ------- | AA--II- | -- | ---- | ------ | ---- | - | - |
| 1 | Primary amine - I at position 7 | -2.94396 | ----- | ------ | -- | -- | ---- | --I- | DD---- | --- | -- | ------- | AA--II- | -- | ---- | ------ | ---- | - | - |
| 2 | Primary amine - I at position 7 | -2.77786 | ----- | ----R- | -- | -- | ---- | DDII | ------ | --- | -- | ------- | AA--II- | -- | ---- | ------ | ---- | - | - |
| 3 | Primary amine - I at position 7 | -2.76544 | ----- | ------ | -- | -- | ---- | DDII | DD---- | --- | -- | ------- | AA--II- | -- | ---- | ------ | ---- | - | - |
| 4 | Primary amine - I at position 7 | -2.21819 | ----- | ------ | -- | -- | ---- | --I- | ------ | --- | -- | ------- | ----IIR | -- | ---- | ------ | ---- | - | - |
| 5 | Primary amine - I at position 7 | -2.20649 | ----- | ------ | -- | -- | ---- | D-I- | ------ | --- | -- | -aa---- | ------- | -- | ---- | ------ | ---- | - | - |
| 1 | Primary amine | -3.66246 | ----- | ------ | dd | -- | a--- | ---- | ------ | --- | d- | ------- | ------- | -- | ---- | ------ | ---- | - | - |
| 2 | Primary amine | -3.30957 | ----- | ------ | dd | -- | ---- | ---- | ------ | --- | -- | -----R- | A------ | -- | ---- | ------ | ---- | - | - |
| 3 | Primary amine | -2.78084 | ----- | ------ | dd | -- | ---- | ---- | ------ | --- | -- | ------- | A------ | -- | ---- | ------ | DDII | - | - |
| 4 | Primary amine | -2.65443 | ----- | ------ | d- | -- | ---- | --I- | ------ | --- | -- | ------- | ------- | -- | ---- | ------ | ---- | - | - |

**Figure S2H** (continued). Interaction Table for MOE analysis of EphB2 docking to G15 and primary amine compounds with a hydrogen, bromine, or iodine at position 7.

MOE docking of compounds into the EphB2 tetramer interface

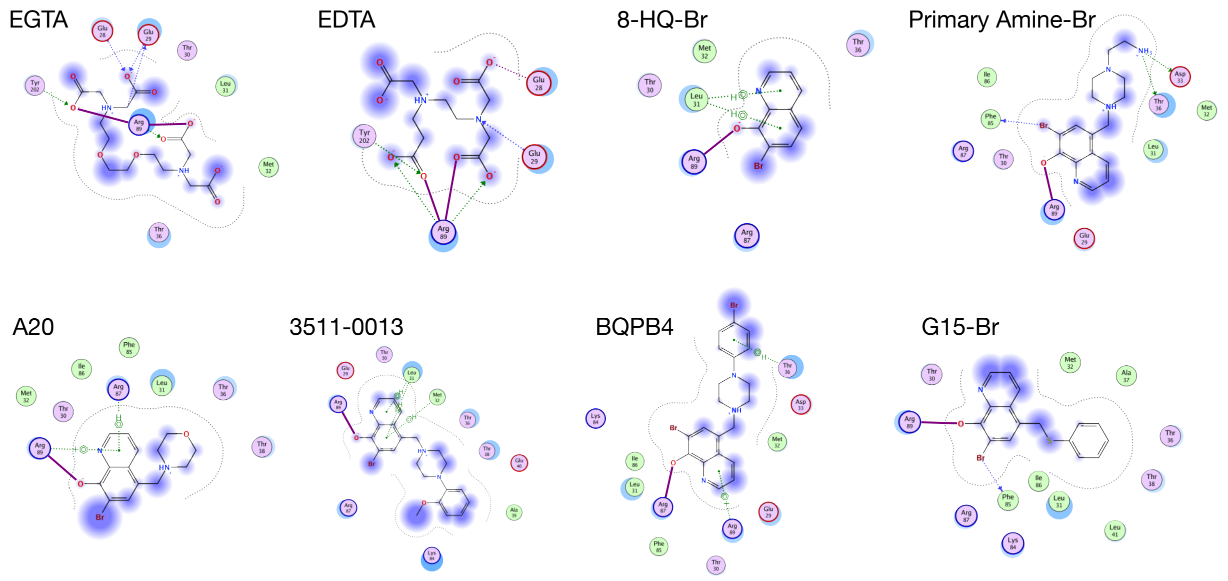


**Figure S2H** (continued). 2D interaction maps for MOE analysis of EphB2 docking to indicated compounds. Maroon lines are ionic bonds, green arrows are sidechain hydrogen bonds, blue arrows are backbone hydrogen bonds, light purple are polar residues and light green are non-polar residues, with receptor exposure indicated by the blue off-set, and blue gradients indicate ligand exposure.

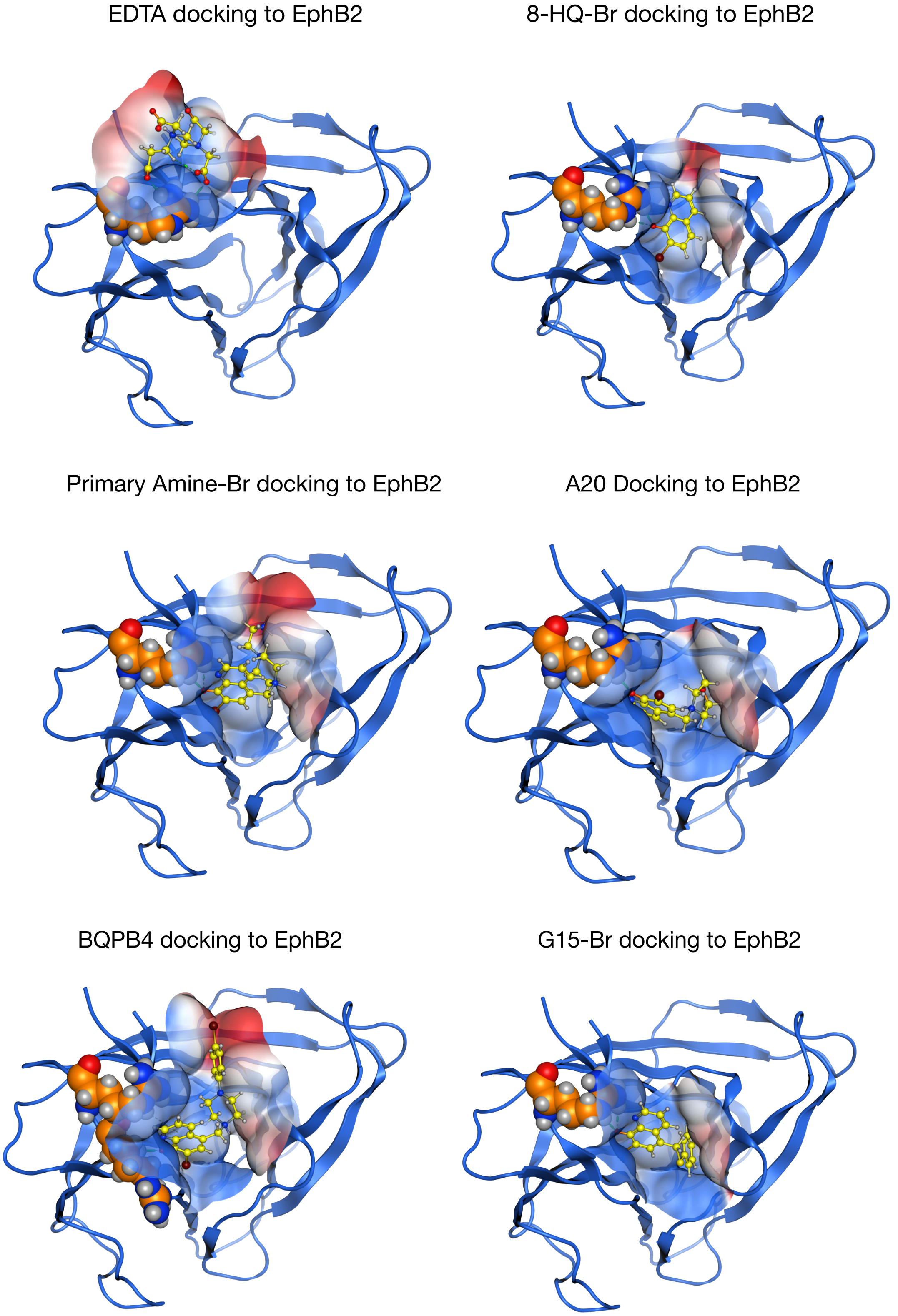


**Figure S2I.** MOE simulations show indicated compounds docking into the EphB2 tetramer interface via interactions with Arg89, which is conserved in all EphB receptors. Within the docking region of EphB2, the space-filling blue highlighted zones are electropositive areas and red are electronegative areas. The highest ranked docking configuration determined by the GBVI/WSA forcefield energy calculation is shown.

NAMD simulations (30 ns) - PBSA binding free energy (ΔG)

EGTA docking to EphB2 | 3511-0013 docking to EphB2

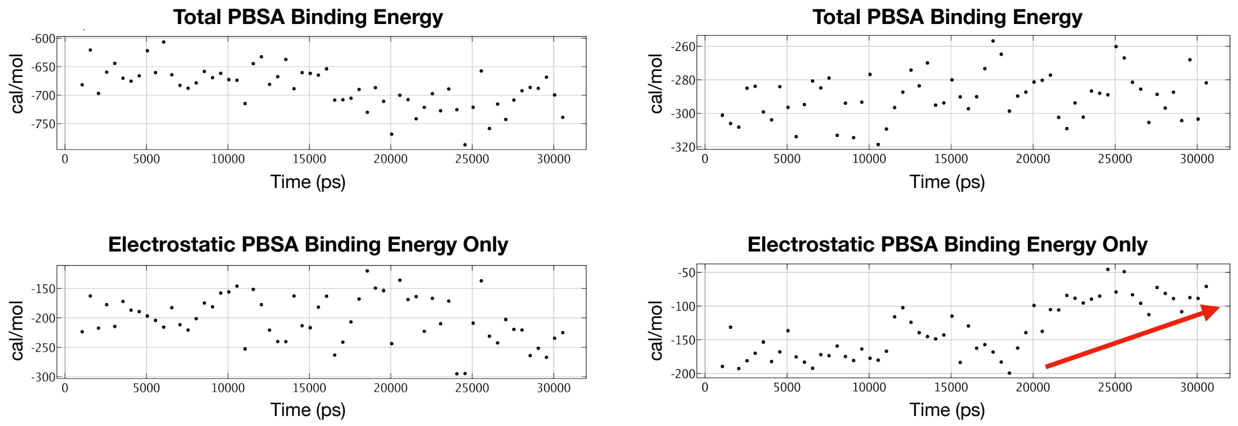


**Figure S2J.** Total and electrostatic PBSA free energy (ΔG) was calculated from 30 ns NMDA simulations of EGTA and 3511-0013 docking to EphB2. Total binding energy shows the multidentate chelator EGTA exhibits a higher overall affinity for EphB2 throughout the 30 ns simulation compared to the bidentate chelator 3511-0013. In the simulations, both EGTA and 3511-0013 exhibited strong electrostatic interactions with EphB2, though 3511-0013 tended to lose binding energy starting at around 20 ns which is is consistent with the observed dissociation of this compound from the receptor at this time (see Videos S12 and S13).

NAMD simulations (30 ns) - PBSA binding free energy (ΔG)

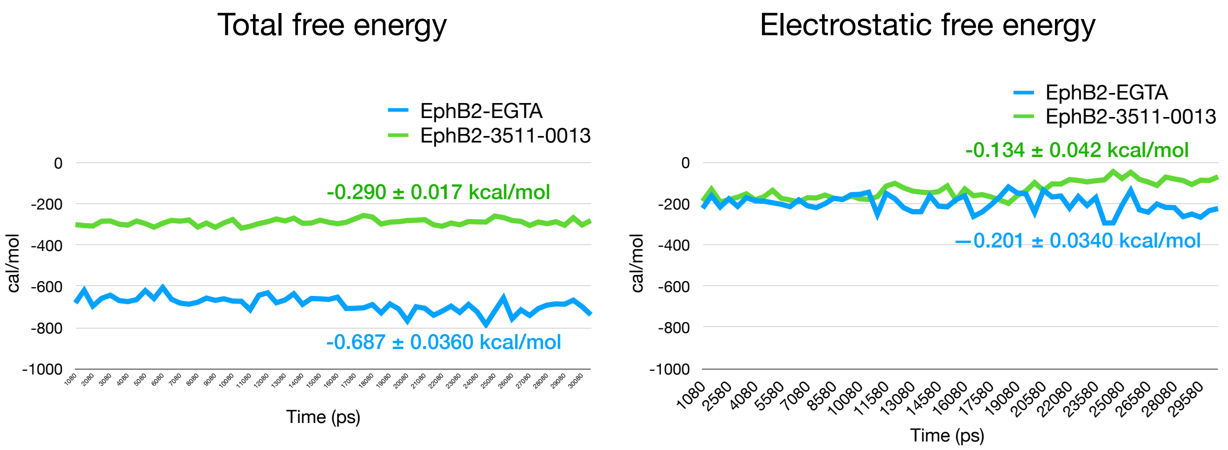


**Figure S2J** (continued). Compared to 3511-0013, EDTA docking to EphB2 is more energetically favorable throughout the 30 ns simulation. Note the divergence in electrostatic free energy starting approximately 20 ns into the simulation when 3511-0013 dissociates from the EphB2 receptor protein (see Videos S12 and S13).

NAMD simulations (30 ns) - Distance apart

EGTA docking to EphB2 | 3511-0013 docking to EphB2

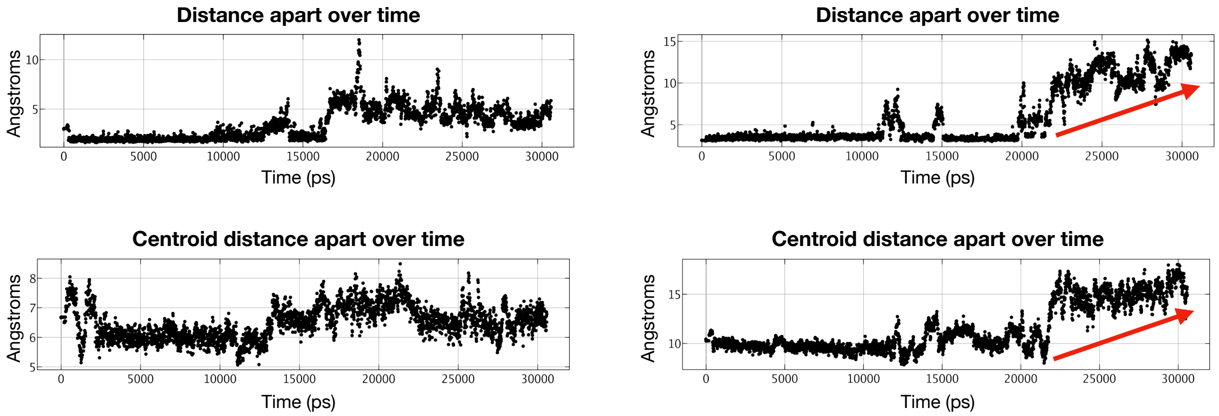


**Figure S2K.** Distance apart from EGTA or 3511-0013 and the Arg87 guanidine group of EphB2 was calculated from the 30 ns NAMD simulations of the two compounds docking to the receptor protein. Distance apart from the centroid of EGTA or 3511-0013 and the centroid of EphB2 was also calculated. In both measurements, EGTA binding to EphB2 is very stable and remained associated with the receptor for the entire 30 nanoseconds, though starting at around 17 ns the distance became variably increased, consistent with the observed partial dissociation and re-association of this compound from the receptor during the later half of the simulation (see Video S12). 3511-0013 also tightly bound to EphB2, though the interaction is less stable than EGTA as this compound showed partial dissociation and re-association events at ~11 and ~15 ns, and completely dissociated from the receptor protein after ~22 ns (red arrows) (see Video S13).

NAMD simulations (30 ns) - Snapshot of docking

EGTA docking to EphB2 | 3511-0013 docking to EphB2

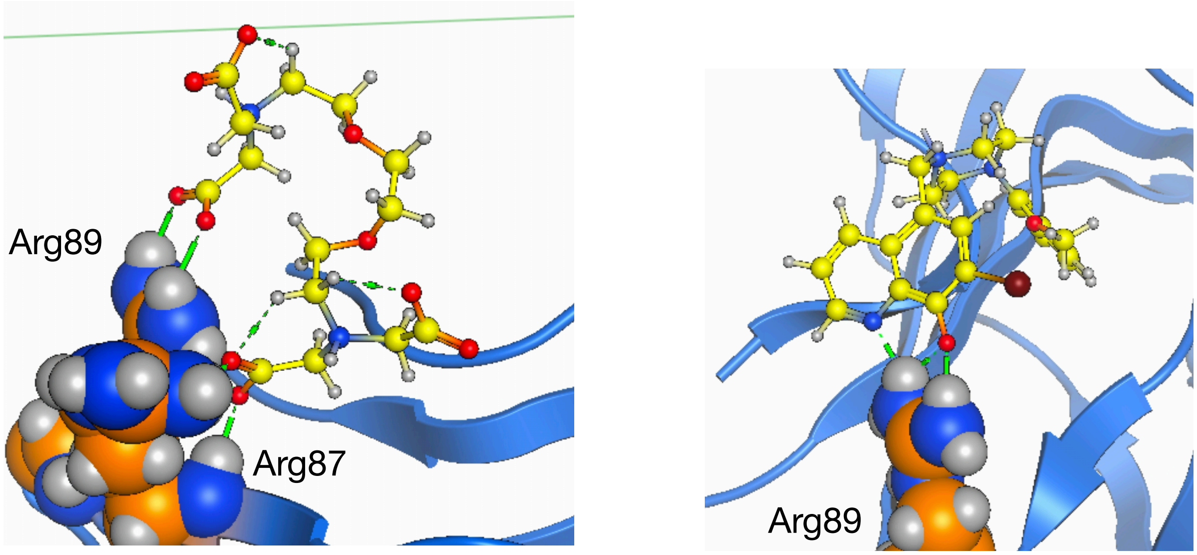


**Figure S2L.** Snapshot of 30 ns NMDA simulations of EGTA and 3511-0013 docking to EphB2 from Videos S12 and S13. The multidentate chelator EGTA interacts using its negative charged oxygens to cage both Arg89 and Arg87 of the EphB2 tetramer interface. The bidentate chelator 3511-0013 interacts with only Arg89 of EphB2, and involves both the oxygen (red) at position 8 and nitrogen (blue) at position 1 of the 8-hydroxyquinoline ring, consistent with the roles for these atoms in caging a metal ion. Electrostatic interactions are indicated in green.

#### Similarity of Ephrin ligands

##### RMSD Backbone Superimposition

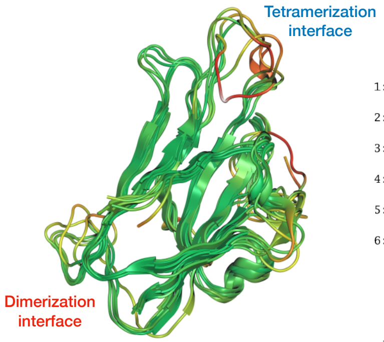


**Globular ectodomains** — RMSD = 2.618 Å

| | 1 | 2 | 3 | 4 | 5 | 6 |
|---|---|---|---|---|---|---|
| 1: EphrinB1 | 0.00 | 1.92 | 2.14 | 3.17 | 2.54 | 2.93 |
| 2: EphrinB2 | 1.92 | 0.00 | 2.34 | 3.02 | 2.32 | 2.59 |
| 3: EphrinB3 | 2.14 | 2.34 | 0.00 | 3.39 | 3.01 | 3.25 |
| 4: EphrinA1 | 3.17 | 3.02 | 3.39 | 0.00 | 2.24 | 2.02 |
| 5: EphrinA2 | 2.54 | 2.32 | 3.01 | 2.24 | 0.00 | 1.61 |
| 6: EphrinA5 | 2.93 | 2.59 | 3.25 | 2.02 | 1.61 | 0.00 |

**C-D Loop (Tetramer)** — RMSD = 3.034 Å

| | 1 | 2 | 3 | 4 | 5 | 6 |
|---|---|---|---|---|---|---|
| 1: EphrinB1 | 0.00 | 1.37 | 2.71 | 3.72 | 3.29 | 3.48 |
| 2: EphrinB2 | 1.37 | 0.00 | 2.06 | 3.77 | 3.43 | 3.61 |
| 3: EphrinB3 | 2.71 | 2.06 | 0.00 | 3.98 | 3.57 | 3.85 |
| 4: EphrinA1 | 3.72 | 3.77 | 3.98 | 0.00 | 1.65 | 1.01 |
| 5: EphrinA5 | 3.29 | 3.43 | 3.57 | 1.65 | 0.00 | 1.28 |
| 6: EphrinA2 | 3.48 | 3.61 | 3.85 | 1.01 | 1.28 | 0.00 |

**F-G Loop (Dimer)** — RMSD = 2.813 Å

| | 1 | 2 | 3 | 4 | 5 | 6 |
|---|---|---|---|---|---|---|
| 1: EphrinB1 | 0.00 | 4.14 | 2.33 | 3.19 | 3.67 | 3.19 |
| 2: EphrinB2 | 4.14 | 0.00 | 3.92 | 2.11 | 1.17 | 1.77 |
| 3: EphrinB3 | 2.33 | 3.92 | 0.00 | 3.42 | 3.54 | 3.26 |
| 4: EphrinA1 | 3.19 | 2.11 | 3.42 | 0.00 | 1.29 | 0.73 |
| 5: EphrinA5 | 3.67 | 1.17 | 3.54 | 1.29 | 0.00 | 0.98 |
| 6: EphrinA2 | 3.19 | 1.77 | 3.26 | 0.73 | 0.98 | 0.00 |

##### BLOSUM Sequence Similarity (% similarity)

**Globular ectodomains**

| | 1 | 2 | 3 | 4 | 5 | 6 |
|---|---|---|---|---|---|---|
| 1:EphrinB1 | | 77.4 | 72.1 | 48.5 | 47.7 | 46.2 |
| 2:EphrinB2 | 78.5 | | 72.9 | 47.7 | 48.5 | 47.0 |
| 3:EphrinB3 | 74.8 | 74.5 | | 50.0 | 45.4 | 47.0 |
| 4:EphrinA1 | 46.7 | 45.3 | 46.4 | | 68.5 | 62.9 |
| 5:EphrinA5 | 45.9 | 46.0 | 42.1 | 68.5 | | 79.5 |
| 6:EphrinA2 | 45.2 | 45.3 | 44.3 | 63.8 | 80.8 | |

**C-D Loop (Tetramer)**

| | 1 | 2 | 3 | 4 | 5 | 6 |
|---|---|---|---|---|---|---|
| 1:EphrinB1 | | 72.7 | 68.2 | 45.5 | 42.9 | 45.5 |
| 2:EphrinB2 | 80.0 | | 68.2 | 45.5 | 47.6 | 50.0 |
| 3:EphrinB3 | 75.0 | 68.2 | | 50.0 | 38.1 | 45.5 |
| 4:EphrinA1 | 50.0 | 45.5 | 50.0 | | 76.2 | 59.1 |
| 5:EphrinA5 | 45.0 | 45.5 | 36.4 | 72.7 | | 72.7 |
| 6:EphrinA2 | 50.0 | 50.0 | 45.5 | 59.1 | 76.2 | |

**F-G Loop (Dimer)**

| | 1 | 2 | 3 | 4 | 5 | 6 |
|---|---|---|---|---|---|---|
| 1:EphrinB1 | | 87.5 | 81.2 | 68.8 | 68.8 | 68.8 |
| 2:EphrinB2 | 87.5 | | 93.8 | 62.5 | 62.5 | 62.5 |
| 3:EphrinB3 | 81.2 | 93.8 | | 62.5 | 62.5 | 62.5 |
| 4:EphrinA1 | 68.8 | 62.5 | 62.5 | | 87.5 | 87.5 |
| 5:EphrinA5 | 68.8 | 62.5 | 62.5 | 87.5 | | 100.0 |
| 6:EphrinA2 | 68.8 | 62.5 | 62.5 | 87.5 | 100.0 | |

**Figure S3A.** Similarity of Ephrin ligand ectodomains determined by RMSD backbone superimposition test and BLOSUM sequence similarity test. Both indicated a clear separation between EphrinA and EphrinB class ligand ectodomains involved in Eph binding. For EphrinA ligands, RMSD superimposition and BLOSUM sequence similarity results show high similarity in both the tetramerization and dimerization interfaces. For EphrinB ligands, RMSD superimposition results show higher similarity in the 3D structure of the tetramerization interface than in the dimerization interface, indicating divergence in the three different EphrinB dimer loops, while BLOSUM sequence similarity results show lower similarity in tetramerization interface than in dimerization interface. The data indicates amino acid backbones of proteins and sequence similarity of proteins do not necessarily correlate.

### Copper metal binding sites on EphrinB2

EphrinB2 Amino Acid Interactions with Copper in Binding Site

| Type | ChainA | SetA | ChainB | Energy | Dist | BB | AtomsA | AtomsB |
|---|---|---|---|---|---|---|---|---|
| Ionic | EphrinB2 | Asp69 | Copper | -16.98 | 1.96 | - | OD1 | CU |
| Ionic | EphrinB2 | Asp69 | Copper | -16.86 | 1.97 | - | OD2 | CU |
| Metal | EphrinB2 | Thr72 | Copper | -3.78 | 1.99 | b | O | CU |
| Metal | EphrinB2 | Asp69 | Copper | -3.06 | 1.96 | - | OD1 | CU |
| Metal | EphrinB2 | Asp69 | Copper | -2.91 | 1.97 | - | OD2 | CU |
| Metal | EphrinB2 | Asp69 | Copper | -2.79 | 2.22 | b | O | CU |
| Metal | EphrinB2 | Lys71 | Copper | -2.77 | 2.03 | b | O | CU |
| Metal | EphrinB2 | Thr72 | Copper | -2.46 | 2.07 | - | OG1 | CU |

**Figure S3B.** MOE identification of specific amino acid residue interactions with copper are listed for predicted copper binding site. BB denotes whether any of the EphrinB2 interacting atoms in the entry are backbone (b) or not (-). AtomsA represents the atoms in residue A which are involved in the contact (when displaying individual contacts without aggregation). AtomsB represents the atoms in residue B which are involved in the contact (when displaying individual contacts without aggregation).

#### NAMD simulations (30 ns) - Copper metal binding site on EphrinB2

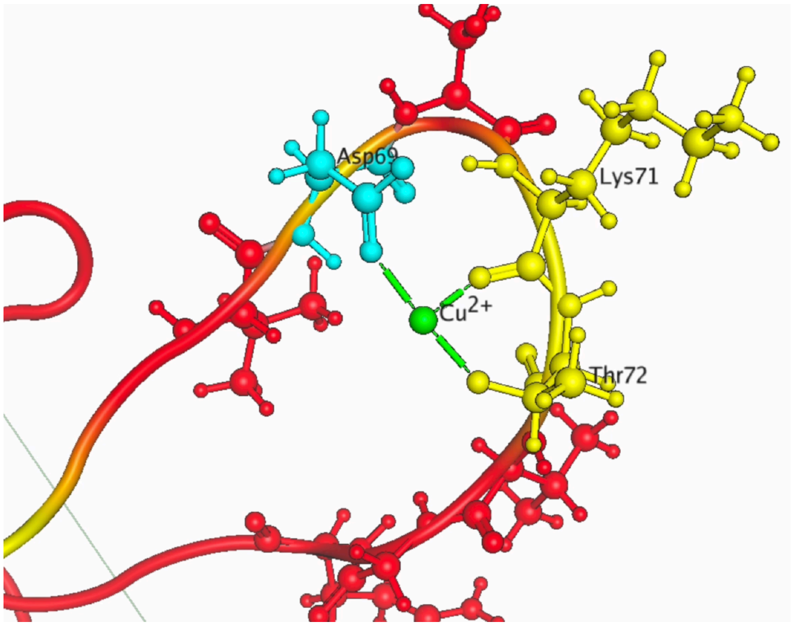


**Figure S3C.** Snapshot of 30 ns NMDA simulation of $Cu^{+2}$ to EphrinB2. This identified a strong potential interaction of copper with the C-D loop of the EphrinB2 tetramerization interface, particularly involving an ionic interaction between $Cu^{2+}$ and Asp69, as well as coordinate covalent bonds between $Cu^{2+}$ and both Lys71 and Thr72. Electrostatic and coordinate covalent interactions are indicated in green.

NAMD simulations (30 ns) - PBSA binding free energy (ΔG)

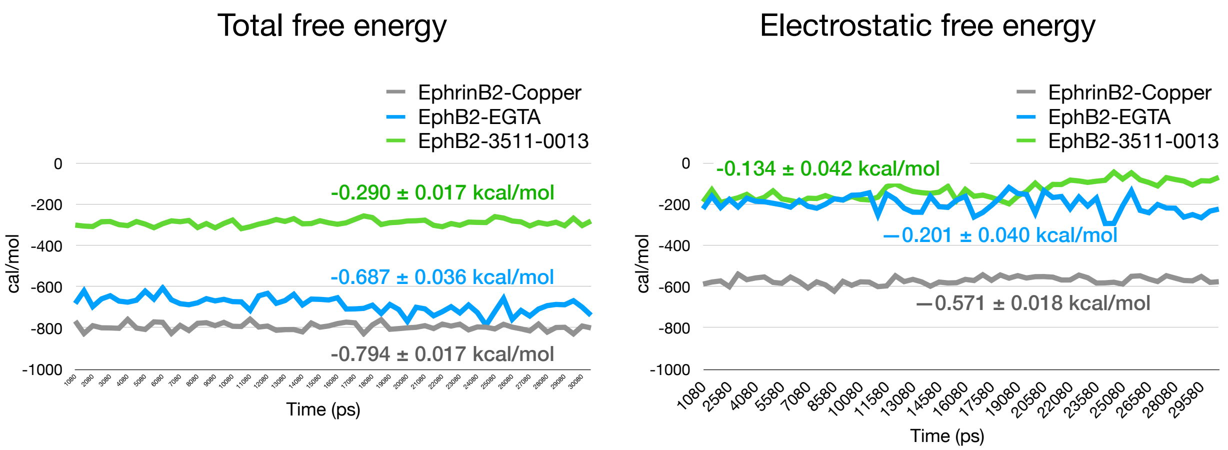


**Figure S3D.** Total and electrostatic PBSA free energy (ΔG) was calculated every 50 ps from a 30 ns NMDA simulation of $Cu^{+2}$ interacting with EphrinB2. Total binding energy shows the copper exhibits a higher overall affinity for EphrinB2 throughout the 30 ns simulation compared to the binding of metal chelators EGTA or 3511-0013 to EphB2. Copper exhibited very strong electrostatic interactions with EphrinB2.

NAMD simulations (30 ns) - Distance apart

EphrinB2 binding to copper during 30 ns simulation - Distance apart over time

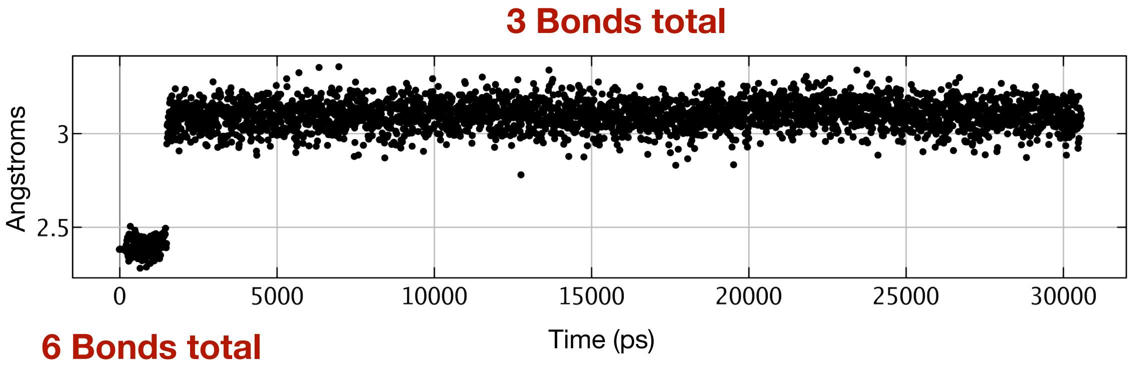


**Figure S3E.** Distance between copper ion and the hydroxyl oxygen on Asp69 of EphrinB2 was calculated every 50 ps from the 30 ns simulation. Data indicates that copper is extremely stable and does not move from the EphrinB2 C-D loop. It is also observed that copper quickly shifted from 6 bonds with EphrinB2 (trigonal pyramidal geometry) to 3 bonds with EphrinB2 (trigonal planar) to reach equilibrium around the 2 ns mark.

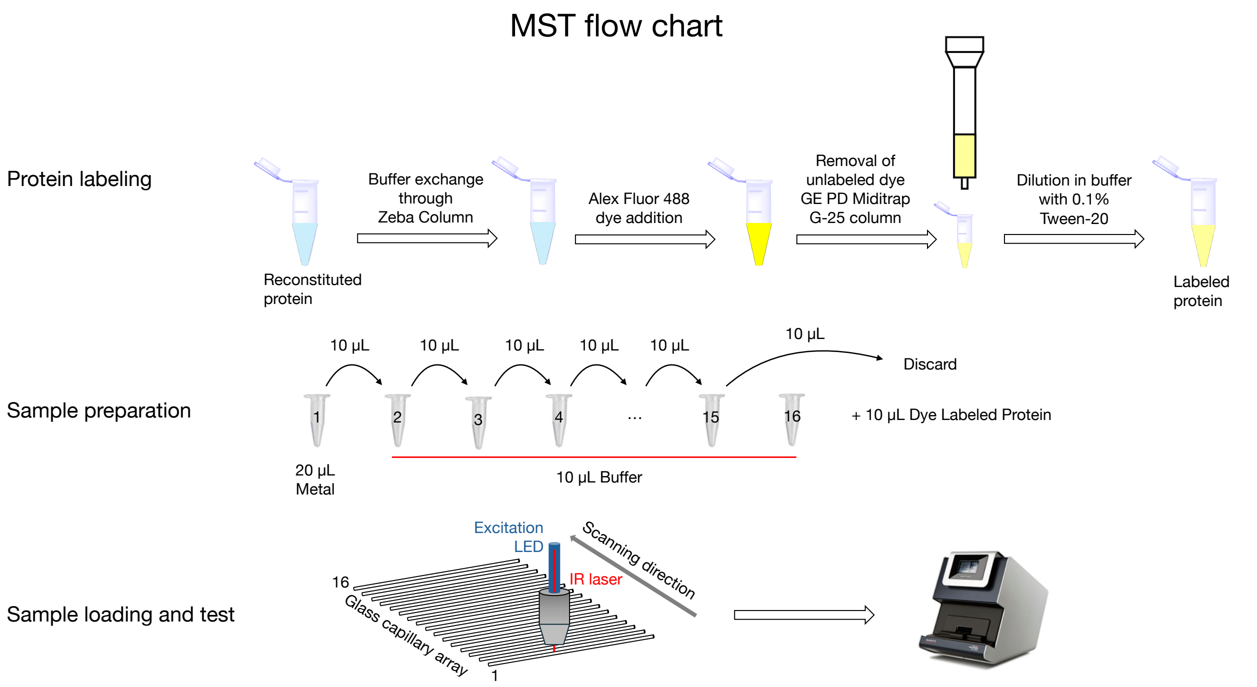


**Figure S3F.** A flow chart of MST experimental procedures, including protein labeling using Alexa Fluor™ 488 NHS Ester (Succinimidyl Ester, from Invitrogen, #A2000), sample preparation, sample loading and test.

### EphrinB2 protein labeling of Alexa Fluor 488 dye

Fluorescence absorbance of dye labeled protein

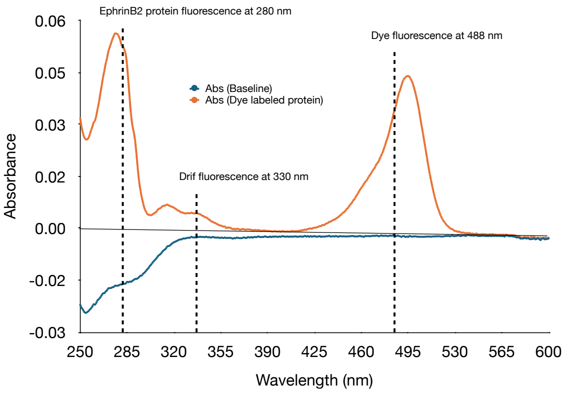


Calculation of dye to protein ratio:

$$\text{Flourescence at 280 from dye} = \frac{0.540}{10^6} * 12400 = 0.0067$$

$$[\text{Dye}] = \frac{0.0394}{73000} * 10^6 = 0.540 \text{ uM}$$

$$[\text{EphrinB2}] = \frac{0.05756 - 2(0.00554) - 0.0067}{35785} * 10^6 = 1.423 \text{ uM}$$

$$\frac{[\text{Dye}]}{[\text{EphrinB2}]} = \frac{0.54}{1.423} = 0.38$$

| | Molar Extinction Coefficient $M^{-1}cm^{-1}$ |
|---|---|
| Dye at wavelength 488 nm | 73000 |
| Dye at wavelength 280 nm | 12400 |
| EphrinB2 protein (Uniprot: P52799) | 35785 |

| | Fluorescence reading (baseline subtracted) |
|---|---|
| Dye at wavelength 488 nm | 0.0394 |
| EphrinB2 protein at wavelength 280 nm | 0.05756 |
| Drif fluorescence at wavelength 330 nm | 0.01108 |

**Figure S3G.** MST protein labeling of Alex Fluor 488 dye on human EphrinB2 ectodomain protein (Sino Biologicals, #10881-HCCH) demonstrates a ratio of dye to protein of 0.38. Approximately, 38 out of 100 human EphrinB2 proteins are labeled with at least one dye. The molar extinction coefficient for human EphrinB2 protein was estimated using ProPram calculator with amino acids 1-229 as listed in the description of the protein from the supplier. Alexa Fluor™ 488 NHS Ester (Succinimidyl Ester, from Invitrogen, #A2000) was used to couple dye to the ε-amino groups of lysine residues on human EphrinB2 under slightly basic conditions (pH ~8.3). After incubation, unreacted dye was removed by size-exclusion column, leaving dye labeled protein. Spectrophotometry was used to calculate the ratio of dye to protein.

### EphrinB2 binds copper

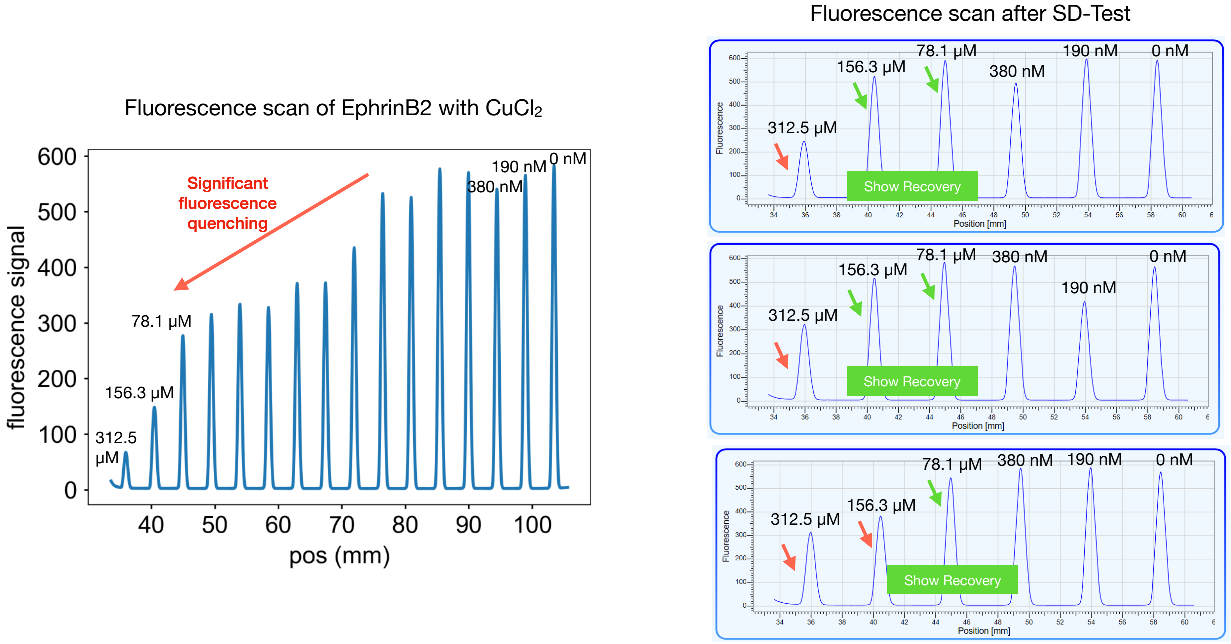


**Figure S3H.** Fluorescence scans of dye labeled human EphrinB2 mixed with increasing concentrations of copper ($CuCl_2$) shows significant quenching of the fluorescent signal with higher copper concentrations (left). SD-Test (SDS denaturation test) shows fluorescence at the higher concentrations of copper is recovered if the sample is heated in detergent to denature the protein. This demonstrates copper specifically binding to the human EphrinB2 ectodomain as identified by the significant quenching of the fluorescence signal and that quenching was due to specific EphrinB2-$Cu^{2+}$ binding rather than non-specific effects such as protein aggregation or adsorption. Fluorescence scans were conducted using a Nano Temper Microscope Thermophoresis instrument. SD-Test was conducted using samples of free protein and protein–metal complexes mixed with an SDS/DTT solution which was heated at 95 °C to fully denature the proteins.

### $CuCl_2$ does not cause general fluorescence quench in MST

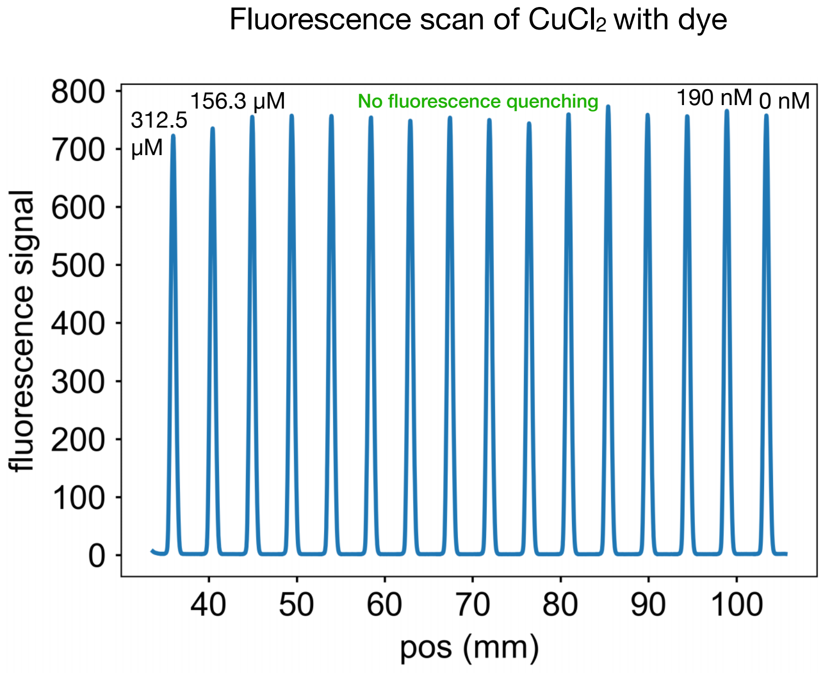


**Figure S3I.** MST of $CuCl_2$ with Alexa Fluor 488 dye demonstrates that $CuCl_2$ does not cause fluorescence quench.

### $CuCl_2$ does not produce fluorescence in MST

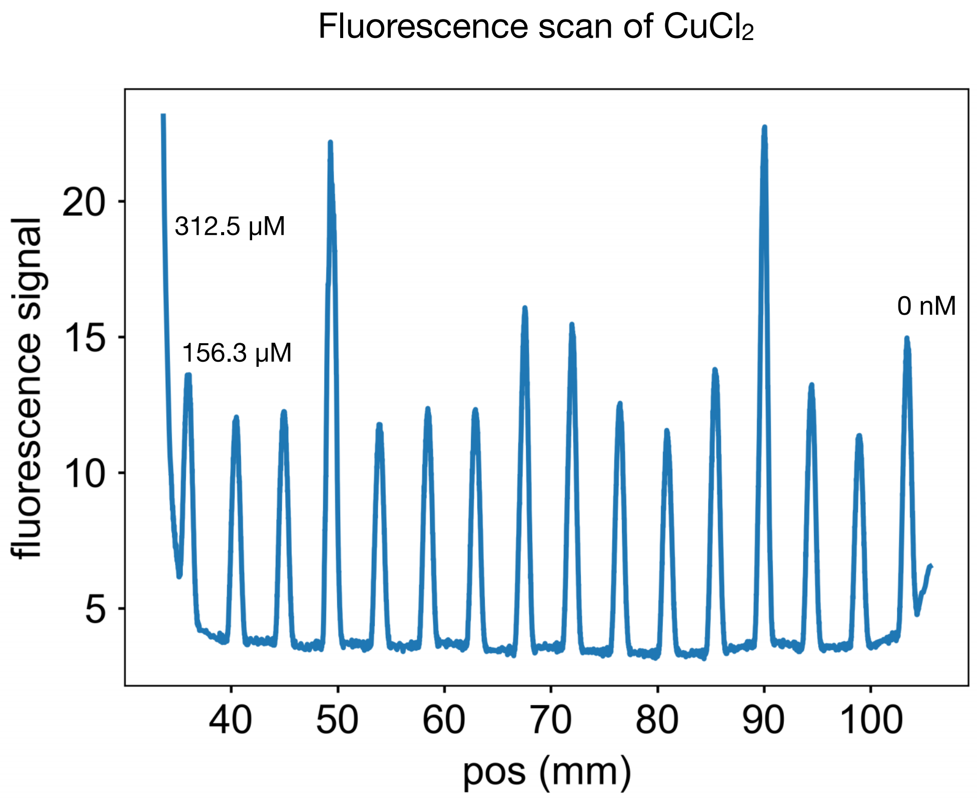


**Figure S3J.** MST of $CuCl_2$ demonstrates that $CuCl_2$ does not produce fluorescence by itself. A insignificant fluorescence signal observed at 0 nM of $CuCl_2$ indicates a background noise.

Cobalt and Nickel do not quench EphrinB2 fluorescence

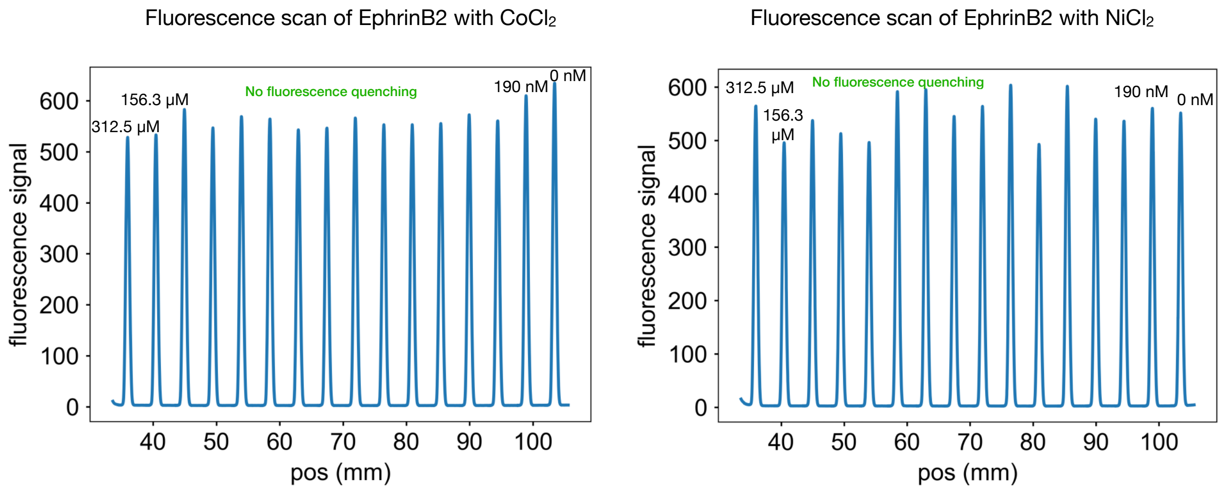


**Figure S3K.** Fluorescence scans of dye labeled human EphrinB2 mixed with increasing concentrations of of cobalt ($CoCl_2$) or nickel ($NiCl_2$) showed no quenching of the fluorescence signal at any concentrations.

#### $CoCl_2$ does not disrupt EphB2-EphrinB tetramerization

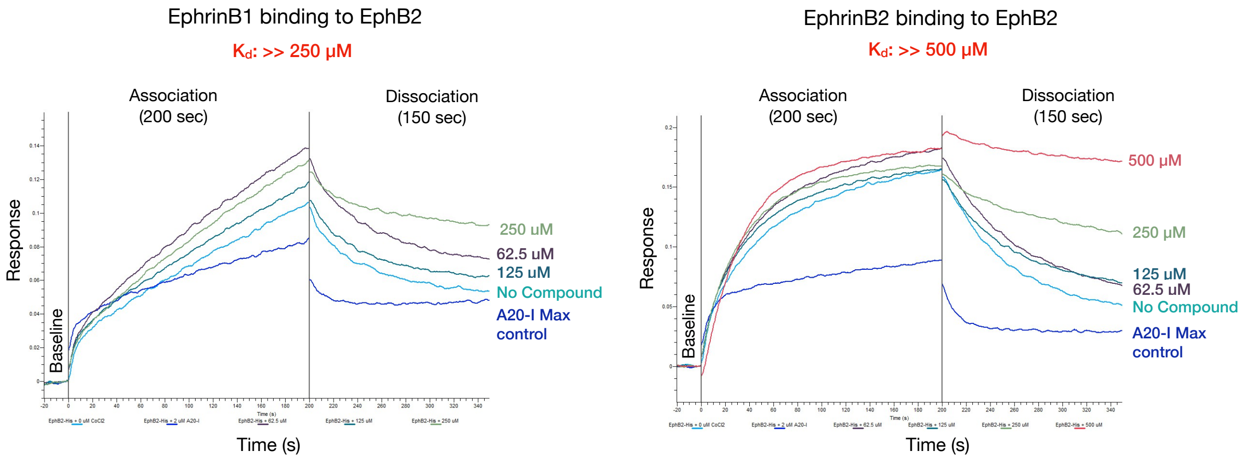


**Figure S3L.** Additional BLI studies of EphrinB1-EphB2 and EphrinB2-EphB2 interactions indicate $CoCl_2$ does not target tetramers at concentrations up to 500 µM. AHC immobilized human EphrinB1-Fc and human EphrinB2-Fc ectodomains binding to 50 nM of the soluble human EphB2-His ectodomain.

#### $NiCl_2$ does not disrupt EphB2-EphrinB tetramerization

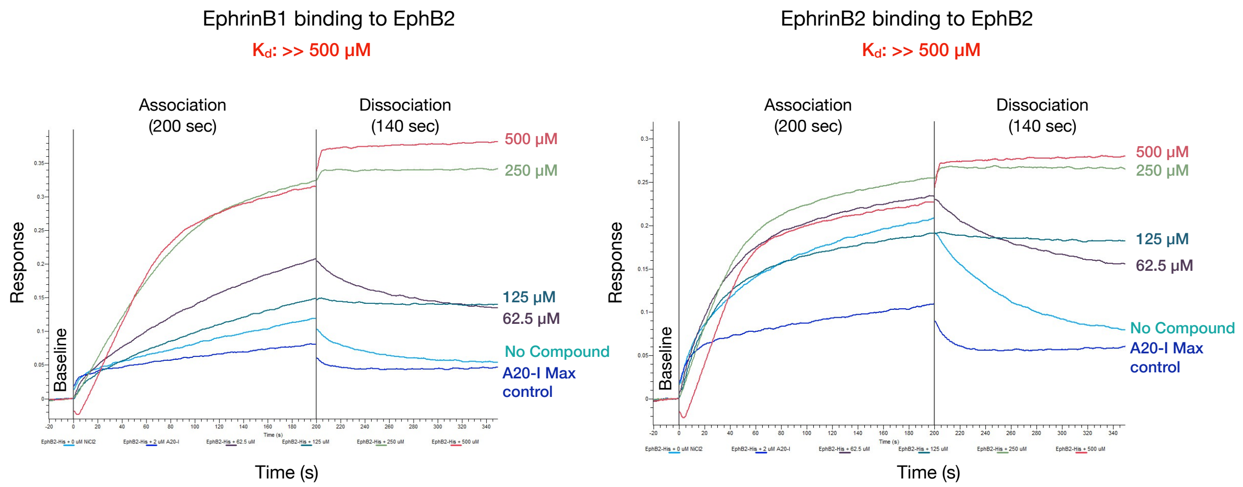


**Figure S3M.** Additional BLI studies of EphrinB1-EphB2 and EphrinB2-EphB2 interactions indicate $NiCl_2$ does not target tetramers at concentrations up to 500 µM. AHC immobilized human EphrinB1-Fc and human EphrinB2-Fc ectodomains binding to 50 nM soluble of the human EphB2-His ectodomain.

### Copper metal binding site on EphrinB2

Assessable Surface Area of lysine residues on EphrinB2

| Res | UID | res_ASA | res_exp |
|---|---|---|---|
| LYS | 71 | 203.3 | 86.32 |
| LYS | 96 | 177.6 | 75.44 |
| LYS | 131 | 112.7 | 47.85 |
| LYS | 133 | 112.6 | 47.81 |
| LYS | 86 | 108.7 | 46.18 |
| LYS | 112 | 99.3 | 42.16 |
| LYS | 95 | 98.0 | 41.62 |
| LYS | 116 | 93.9 | 39.89 |
| LYS | 44 | 70.8 | 30.06 |
| LYS | 60 | 67.8 | 28.78 |
| LYS | 166 | 44.9 | 19.05 |
| LYS | 80 | 29.2 | 12.41 |
| LYS | 67 | 20.4 | 8.68 |
| LYS | 162 | 14.9 | 6.32 |

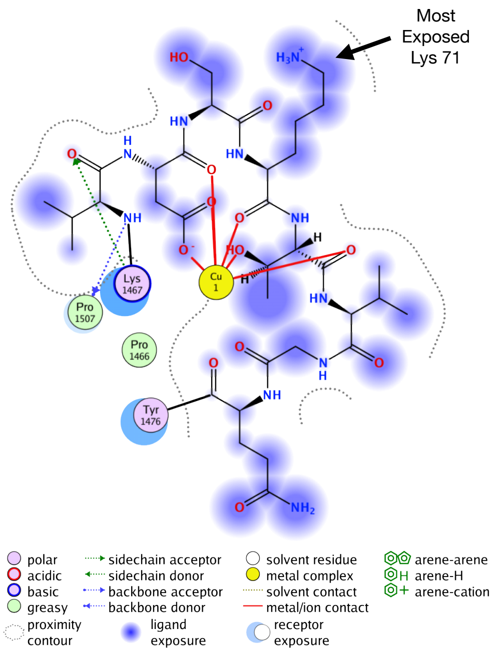


**Figure S3N.** MOE estimates relative accessible surface area (ASA) for all lysines on the mouse EphrinB2 ectodomain domain (1IKO). Protein was protonated at pH 7.4 and then minimized before using the MOE QuickPrep function. The most accessible lysine is Lys71 within the tetramerization interface and within 10 Å from the predicted copper biding site.

### NAMD simulations (30 ns) - EphB2-EphrinB2 dimer and tetramer

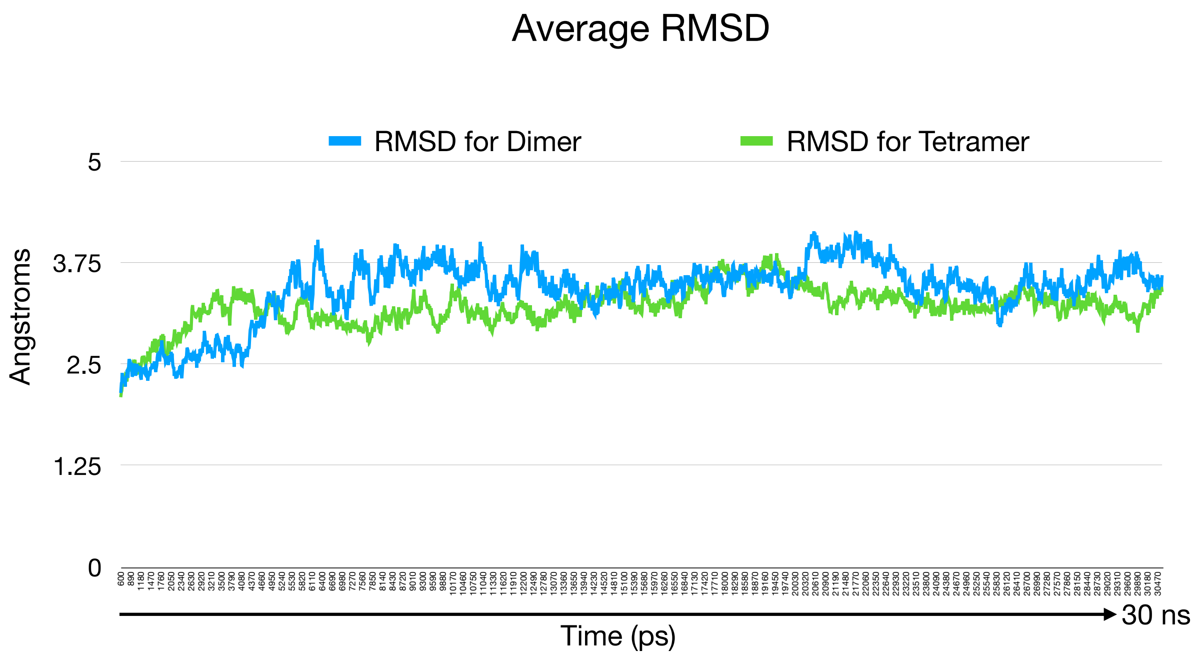


**Figure S4A.** Shown is the average RMSD obtained for EphB2-EphrinB2 dimer and tetramer structures during a 30 ns NAMD simulation. Every 10 ps RMSD was calculated relative to the backbone loops of EphB2 and EphrinB2 to account for global drift of the oligomers during the simulation. Both interactions reached equilibrium with about 3.4 angstroms RMSD after ~5 nanoseconds.

#### NAMD simulations (30 ns) - EphB2-EphrinB2 dimer and tetramer

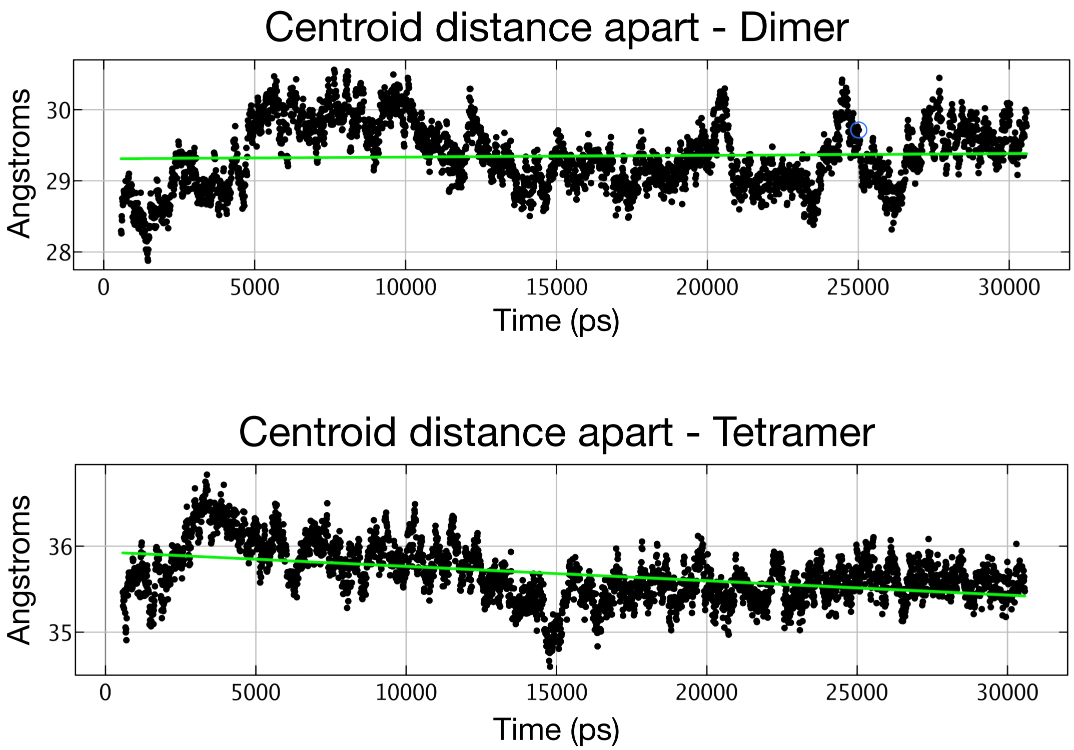


**Figure S4B** Centroid distance apart for the EphB2-EphrinB2 dimer and tetramer structures was calculated every 10 ps from the 30 ns NAMD simulations. After an initial 5 ns to reach equilibrium, the EphB2 and EphrinB2 monomers that form the dimer do not move apart from each other, whereas the two juxtaposed EphB2-EphrinB2 dimers in the tetramer seem to be moving slightly closer together with time.

#### NAMD simulations (30 ns) - EphB2-EphrinB2 dimer and tetramer free energy (ΔG)

| EphB2-EphrinB2 interaction | Free Energy (ΔG) computationally simulated by NAMD |
|---|---|
| Dimer | -6.588 kcal/mol |
| Tetramer | -10.519 kcal/mol |

The relative stability ratio between dimer and tetramer was calculated using the following equations at 300 K: ($K_{a1}$ and $\Delta G_1$ are for tetramer while $\Delta G_2$ and $K_{a2}$ are for dimer)

Equilibrium constant equation: $K_a = e^{-\Delta G/RT}$

$$\text{Relative stability ratio} = \frac{K_{a1}}{K_{a2}} = e^{-(\Delta G_1 - \Delta G_2)}/RT$$

$$\text{Relative stability ratio} = \frac{K_{a1}}{K_{a2}} = e^{-(-10.519-(-6.588))}/(0.001987 * 300) = 773.9$$

Hence, the tetramer is >750x more stable than the dimer.

**Figure S4C.** Free energies of EphB2–EphrinB2 dimer and tetramer interactions were estimated using molecular dynamics simulations in phosphate-buffered saline (PBS) solvent at 300 K over a 30 ns simulation window. The computed binding free energies were –6.588 kcal/mol for the dimer and –10.519 kcal/mol for the tetramer, yielding a relative free energy difference (ΔΔG) of 3.931 kcal/mol. The equilibrium association constant was calculated using R=0.001987 kcal $mol^{-1}$ $K^{-1}$ and T=300K. The resulting relative stability ratio ($K_{a1}/K_{a2}$) indicates that the tetramer is >750 times more stable than the dimer under these conditions.

#### EphB2-EphrinB2 tetramerization is effectively irreversible at physiological temperatures

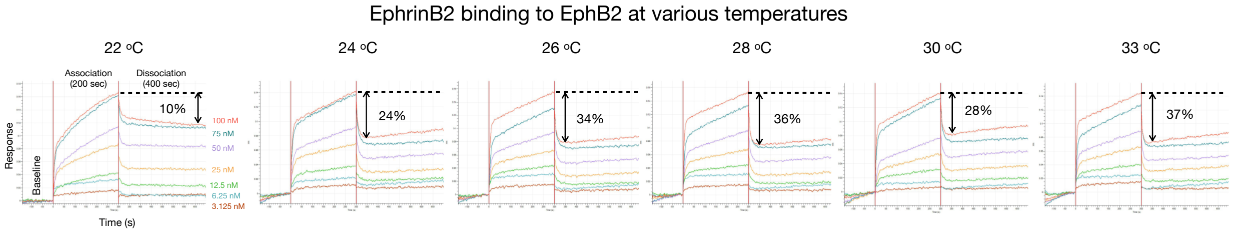


Eph monomer + Ephrin monomer $\underset{k_{dis2}}{\overset{k_{a2}}{\rightleftharpoons}}$ Dimer + Heat

Dimer + Dimer $\underset{k_{dis1}}{\overset{k_{a1}}{\rightleftharpoons}}$ Tetramer + Heat

| | 22 °C | | 24 °C | | 26 °C | | 28 °C | | 30 °C | | 33 °C | |
|---|---|---|---|---|---|---|---|---|---|---|---|---|
| No change | $K_{D1}$ | <1.0E-12 | $K_{D1}$ | <1.0E-12 | $K_{D1}$ | <1.0E-12 | $K_{D1}$ | <1.0E-12 | $K_{D1}$ | <1.0E-12 | $K_{D1}$ | <1.0E-12 |
| ↑ | $K_{D2}$ | 1.24E-09 | $K_{D2}$ | 1.74E-08 | $K_{D2}$ | 1.63E-08 | $K_{D2}$ | 2.02E-08 | $K_{D2}$ | 2.70E-08 | $K_{D2}$ | 2.9E-08 |
| No change | $k_{a1}$ | 1.57E+05 | $k_{a1}$ | 2.12E+05 | $k_{a1}$ | 1.85E+05 | $k_{a1}$ | 1.85E+05 | $k_{a1}$ | 1.85E+05 | $k_{a1}$ | 2.18E+05 |
| ↑ | $k_{a2}$ | 3.21E+06 | $k_{a2}$ | 6.73E+06 | $k_{a2}$ | 8.16E+06 | $k_{a2}$ | 6.45E+06 | $k_{a2}$ | 4.83E+06 | $k_{a2}$ | 1.66+07 |
| No change | $k_{dis1}$ | <1.0E-07 | $k_{dis1}$ | <1.0E-07 | $k_{dis1}$ | <1.0E-07 | $k_{dis1}$ | <1.0E-07 | $k_{dis1}$ | <1.0E-07 | $k_{dis1}$ | <1.0E-07 |
| ↑ | $k_{dis2}$ | 3.99E-03 | $k_{dis2}$ | 1.17E-01 | $k_{dis2}$ | 1.33E-01 | $k_{dis2}$ | 1.30E-01 | $k_{dis2}$ | 1.31E-01 | $k_{dis2}$ | 1.67E-01 |

**Figure S4D.** BLI studies of EphB2-EphrinB2 interactions indicate temperature affects dimerization kinetics with minimal effect on tetramerization dynamics. Shown are BLI sensorgram traces of AHC immobilized human EphrinB2-Fc ectodomain binding to soluble human EphB2-His ectodomain at various temperatures including 22, 24, 26, 28, 30, and 33 °C. The concentrations of EphB2 were 3.125, 6.25, 12.5, 25, 50, 75, and 100 nM for each temperature, allowing for global fit 2:1 calculations of $K_D$, $k_a$, and $k_{dis}$ for both dimer and tetramer. The data indicates the EphB2-EphrinB2 tetramer is more stable than the dimer.

BLI experimentally determined free energy of EphB2-EphrinB2 dimer and tetramer

| EphB2-EphrinB2 interaction | $K_D$ determined by BLI | Free Energy (ΔG) |
|---|---|---|
| Dimer | 70 nM | -9.821 kcal/mol |
| Tetramer | 0.28 nM | -13.112 kcal/mol |

Free energy was calculated using the following equation at 300 K:

$$\Delta G_{dimer} = RTln(KD) = (0.001987 * 300)ln(70 * 10^{-9}) = -9.821$$

$$\Delta G_{tetramer} = RTln(KD) = (0.001987 * 300)ln(2.8 * 10^{-10}) = -13.112$$

The relative stability of dimer and tetramer was calculated using the following equation:
($K_{a1}$ and $\Delta G_1$ are for tetramer while $\Delta G_2$ and $K_{a2}$ are for dimer)

$$\text{Relative stability ratio} = \frac{K_{a1}}{K_{a2}} = e^{-(\Delta G_1 - \Delta G_2)}/RT = e^{-(-13.112-(-9.821))}/(0.001987 * 300) = 250$$

Hence, the tetramer is ~250x more stable than the dimer given experimental data.

**Figure S4E.** Free energies were derived from experimentally measured equilibrium dissociation constants ($K_D$) obtained by BLI studies of the EphB2–EphrinB2 interaction. Calculations were performed at 300 K using the thermodynamic relation ΔG = RTln(KD), yielding binding free energies of –9.821 kcal/mol for the dimer and –13.112 kcal/mol for the tetramer. The relative free energy difference (ΔΔG = –3.291 kcal/mol) corresponds to a ~250-fold increase in stability of the tetramer compared to the dimer. Because the $K_{D1}$ and $K_{dis1}$ values determined using BLI for the EphB2-EphrinB2 tetramer in the experiments presented in Figures 4D and S4D indicated irreversible formation of the tetramer at all temperatures tested, here for calculations of free energy, we used a $K_{D1}$ = 0.28 nM for tetramer and $K_{D2}$ = 70 nM for the dimer. These more conservative $K_D$ values are what is reported for the EphB2-EphrinB2 interaction in the accompanying Wang, Khambete, et al manuscript (submitted) to ensure the free energy calculations are not over-estimated.

NAMD simulations (30 ns) - EphB2-EphrinB2 vs EphB4-EphrinB2 dimers and tetramers

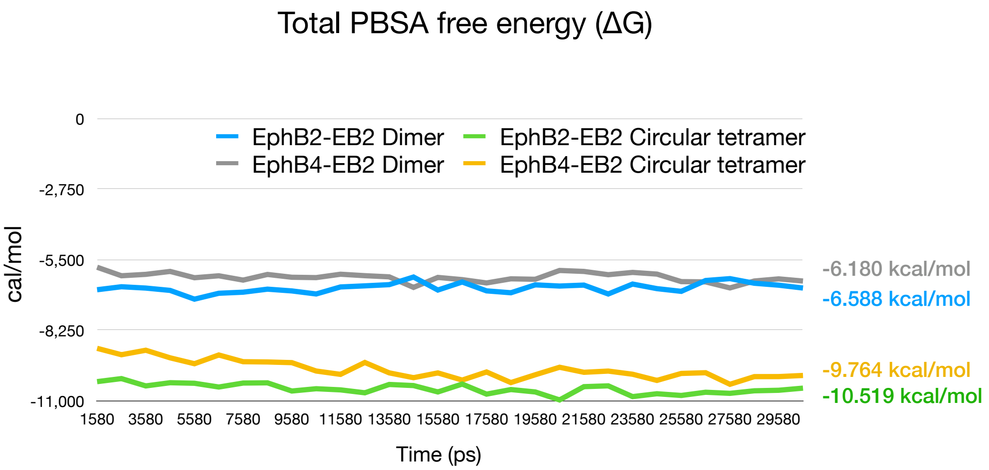


**Figure S4F.** Total PBSA free energies (ΔG) of EphB2-EphrinB2 and EphB4-EphrinB2 dimer/tetramer binding interactions were calculated every 50 picoseconds from 30 nanosecond NAMD simulations in phosphate-buffered saline solvent. The data indicates tetramer interactions for both EphB2-EphrinB2 (orange) and EphB4-EphrinB2 (green) are thermodynamically much more stable than their corresponding dimer interactions. Regarding the two different tetramer interactions, the free energy difference (ΔΔG) of 0.755 kcal/mol, provides a relative stability ratio ($K_{a1}/K_{a2}$) (calculated from equation in Figure S4C using R=0.001987 kcal $mol^{-1}$ $K^{-1}$ and T=300K) that indicates the EphB2-EphrinB2 tetramer is 3.55x more stable than the EphB4-EphrinB2 tetramer under these conditions. Additional calculations from these simulations indicate the EphB2-EphrinB2 tetramer is 774x more stable than its dimer as shown in Figure S4C, while the EphB4-EphrinB2 tetramer is 409x more stable than its corresponding dimer.

#### Interactions in EphB2 tetramerization interface with EphrinB2

| Type | ChainA | PosA | SetA | ChainB | PosB | SetB | Energy | Dist | BB | Freq | Cons | Area |
|---|---|---|---|---|---|---|---|---|---|---|---|---|
| DIH | EphB2 | 61 | Arg87 | EphrinB2 | 39 | Asp69 | -19.37 | 3.67 | -- | 11 | 1 | 26.03 |
| D | EphB2 | 109 | Phe135 | EphrinB2 | 4 | Glu34 | -4.04 | 3.89 | -- | 26 | 1 | 50.13 |
| DH | EphB2 | 153 | Arg179 | EphrinB2 | 13 | Ser43 | -3.22 | 3.68 | -- | 4 | 1 | 34.31 |
| D | EphB2 | 102 | Phe128 | EphrinB2 | 7 | Tyr37 | -2.56 | 3.99 | -* | 25 | 1 | 39.86 |
| D | EphB2 | 102 | Phe128 | EphrinB2 | 12 | Asn42 | -2.27 | 3.91 | -* | 8 | 1 | 25.88 |
| D | EphB2 | 63 | Arg89 | EphrinB2 | 42 | Thr72 | -1.35 | 3.84 | b- | 2 | 1 | 14.45 |
| D | EphB2 | 106 | Thr132 | EphrinB2 | 5 | Pro35 | -1.16 | 4.01 | -- | 2 | 1 | 25.59 |
| D | EphB2 | 102 | Phe128 | EphrinB2 | 11 | Ser41 | -1.00 | 4.00 | -* | 4 | 1 | 20.57 |
| D | EphB2 | 61 | Arg87 | EphrinB2 | 42 | Thr72 | -0.65 | 4.13 | -- | 4 | 1 | 17.48 |
| D | EphB2 | 104 | Leu130 | EphrinB2 | 14 | Lys44 | -0.56 | 4.10 | -- | 1 | 1 | 9.95 |
| D | EphB2 | 154 | Asn180 | EphrinB2 | 42 | Thr72 | -0.44 | 4.23 | *- | 6 | 1 | 24.95 |
| D | EphB2 | 104 | Leu130 | EphrinB2 | 4 | Glu34 | -0.29 | 4.11 | *- | 4 | 1 | 8.70 |
| D | EphB2 | 106 | Thr132 | EphrinB2 | 3 | Leu33 | -0.25 | 3.86 | -b | 1 | 1 | 5.14 |
| D | EphB2 | 101 | Asp127 | EphrinB2 | 11 | Ser41 | -0.20 | 3.92 | -* | 6 | 1 | 13.24 |
| D | EphB2 | 62 | Arg88 | EphrinB2 | 42 | Thr72 | -0.15 | 4.39 | b- | 1 | 1 | 0.20 |
| D | EphB2 | 101 | Asp127 | EphrinB2 | 13 | Ser43 | 0.00 | 4.09 | -- | 1 | 1 | 3.77 |
| D | EphB2 | 108 | Thr134 | EphrinB2 | 2 | Val32 | 0.00 | 4.22 | -b | 1 | 1 | 11.76 |
| D | EphB2 | 101 | Asp127 | EphrinB2 | 12 | Asn42 | 0.01 | 4.27 | -b | 1 | 1 | 0.69 |
| D | EphB2 | 103 | Asp129 | EphrinB2 | 7 | Tyr37 | 0.06 | 4.18 | *- | 2 | 1 | 26.58 |
| D | EphB2 | 154 | Asn180 | EphrinB2 | 11 | Ser41 | 0.15 | 4.02 | -- | 2 | 1 | 23.43 |
| D | EphB2 | 104 | Leu130 | EphrinB2 | 5 | Pro35 | 0.15 | 4.15 | b* | 6 | 1 | 17.45 |
| D | EphB2 | 61 | Arg87 | EphrinB2 | 41 | Lys71 | 1.44 | 3.82 | -- | 4 | 1 | 18.36 |
| D | EphB2 | 104 | Leu130 | EphrinB2 | 7 | Tyr37 | 2.34 | 3.89 | *- | 6 | 1 | 21.14 |
| D | EphB2 | 63 | Arg89 | EphrinB2 | 41 | Lys71 | 5.07 | 3.94 | -- | 10 | 1 | 57.53 |

**Figure S4G.** Interactions in EphB2 tetramerization interface with EphrinB2 are listed in the table. The interactions from Arg87, Arg88, Arg89, and Arg179 of EphB2 are highlighted yellow.

#### Conformational changes in key loops of EphB2 and EphrinB2 upon dimerization and tetramerization

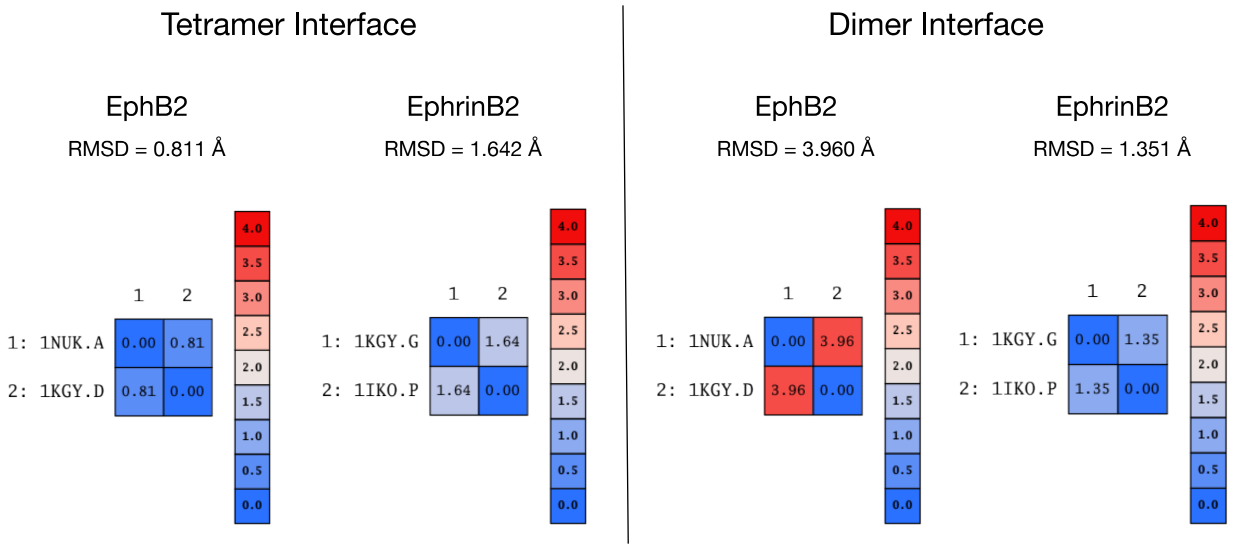


**Figure S4H.** The EphB2 and EphrinB2 structures from the EphB2-EphrinB2 circular tetramer crystal structure (1KGY) were extracted as individual proteins and compared to the EphB2 monomer crystal structure (1NUK) or EphrinB2 monomer crystal structure (1IKO), respectively. No protonation or minimization was performed to maintain the original crystal structures. MOE sequence alignment compared the two structures (tetramer-bound and monomer) and the RMSD values for loop backbones in the dimer and tetramer interfaces determined. The data indicates that the EphB2 and EphrinB2 tetramer interface loops do not change in conformation significantly, however, the EphB2 dimerization loop changes conformation dramatically, pointing towards the flexible, hydrophobicity-driven dimer interface of EphB2.

EphB2 and EphrinB2 amino acid R-group conformational changes upon dimerization

EphB2 dimerization interface | EphrinB2 dimerization interface

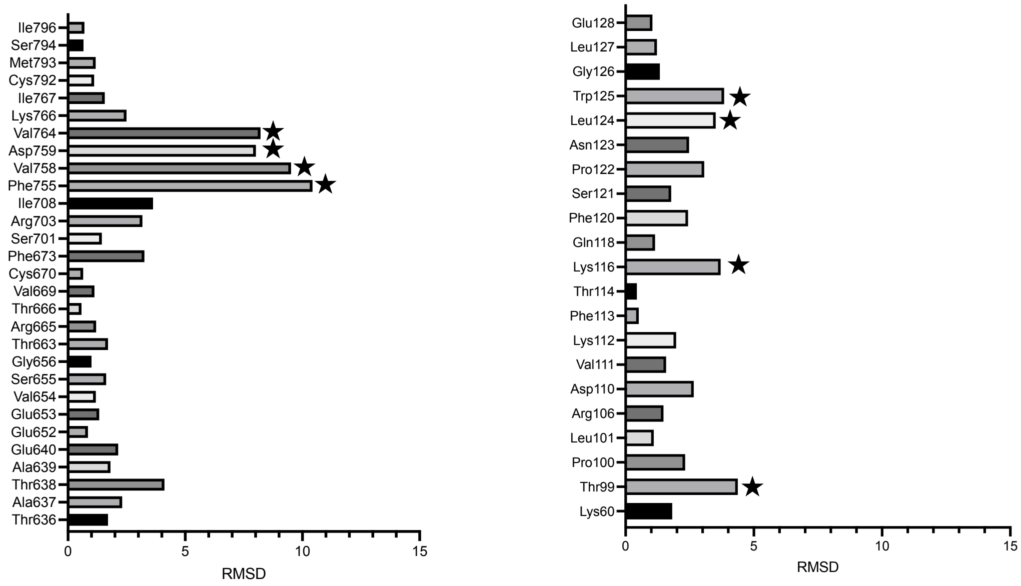


**Figure S4I.** A specialized MOE script was utilized to calculate the RMSD fluctuations of the R-groups of specific amino acids in the dimerization interface of EphB2 and EphrinB2, comparing their position in the respective unbound monomer with that found in the EphB2-EphrinB2 dimer. Overall, the amino acid R-groups of the EphB2 dimerization interface fluctuate significantly more than those from the cognate EphrinB2 dimerization interface.

EphB2 and EphrinB2 amino acid R-group conformational changes upon tetramerization

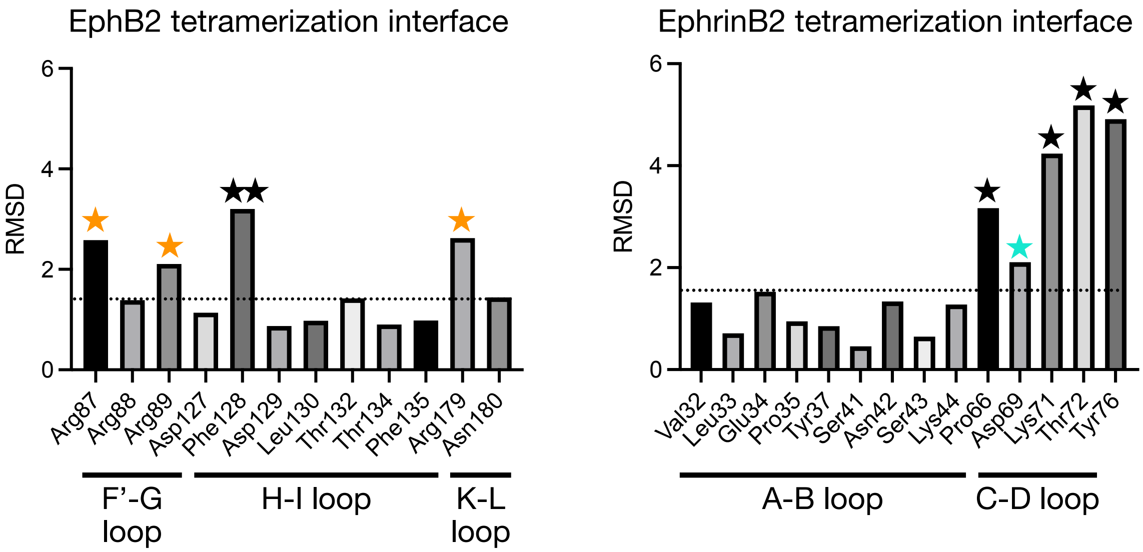


**Figure S4J.** A specialized MOE script was used to calculate the RMSD fluctuations of the R-groups of specific amino acids in the tetramerization interface of EphB2 and EphrinB2, comparing respective unbound tetramer interface residues with those found in the EphB2-EphrinB2 circular tetramer. Overall, the electropositive Arg87, Arg89, and Arg179 and the previously noted Phe128 (in Figure 2A) in the EphB2 tetramerization interface show significant movement following tetramer formation. Asp69 and surrounding Lys71, Thr72, and the previously noted Pro66 (in Figure 2A) in the EphrinB2 tetramerization interface also show significant movement following tetramer formation, as does Tyr76, another residue in the C-D loop immediately adjacent to the EphrinB2 tetramerization interface.

#### EphB2-EphrinB2 dimers are needed for tetramer formation

Binding sites in EphB2 monomer predicted by MOE SiteFinder

| Site | Size | PLB | Hyd | Location |
|---|---|---|---|---|
| 1 | 110 | 2.60 | 39 | Dimer |
| 2 | 45 | 0.55 | 11 | Other |
| 3 | 15 | 0.00 | 5 | Other |
| 4 | 18 | -0.15 | 5 | Other |
| 5 | 8 | -0.32 | 6 | Tetramer |
| 6 | 5 | -0.58 | 6 | Other |
| 7 | 17 | -0.61 | 8 | Other |
| 8 | 20 | -0.63 | 6 | Other |
| 9 | 6 | -0.85 | 2 | Other |

Binding sites in EphrinB2 monomer predicted by MOE SiteFinder

| Site | Size | PLB | Hyd | Residues |
|---|---|---|---|---|
| 1 | 27 | 2.27 | 8 | Tetramer |
| 2 | 23 | 1.55 | 5 | Other |
| 3 | 16 | 1.31 | 7 | Other |
| 4 | 29 | 0.42 | 13 | Dimer |
| 5 | 9 | 0.28 | 5 | Other |
| 6 | 10 | -0.07 | 7 | Other |
| 7 | 15 | -0.42 | 8 | Other |
| 8 | 4 | -0.46 | 3 | Other |
| 9 | 10 | -0.67 | 3 | Other |
| 10 | 4 | -0.69 | 5 | Other |
| 11 | 8 | -0.74 | 11 | Other |
| 12 | 15 | -0.93 | 6 | Other |
| 13 | 11 | -0.93 | 6 | Other |
| 14 | 4 | -0.93 | 5 | Other |

Binding sites in EphB2-EphrinB2 dimer predicted by MOE SiteFinder

| Site | Size | PLB | Hyd | Residues |
|---|---|---|---|---|
| 1 | 49 | 2.94 | 18 | EphrinB2 Tetramer Interface |
| 2 | 42 | 1.60 | 10 | Other |
| 3 | 55 | 1.37 | 19 | Other |
| 4 | 15 | 0.69 | 5 | Other |
| 5 | 20 | 0.28 | 7 | Other |
| 6 | 29 | 0.10 | 7 | Other |
| 7 | 14 | -0.22 | 6 | Other |
| 8 | 13 | -0.47 | 6 | Other |
| 9 | 16 | -0.48 | 10 | Other |
| 10 | 13 | -0.48 | 6 | Other |
| 11 | 13 | -0.48 | 4 | Other |
| 12 | 22 | -0.59 | 4 | Other |
| 13 | 11 | -0.64 | 14 | Other |
| 14 | 7 | -0.68 | 9 | Other |
| 15 | 11 | -0.71 | 6 | Other |
| 16 | 10 | -0.73 | 3 | Other |
| 17 | 11 | -0.74 | 2 | Other |
| 18 | 10 | -0.77 | 4 | Other |

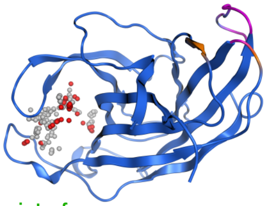


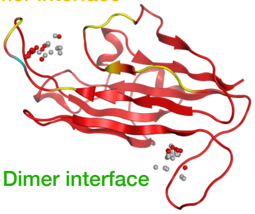


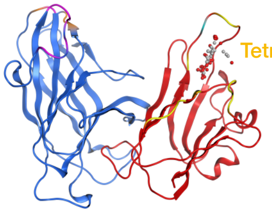


**Figure S4K.** EphB2 ectodomain monomer (1NUK), EphrinB2 ectodomain monomer (1IKO), and the EphB2-EphrinB2 dimer (1KGY) were protonated and minimized in MOE and then the SiteFinder tool was used to discover potential binding pockets on the surface of the different proteins. SiteFinder is a common reference in computational biology and drug discovery as it identifies, sizes, and ranks possible binding pockets on the surface of a protein of interest that might interact with other ligand molecules, such as another protein or a small molecule. The Propensity for Ligand Binding (PLB) scores are representative of how likely a ligand molecule is predicted to bind to that particular site. The location of each predicted binding site was confirmed to be “dimer” interface, “tetramer” interface, or “other” based on the amino acids contained within the binding site. This analysis revealed the largest ligand binding site in the EphB2 monomer is the known dimerization interface, which has the largest PLB score and size, indicating that the dimer interaction is favorable. MOE SiteFinder analysis of the EphrinB2 monomer identified the tetramer interface of EphrinB2 as the most likely available binding site. MOE SiteFinder analysis of the EphB2-EphrinB2 dimer again identified the tetramer interface of EphrinB2 as the most likely available binding site, which is consistent with the dimer-binding sites of the EphB2 and EphrinB2 monomers now bound together and not available for interactions. Altogether, the SiteFinder results are consistent with the general idea that EphB2 and EphrinB2 monomers first interact to form a dimer, and then two dimers would be available to align together in a precise, tandem head-to-tail like fashion to juxtapose the two cognate tetramer interfaces of the opposing dimers to snap them into the circular tetramer.

#### Specific amino acids for each binding pocket identified in EphB2 monomer

##### EphB2 Monomer

| Site | Size | PLB | Hyd | Side | Residues |
|---|---|---|---|---|---|
| 1 | 110 | 2.60 | 39 | 60 | 1:(VAL54 SER55 GLY56 TYR57 ASP58 GLU59 ASN60 MET61 ILE64 THR66 GLN68 VAL69 CYS70 SER101 ARG103 PHE155 VAL158 ASP159 LEU160 GLY161 GLY162 ARG163 VAL164 LYS166 ILE167 CYS192 MET193 SER194 ILE196) |
| 2 | 45 | 0.55 | 11 | 24 | 1:(GLY90 ALA91 HIS92 ARG93 ILE94 HIS95 TYR202 ARG203 LYS204 CYS205 PRO206 ARG207) |
| 3 | 15 | 0.00 | 5 | 14 | 1:(GLY42 TRP43 MET44 ARG82 THR83 LYS84 PHE85 ASP129 ALA131 TYR183) |
| 4 | 18 | -0.15 | 5 | 11 | 1:(ASN71 VAL72 PHE73 GLU74 SER75 SER76 GLN77 PRO112 SER114 LYS116 TYR189 GLY190) |
| 5 | 8 | -0.32 | 6 | 15 | 1:(ARG87 ARG88 ARG89 GLY90 ARG179 ASN180) |
| 6 | 5 | -0.58 | 6 | 12 | 1:(MET32 ALA37 ALA39 GLU40 LEU41 TRP43 LYS84) |
| 7 | 17 | -0.61 | 8 | 18 | 1:(THR35 TYR57 ASP58 GLU59 ASN62 THR63 ILE64 LYS99 ILE196 ALA197) |
| 8 | 20 | -0.63 | 6 | 16 | 1:(PRO48 SER49 GLY50 TRP51 GLU52 GLN68 VAL69 CYS70 ASN71 PHE73 GLU74) |
| 9 | 6 | -0.85 | 2 | 7 | 1:(ASP152 GLU153 MET165 LYS166 ILE167 ASN168) |

**Figure S4K** (continued). Amino acid sequences of the binding pockets identified by SiteFinder for the EphB2 monomer (1NUK).

#### Specific amino acids for each binding pocket identified in EphrinB2 monomer

EphrinB2 Monomer

| Site | Size | PLB | Hyd | Side | Residues |
|---|---|---|---|---|---|
| 1 | 27 | 2.27 | 8 | 18 | 3:(ASN39 SER40 SER41 PRO66 LYS67 VAL68 ASP69 LYS71 THR72 GLY74 GLN75 GLU77) |
| 2 | 23 | 1.55 | 5 | 13 | 3:(MET83 VAL84 ASP85 GLN88 GLU97 ASN98 THR99) |
| 3 | 16 | 1.31 | 7 | 16 | 3:(ASN39 SER40 SER41 ASN42 SER43 PHE45 ARG59 MET61) |
| 4 | 29 | 0.42 | 13 | 20 | 3:(PHE113 THR114 ILE115 LYS116 GLN118 SER121 PRO122 ASN123 GLY126 LEU127 PHE129) |
| 5 | 9 | 0.28 | 5 | 9 | 3:(GLY48 ALA89 ASP90 ARG91 CYS92 CYS156 GLN157 ALA160 MET161 LYS162) |
| 6 | 10 | -0.07 | 7 | 11 | 3:(MET83 VAL84 ASP85 LYS86 ASP134 TYR135 TYR136) |
| 7 | 15 | -0.42 | 8 | 10 | 3:(GLY49 GLY50 LEU51 VAL52 LYS162 ILE163 LEU164) |
| 8 | 4 | -0.46 | 3 | 5 | 3:(GLN75 TYR76 TYR78 GLY143 SER144 LEU145) |
| 9 | 10 | -0.67 | 3 | 11 | 3:(SER40 GLU77 TYR79 ASN142 GLY143 VAL155 ARG159) |
| 10 | 4 | -0.69 | 5 | 9 | 3:(GLY48 GLY49 ASP90 TYR136 LYS162 LEU164) |
| 11 | 8 | -0.74 | 11 | 15 | 3:(LEU33 GLU34 ILE36 LYS44 LEU53) |
| 12 | 15 | -0.93 | 6 | 13 | 3:(TYR37 ASN39 ASN42 LYS44 PRO66) |
| 13 | 11 | -0.93 | 6 | 11 | 3:(TYR37 ILE64 PRO66 ASP110 LYS112) |
| 14 | 4 | -0.93 | 5 | 5 | 3:(PHE113 TRP125 GLY126 LEU127) |

**Figure S4K** (continued). Amino acid sequences of the binding pockets identified by SiteFinder for the EphrinB2 monomer (1IKO).

#### Specific amino acids for each binding pocket identified in EphB2-EphrinB2 dimer

##### EphB2-EphrinB2 Dimer

| Site | Size | PLB | Hyd | Side | Residues |
|---|---|---|---|---|---|
| 1 | 49 | 2.94 | 18 | 34 | 3:(TRP38 ASN39 SER40 SER41 CYS65 PRO66 LYS67 VAL68 ASP69 LYS71 THR72 GLY74 GLN75 TYR76 GLU77 TYR79) |
| 2 | 42 | 1.60 | 10 | 23 | 1:(GLY90 ALA91 HIS92 ARG93 ILE94 HIS95 TYR202 ARG203 LYS204 CYS205 PRO206 ARG207) |
| 3 | 55 | 1.37 | 19 | 36 | 1:(PRO48 SER49 GLY50 TRP51 GLU52 GLN68 VAL69 CYS70 ASN71 PHE73 GLU74)3:(ILE57 LYS116 GLN118 GLU119 PHE120 SER121 PRO122) |
| 4 | 15 | 0.69 | 5 | 14 | 1:(GLY42 TRP43 MET44 ARG82 THR83 LYS84 PHE85 ASP129 ALA131 TYR183) |
| 5 | 20 | 0.28 | 7 | 15 | 3:(ASN39 SER40 SER41 ASN42 SER43 PHE45 ARG159 MET161) |
| 6 | 29 | 0.10 | 7 | 20 | 1:(THR35 THR36 THR38 ASP58 ASN62 THR63 ILE64)3:(LEU101 LEU102 ASN103 ARG106 VAL111) |
| 7 | 14 | -0.22 | 6 | 13 | 1:(MET32 ASP33 SER34 THR36 LEU41 TRP43 LYS84) |
| 8 | 13 | -0.47 | 6 | 11 | 1:(TYR57 ASN60 MET61 ASN62 THR63)3:(LYS80 PRO100 LEU101 LEU102 ASN103) |
| 9 | 16 | -0.48 | 10 | 14 | 3:(GLY49 GLY50 LEU51 VAL52 ASP90 TYR136 LYS162 ILE163 LEU164) |
| 10 | 13 | -0.48 | 6 | 15 | 1:(MET44 VAL45 HIS46 LYS133 TRP138) |
| 11 | 13 | -0.48 | 4 | 8 | 1:(PRO48 GLU52)3:(ILE31 VAL32 GLY58 ASP59 LYS60 LYS116) |
| 12 | 22 | -0.59 | 4 | 10 | 3:(MET83 VAL84 ASP85 GLN88 GLU97 ASN98 THR99) |
| 13 | 11 | -0.64 | 14 | 18 | 3:(LEU33 GLU34 ILE36 LYS44 LEU53 LEU61) |
| 14 | 7 | -0.68 | 9 | 12 | 1:(GLN157 VAL158 LEU160)3:(MET83 LEU127 GLU128 TYR135) |
| 15 | 11 | -0.71 | 6 | 15 | 1:(THR35 TYR57 ASP58 GLU59 ILE64 LYS99 ILE196 ALA197) |
| 16 | 10 | -0.73 | 3 | 11 | 3:(SER40 GLU77 TYR79 ASN142 GLY143 VAL155 ARG159) |
| 17 | 11 | -0.74 | 2 | 10 | 1:(GLU59 ASP152 GLU153 VAL164 MET165 ILE167 ASN168 THR169) |
| 18 | 10 | -0.77 | 4 | 8 | 1:(GLU40 GLU53)3:(LYS60 ASP62 THR114) |

**Figure S4K** (continued). Amino acid sequences of the binding pockets identified by SiteFinder for the EphB2-EphrinB2 dimer (1KGY).

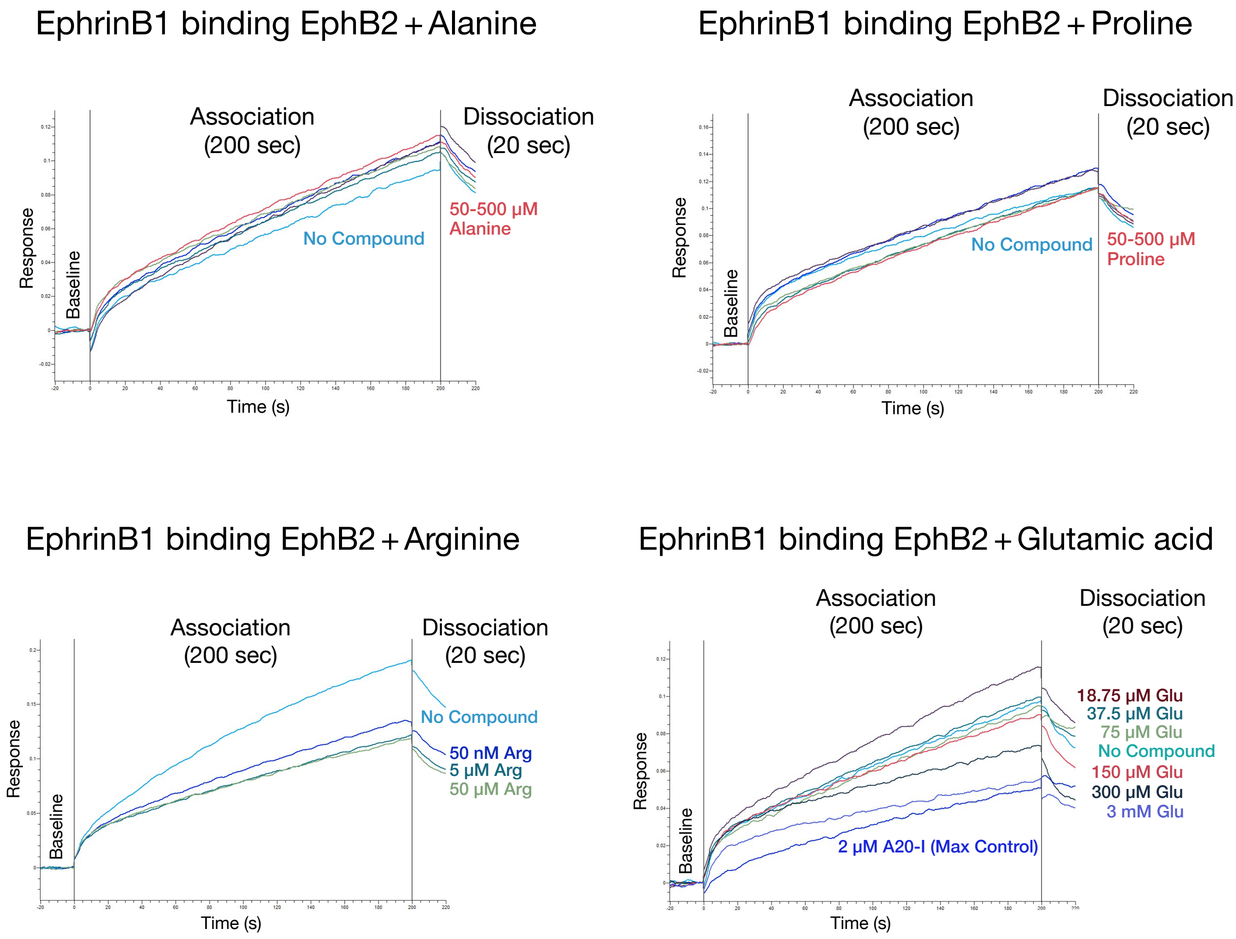


**Figure S5A.** BLI sensorgrams of immobilized human EphrinB1-Fc binding to 50 nM soluble human EphB2-His, testing the effects of 50 nM–500 µM excess L-alanine, L-proline, L-arginine, and L-glutamate added to the buffer conditions. Excess alanine and proline had little if any effect on the EphB2-EphrinB1 interaction, whereas arginine and glutamic acid led to concentration-dependent reductions in tetramer formation with little if any effect on dimer kinetics. The effect with Arginine was quite strong with as little as 50 nM exhibiting potent ability to reduce tetramer formation.

### Small EphB2-WT and EphB2-R79P proteins containing residues that form the dimer and tetramer interfaces

**Human EphB2-WT (19-210)** - His tag - Avi-tag

19
SinoSPVEETLMDSTTATAELGWMVHPPSGWEEVSGYDENMNTIRTYQVCNVFESSQN
NWLRTKFIRRRGAHRIHVEMKFSVRDCSSIPSVPGSCKETFNLYYYEADFDSATKTFPN
WMENPWVKVDTIAADESFSQVDLGGRVMKINTEVRSFGPVSRSGFYLAFQDYGGCM
SLIAVRVFYRK**S**PRIIQNGAIFQETGSHHHHHHHHHHGSGLNDIFEAQKIEWHE
197 210

**Human EphB2-R79P (19-210)** - His tag - Avi-tag

19
SinoSPVEETLMDSTTATAELGWMVHPPSGWEEVSGYDENMNTIRTYQVCNVFESSQN
NWLRTKFIPRRGAHRIHVEMKFSVRDCSSIPSVPGSCKETFNLYYYEADFDSATKTFPN
WMENPWVKVDTIAADESFSQVDLGGRVMKINTEVRSFGPVSRSGFYLAFQDYGGCM
SLIAVRVFYRK**S**PRIIQNGAIFQETGSHHHHHHHHHHGSGLNDIFEAQKIEWHE
197 210

The unpaired **Cysteine** at position 197 was changed to **Serine** to aid expression.

**Figure S6A.** Wild-type and R79P mutant versions of the N-terminal ligand-binding domain of EphB2 with indicated signal peptide and His/Avi tags were expressed and purified from 293 cells (work done by Sino Biological). Green highlighted residues are those involved in tetramerization with EphrinB2, with the proline substitution in R79P indicated in orange. As the unpaired cysteine residue at position 197 interfered with protein expression and is downstream of the region involved in ligand binding, it was changed to a serine in both proteins (blue). Note that Arg79 in human EphB2 corresponds to Arg87 in the mouse protein, which is the numbering used for crystal structures and the labeling of images and movies in the manuscript.

#### Sequence of small EphB2-WT and EphB2-R79P proteins confirmed using mass spectrometry

##### EphB2-WT (19-210)

| | |
|---|---|
| High | [F].ESSQNNW.[L] |
| High | [F].ESSQNNWLRTKF.[I] |
| High | [F].RRRGAHRIHVEM.[K] |
| High | [F].RRRGAHRIHVEMKF.[S] |
| High | [F].RRRGAHRIHVEMKF.[S] |
| High | [H].RIHVEMKF.[S] |

##### EphB2-R79P (19-210)

| | | |
|---|---|---|
| 37 | High | [F].ESSQNNWL.[R] |
| 38 | High | [F].ESSQNNWLRTKF.[I] |
| 39 | High | [W].LRTKFIPRRGAH.[R] |
| 40 | High | [F].PRRGAH.[R] |
| 41 | High | [F].PRRGAHRIHVEM.[K] |
| 42 | High | [F].PRRGAHRIHVEM.[K] |
| 43 | High | [F].PRRGAHRIHVEMKF.[S] |
| 44 | High | [F].PRRGAHRIHVEMKF.[S] |
| 45 | High | [H].RIHVEMKF.[S] |
| 46 | High | [H].RIHVEMKF.[S] |

**Figure S6B.** Mass spectrometry at the UT Southwestern Proteomics Core facility was used to verify the amino acid sequences of the human EphB2-WT and and EphB2-R79P proteins.

Silver stain confirms EphB2-WT and EphB2-R79P proteins

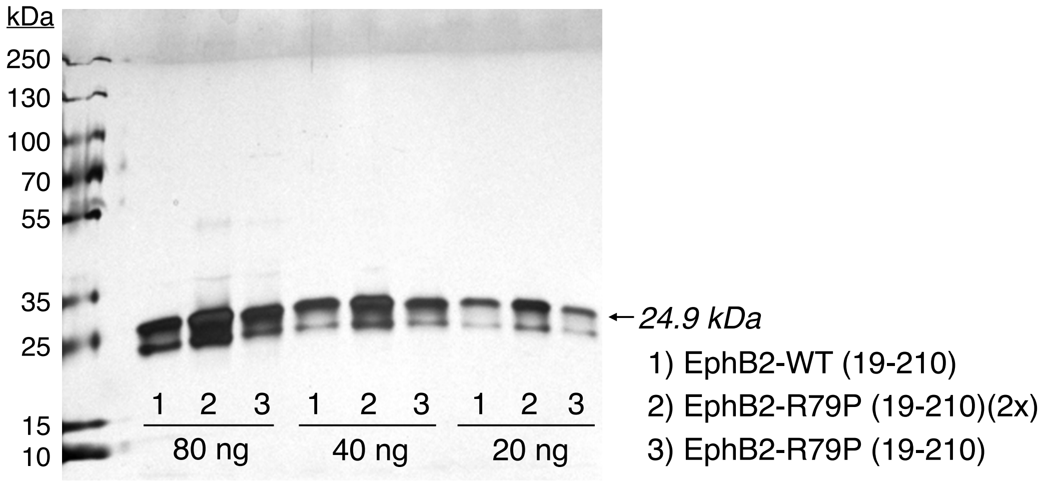


Silver Stained Gel

**Figure S6C.** Indicated amounts of human EphB2-WT and EphB2-R79P proteins were loaded and resolved on a 4-20% SDS-PAGE gradient gel, fixed, and then reacted with silver stain. The two proteins showed equal loading.

#### Immunoblot confirms EphB2-WT and EphB2-R79P proteins

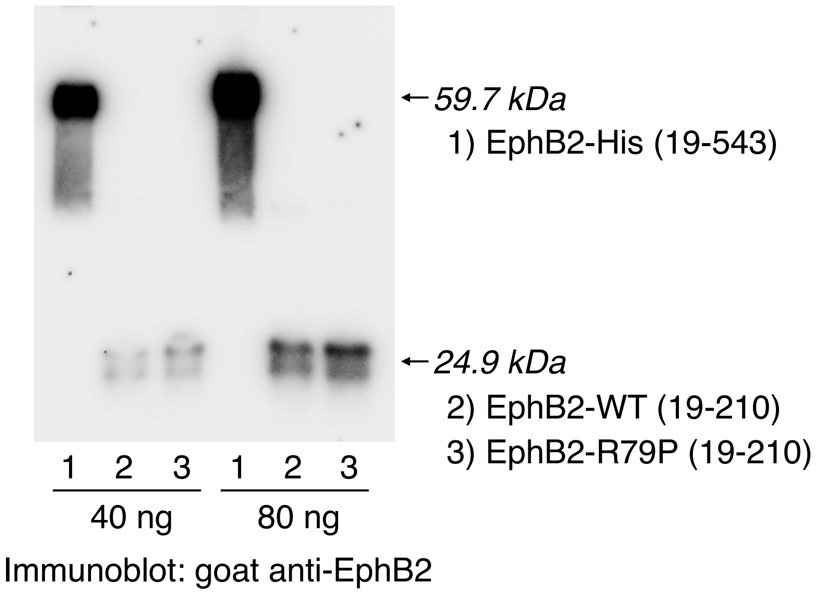


**Figure S6D.** Indicated amounts of human EphB2-His, EphB2-WT, and EphB2-R79P proteins were loaded and resolved on a 4-20% gradient SDS-PAGE gel, transferred to a Protran nitrocellulose membrane, and then immunoblotted with polyclonal goat anti-EphB2 antibodies raised against the receptor's full extracellular domain (R&D Systems). The antibody strongly recognized the larger EphB2-His protein containing the entire ectodomain (residues 19-210) and also detected the two smaller proteins, showing equal loading.

### EphrinB3 binding to EphB2-WT and EphB2-R79P proteins

EphB2-WT

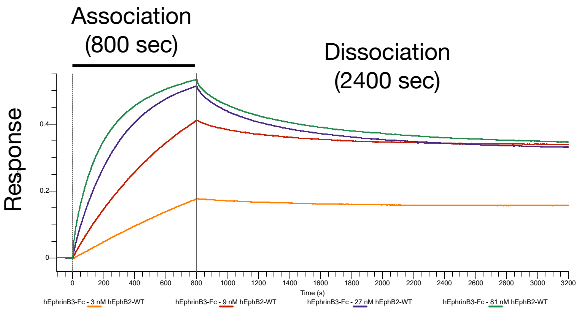


EphB2-R79P

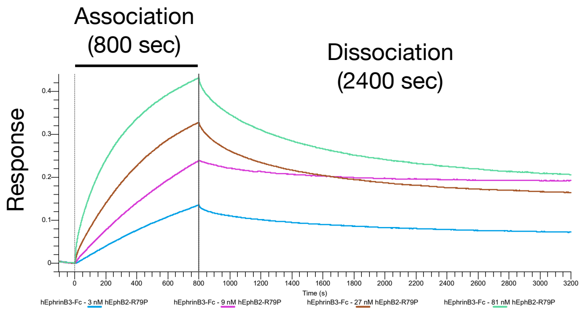


**EphrinB3 + EphB2-WT (19-210)**

$K_{D1}$ (tetramer) = 0.70 nM
$K_{a1}$ (tetramer) = 4.78E+04
$Kdis_1$ (tetramer) = 3.32E-05
$t_{1/2}$ (tetramer) = 0.24 days

$K_{D2}$ (dimer) = 7.77 nM
$K_{a2}$ (dimer) = 3.43E+05
$Kdis_2$ (dimer) = 2.66E-03
$t_{1/2}$ (tetramer) = 4.3 minutes

$R^2$ (association) = 0.9997
$R^2$ (dissociation) = 0.9993

**EphrinB3 + EphB2-R79P (19-210)**

$K_{D1}$ (tetramer) = 1.62 nM
$K_{a1}$ (tetramer) = 3.29E+04
$Kdis_1$ (tetramer) = 5.32E-05
$t_{1/2}$ (tetramer) = 0.15 days

$K_{D2}$ (dimer) = 4.87 nM
$K_{a2}$ (dimer) = 4.54E+05
$Kdis_2$ (dimer) = 2.21E-03
$t_{1/2}$ (tetramer) = 5.2 minutes

$R^2$ (association) = 0.9998
$R^2$ (dissociation) = 0.9984

**Figure S6E.** Octet global fit sensorgrams of 3, 9, 27, and 81 nM tests of EphB2-WT and EphB2-R79P proteins binding to AHC2 immobilized EphrinB3-Fc. Both proteins bound with high-affinity to immobilized EphrinB3, with dimer:tetramer global fit analysis indicating the mutant protein has a weaker affinity the tetramer interaction that is associated with a shortened half-life.

#### EphrinB3 binding to EphB2-WT and EphB2-R79P proteins

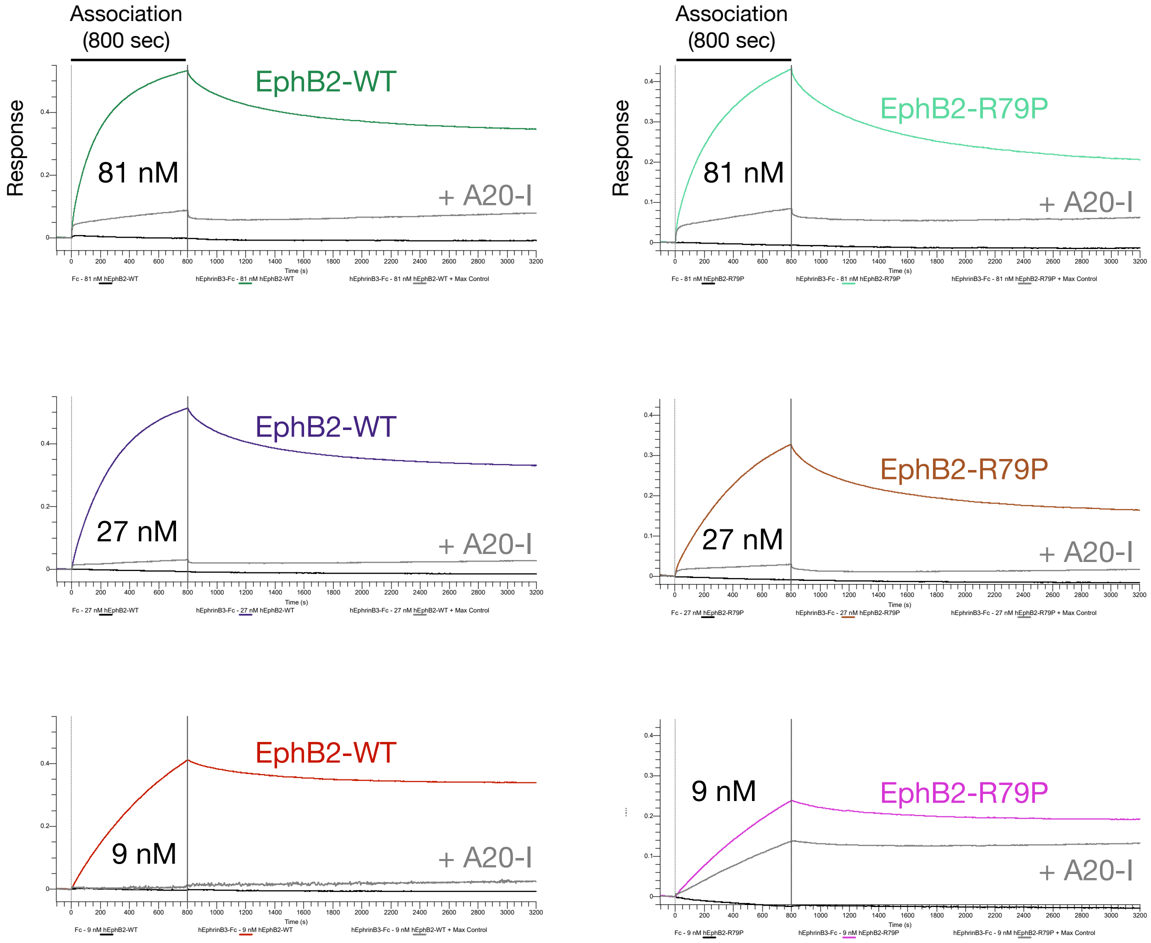


**Figure S6E** (continued)**.** The experiment above testing EphB2-WT and EphB2-R79P proteins binding to immobilized EphrinB3-Fc also examined if either protein showed non-specific binding to AHC2 sensor chips loaded with Fc only protein or whether the interaction with EphrinB3 was affected by addition of 2.43 μM A20-I to the binding buffer. The data showed neither protein bound to biosensors loaded with Fc only protein (black traces), while A20-I tetramer inhibitor strongly reduced binding of both proteins to EphrinB3 (gray traces).

### EphrinB1 binding to EphB2-WT and EphB2-R79P proteins

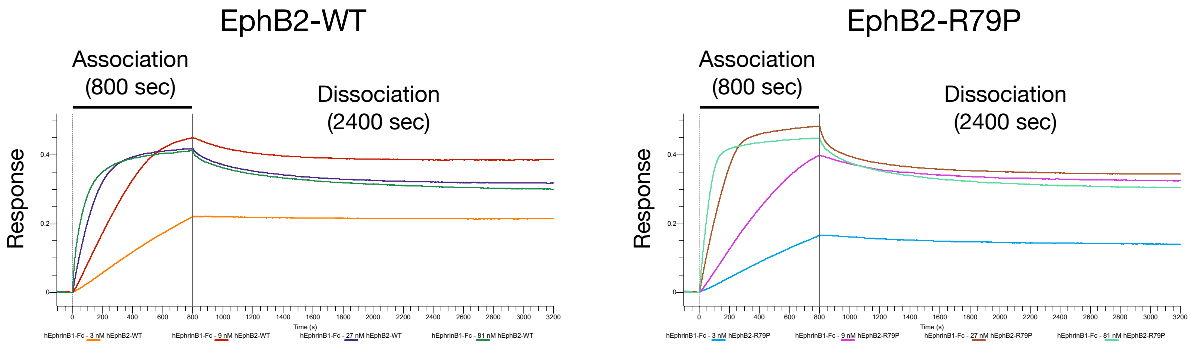


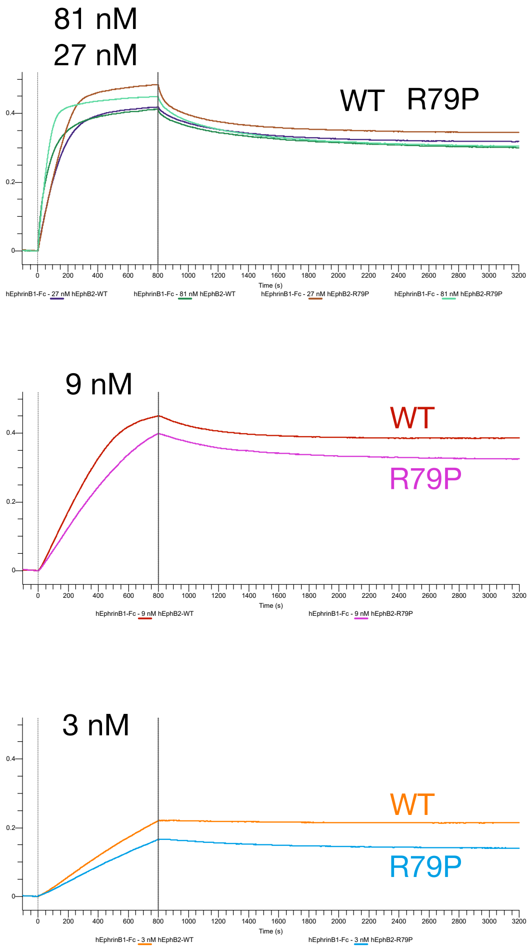


**EphrinB1 + EphB2-WT (19-210)**

**$K_{D1}$ (tetramer) = 0.19 nM**
**$K_{a1}$ (tetramer) = 6.16E+04**
**$Kdis_1$ (tetramer) = 1.19E-05**
**$t_{1/2}$ (tetramer) = 0.67 days**

**$K_{D2}$ (dimer) = 15.3 nM**
**$K_{a2}$ (dimer) = 2.54E+05**
**$Kdis_2$ (dimer) = 2.84E-03**
**$t_{1/2}$ (tetramer) = 4.1 minutes**

**$R^2$ (association) = 0.9973**
**$R^2$ (dissociation) = 0.9991**

**EphrinB1 + EphB2-R79P (19-210)**

**$K_{D1}$ (tetramer) = 2.03 nM**
**$K_{a1}$ (tetramer) = 1.17E+04**
**$Kdis_1$ (tetramer) = 2.34E-05**
**$t_{1/2}$ (tetramer) = 0.2 days**

**$K_{D2}$ (dimer) = 19.3 nM**
**$K_{a2}$ (dimer) = 1.35E+05**
**$Kdis_2$ (dimer) = 4.85E-03**
**$t_{1/2}$ (tetramer) = 2.4 minutes**

**$R^2$ (association) = 0.9975**
**$R^2$ (dissociation) = 0.9984**

**Figure S6F.** Octet global fit sensorgrams of 3, 9, 27, and 81 nM tests of EphB2-WT and EphB2-R79P proteins binding to AHC2 immobilized EphrinB1-Fc. Both proteins bound with high-affinity to immobilized EphrinB1, though the mutant showed reduced capability at 3 and 9 nM concentrations. Global fit analysis using our newly developed dimer:tetramer mathematical model indicated the mutant protein has a weaker affinity for both the dimer and tetramer, which was also reflected in a shortened half-life.

### EphrinB1 binding to EphB2-WT and EphB2-R79P proteins

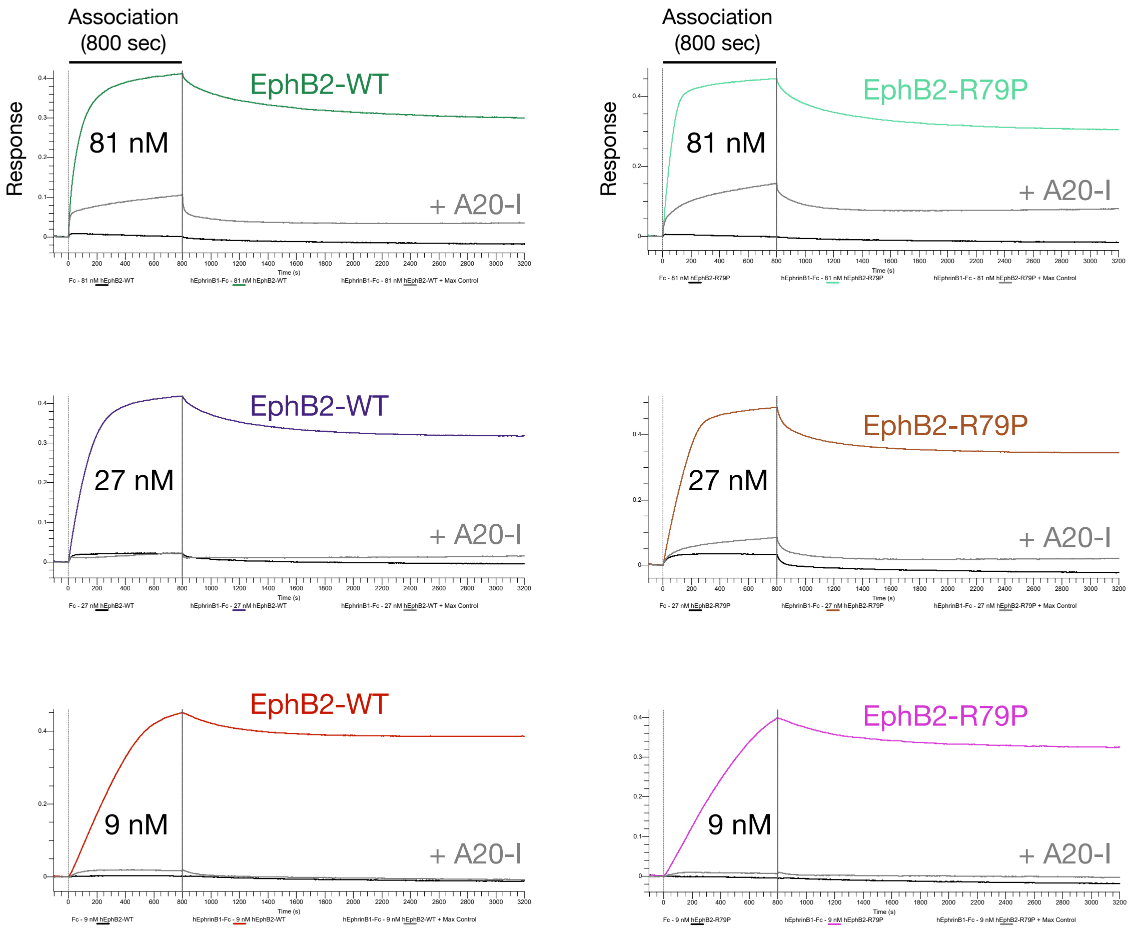


**Figure S6F** (continued)**.** The experiment above testing EphB2-WT and EphB2-R79P proteins binding to immobilized EphrinB1-Fc also examined if either protein showed non-specific binding to AHC2 sensor chips loaded with Fc only protein as a negative control or whether the interaction with EphrinB1 was affected by addition of 2.43 μM A20-I to the binding buffer. The data showed neither protein bound to biosensors loaded with Fc only protein (black traces), while A20-I tetramer inhibitor strongly reduced binding of both proteins to EphrinB1 (gray traces).

#### EphrinB2 binding to EphB2-WT and EphB2-R79P proteins

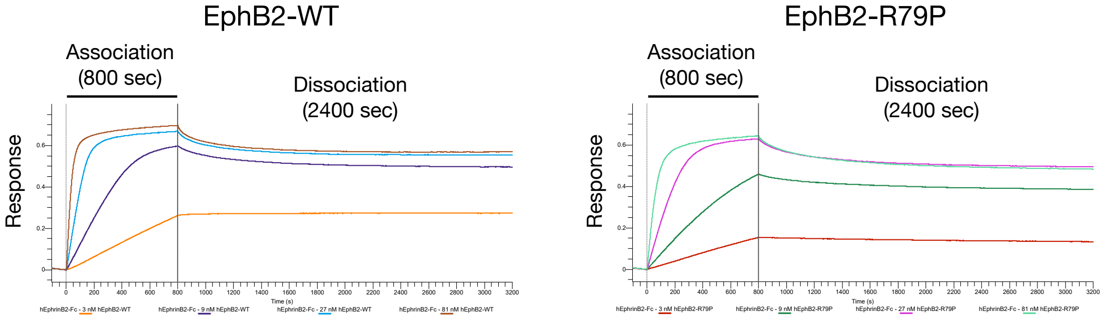


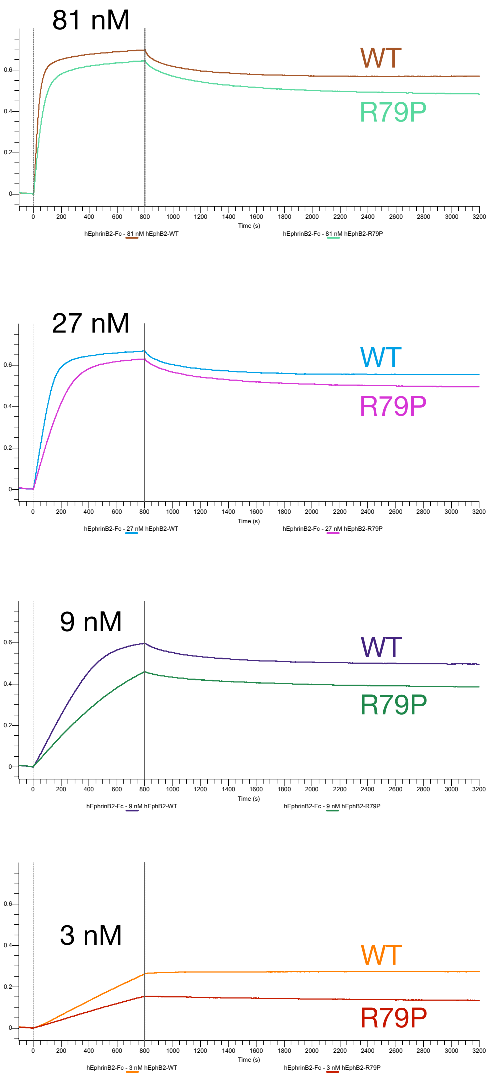


**EphrinB2 + EphB2-WT (19-210)**

$K_{D1}$ (tetramer) = 2.16 nM
$K_{a1}$ (tetramer) = 7.19E+03
$Kdis_1$ (tetramer) = 1.56E-05
$t_{1/2}$ (tetramer) = 0.52 days

$K_{D2}$ (dimer) = 17.80 nM
$K_{a2}$ (dimer) = 2.70E+05
$Kdis_2$ (dimer) = 4.80E-03
$t_{1/2}$ (tetramer) = 2.4 minutes

$R^2$ (association) = 0.9935
$R^2$ (dissociation) = 0.9952

**EphrinB2 + EphB2-R79P (19-210)**

$K_{D1}$ (tetramer) = 5.64 nM
$K_{a1}$ (tetramer) = 1.51E+03
$Kdis_1$ (tetramer) = 2.34E-05
$t_{1/2}$ (tetramer) = 0.24 days

$K_{D2}$ (dimer) = 21.74 nM
$K_{a2}$ (dimer) = 1.35E+05
$Kdis_2$ (dimer) = 4.85E-03
$t_{1/2}$ (tetramer) = 2.9 minutes

$R^2$ (association) = 0.9986
$R^2$ (dissociation) = 0.9998

**Figure S6G.** Octet global fit sensorgrams of 3, 9, 27, and 81 nM tests of EphB2-WT and EphB2-R79P proteins binding to AHC2 immobilized EphrinB2-Fc. Both proteins bound with high-affinity to immobilized EphrinB2, though the mutant showed reduced capability at all four concentrations. Dimer:tetramer global fit analysis indicated the mutant protein has a weaker affinity for both the dimer and tetramer interactions, with the tetramer interaction also showing a shortened half-life.

#### EphrinB2 binding to EphB2-WT and EphB2-R79P proteins

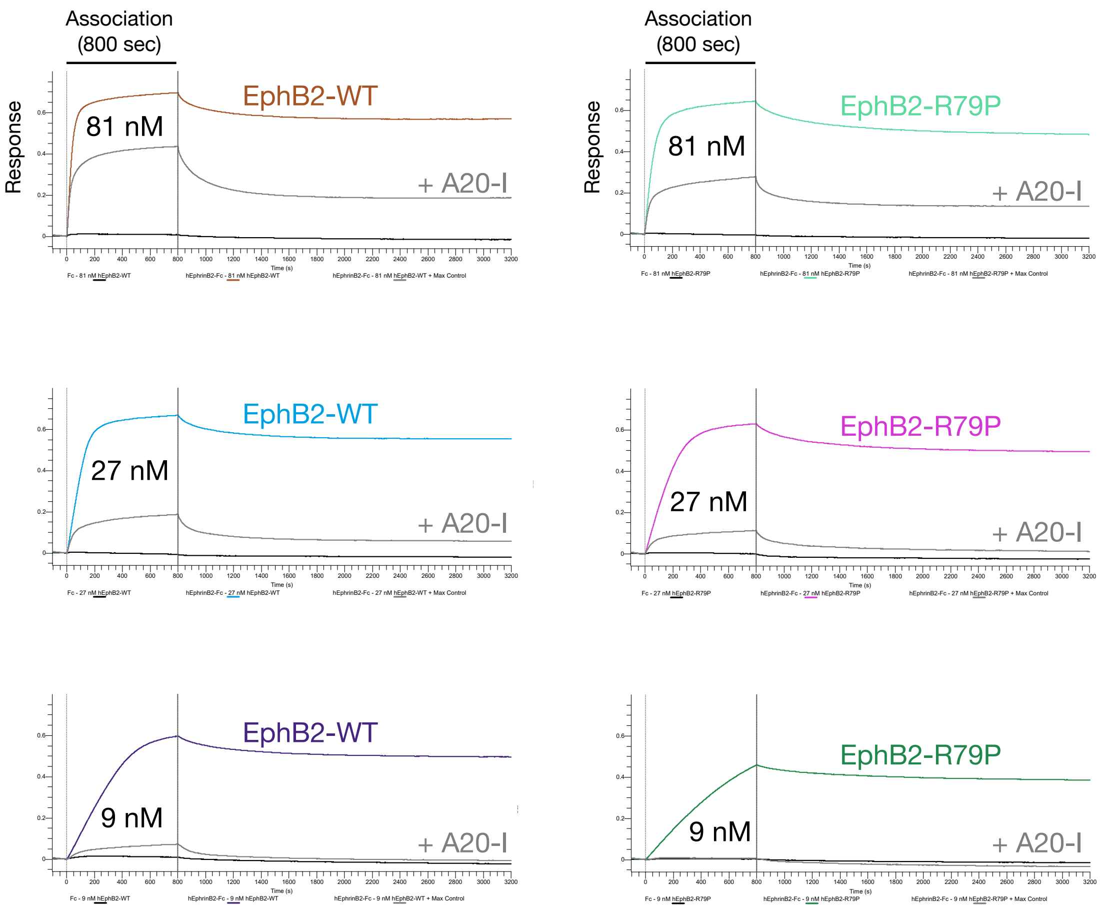


**Figure S6G** (continued)**.** The experiment above testing EphB2-WT and EphB2-R79P proteins binding to immobilized EphrinB2-Fc also examined if either protein showed non-specific binding to AHC2 sensor chips loaded with Fc only protein or whether the interaction with EphrinB2 was affected by addition of 2.43 μM A20-I to the binding buffer. The data showed neither protein bound to biosensors loaded with Fc only protein (black traces), while A20-I tetramer inhibitor strongly reduced binding of both proteins to EphrinB2 (gray traces).

#### EphB2-WT vs EphB2-R79P [2/2]

Student's t-test, two-tailed, equal variance, within one ligand x buffer x protocol (association / dissociation length); buffers and protocols are never pooled.
n = runs with a value for that constant; p only where both variants have n >= 2.

**EphrinB3-Fc, PBST, 90-min association / 15 h dissociation: WT = 2026.101, 2026.103, 2026.108; R79P = 2026.106, 2026.107, 2026.109**

| Constant | WT mean ± SD | R79P mean ± SD | R79P / WT | Student's p |
|---|---|---|---|---|
| Ka1 (1/Ms) | 8.163e+03 ± 1.265e+03 (n=3) | 8.433e+03 ± 1.233e+03 (n=3) | 1.03 | 0.805 |
| Kdis1 (1/s) | 1.038e-06 ± 3.521e-07 (n=3) | 3.010e-06 ± 1.033e-06 (n=3) | 2.90 | 0.035 |
| KD1 (nM) | 0.13 ± 0.0484 (n=3) | 0.36 ± 0.117 (n=3) | 2.77 | 0.035 |
| Ka2 (1/Ms) | 1.463e+05 ± 2.418e+04 (n=3) | 1.410e+05 ± 4.925e+04 (n=3) | 0.96 | 0.876 |
| Kdis2 (1/s) | 5.178e-04 ± 6.677e-05 (n=3) | 1.362e-03 ± 2.996e-04 (n=3) | 2.63 | 0.009 |
| KD2 (nM) | 3.58 ± 0.586 (n=3) | 10.2 ± 2.76 (n=3) | 2.85 | 0.015 |

#### EphB2-WT vs EphB2-R79P: half-lives

Student's t-test, two-tailed, equal variance, on the per-run half-lives ($t_{1/2}$ = ln 2 / Kdis), within one ligand x buffer x protocol (association / dissociation length);
buffers and protocols are never pooled. n = runs.

**EphrinB3-Fc, PBST, 90-min association / 15 h dissociation: WT = 2026.101, 2026.103, 2026.108; R79P = 2026.106, 2026.107, 2026.109**

| Quantity | WT mean ± SD | R79P mean ± SD | R79P / WT | Student's p |
|---|---|---|---|---|
| Tetramer half-life (days) | 8.46 ± 3.29 (n=3) | 2.85 ± 0.821 (n=3) | 0.34 | 0.046 |
| Dimer half-life (min) | 22.6 ± 3.14 (n=3) | 8.73 ± 1.73 (n=3) | 0.39 | 0.003 |

**Figure S6H.** Kinetic results of EphB2-WT and EphB2-R79P mutant proteins binding to immobilized EphrinB3-Fc; three independent experiments were assessed for each interaction.

### EphrinA5 binding to EphB2-WT and EphB2-R79P proteins

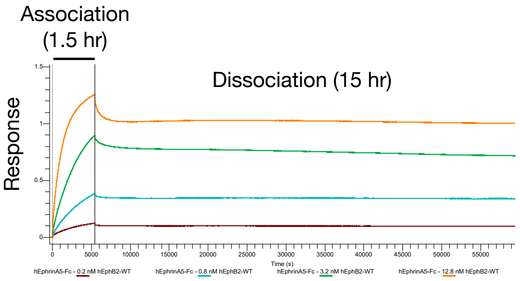


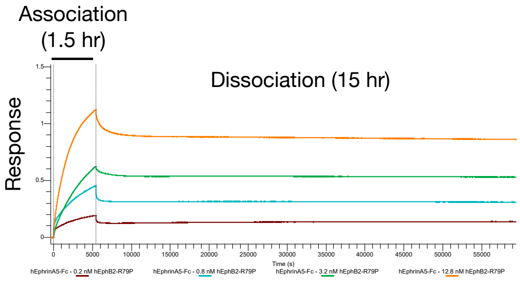


**EphrinA5 + EphB2-WT (19-210)**

$K_{D1}$ (tetramer) = 0.05 nM
$K_{a1}$ (tetramer) = 1.36E+04
$Kdis_1$ (tetramer) = 6.65E-07
$t_{1/2}$ (tetramer) = 12.1 days

$K_{D2}$ (dimer) = 6.12 nM
$K_{a2}$ (dimer) = 3.60E+05
$Kdis_2$ (dimer) = 2.21E-03
$t_{1/2}$ (tetramer) = 5.2 minutes

$R^2$ (association) = 0.9989
$R^2$ (dissociation) = 0.9990

**EphrinA5 + EphB2-R79P (19-210)**

$K_{D1}$ (tetramer) = 0.23 nM
$K_{a1}$ (tetramer) = 2.63E+03
$Kdis_1$ (tetramer) = 5.90E-07
$t_{1/2}$ (tetramer) = 13.6 days

$K_{D2}$ (dimer) = 37.88 nM
$K_{a2}$ (dimer) = 6.67E+04
$Kdis_2$ (dimer) = 2.23E-03
$t_{1/2}$ (tetramer) = 4.6 minutes

$R^2$ (association) = 0.9987
$R^2$ (dissociation) = 0.9992

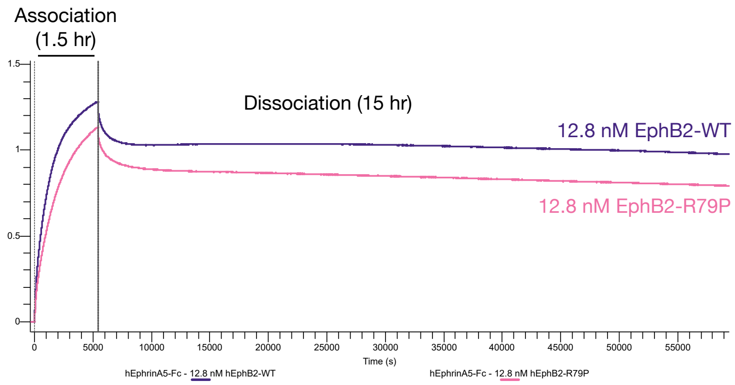


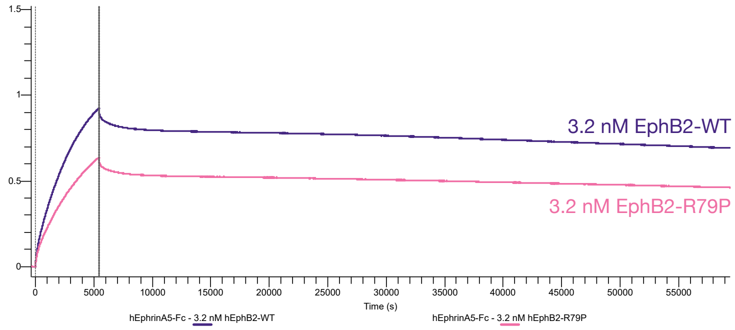


**Figure S6I.** Octet global fit sensorgrams of 0.2, 0.8, 3.2, and 12.8 nM tests of EphB2-WT and EphB2-R79P proteins binding to AHC2 immobilized EphrinA5-Fc. Both proteins bound with high-affinity to immobilized EphrinA5, though the mutant showed reduced capability as shown for the higher 3.2 and 12.8 nM concentration runs. Dimer:tetramer global fit analysis indicated the mutant protein has a weaker affinity for both the dimer and tetramer interactions.

### Supplementary Table S1: Copper chelation assay testing FDA drugs

| # | Chemical Structure | Common Nomenclature (if applicable) | Molecular weight (g/mol) | Octet $IC_{50}$ data EphrinB2-EphB2 (Absolute) | Cu chelator? | *In vitro* copper chelation assay |
|---|---|---|---|---|---|---|
| 1 | 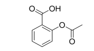 | Aspirin | 180.16 | No Activity (up to 100 μM) | Yes | 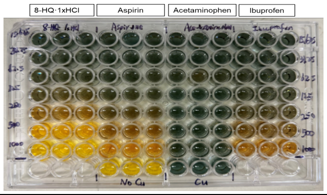 |
| 2 | 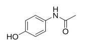 | Acetaminophen | 151.16 | No Activity (up to 100 μM) | No | |
| 3 | 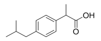 | Ibuprofen | 206.29 | No Activity (up to 100 μM) | Yes | |
| 4 | 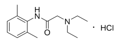 | Lidocaine hydrochloride | 288.81 | No Activity (up to 100 μM) | No | 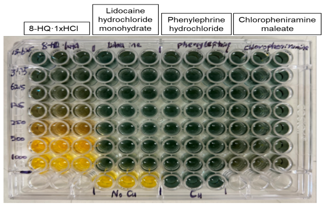 |
| 5 | 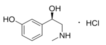 | Phenylephrine hydrochloride | 203.67 | No Activity (up to 100 μM) | No | |
| 6 | 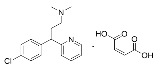 | Chloropheniramine maleate | 390.86 | No Activity (up to 100 μM) | No | |
| 7 | 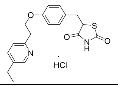 | Pioglitazone hydrochloride | 392.9 | No Activity (up to 100 μM) | Yes | 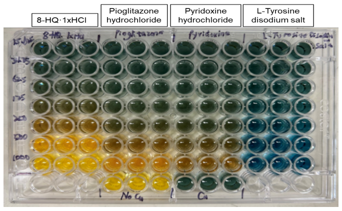 |
| 8 | 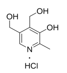 | Pyridoxine hydrochloride | 205.64 | No Activity (up to 100 μM) | Yes | |
| 10 | 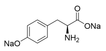 | L-Tyrosine disodium salt | 225.2 | No Activity (up to 100 μM) | No | |
| 9 | 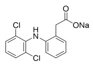 | Diclofenac sodium salt | 318.13 | No Activity (up to 100 μM) | No | 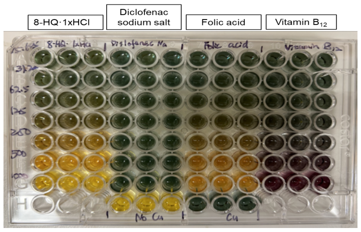 |
| 11 | 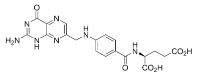 | Folic acid | 441.4 | No Activity (up to 100 μM) | Yes | |
| 12 | 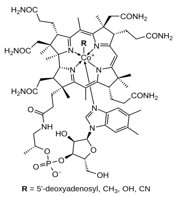 | Vitamin $B_{12}$ | 1,355.38 | No Activity (up to 100 μM) | No | |

### Supplementary Chemistry Information

#### Materials and Methods

All chemicals for synthesis were obtained from commercial suppliers (Thermo Scientific, Sigma-Aldrich, or Ambeed) in “reagent grade” purity or better and used as received. Deuterated solvents used for NMR experiments were purchased from Cambridge Isotope Laboratories.

For purification of reaction products by flash column chromatography, silica gel from Sigma-Aldrich (Silica gel, high-purity grade, pore size 60 Å, 220-440 mesh particle size, 35-75 µm particle size) was used. Thin layer chromatography was performed using aluminum sheets pre-coated with silica gel 60 matrix and fluorescent indicator for 254 nm, purchased from Millipore Sigma. Visualization of the compounds occurred under UV light of 254 nm wavelength.

NMR spectra were recorded on a Bruker Avance NEO instrument operating at 400 proton frequency. Samples were recorded at 295 k in 5 mm diameter tubes. Coupling patterns are abbreviated with s (singlet), d (doublet), dd (doublet of doublets), ddd (doublet of doublet of doublets), t (triplet), hept (heptet) and m (multiplet) with indicated coupling constants J in Hertz (Hz).

High-resolution mass spectra (HRMS) were recorded by Dr. Hamid Baniasadi (UT Southwestern Medical Center, Metabolomics Core) on Sciex TripleTOF® 6600 System. The data was processed by Sciex OS software.

#### Synthetic Schemes

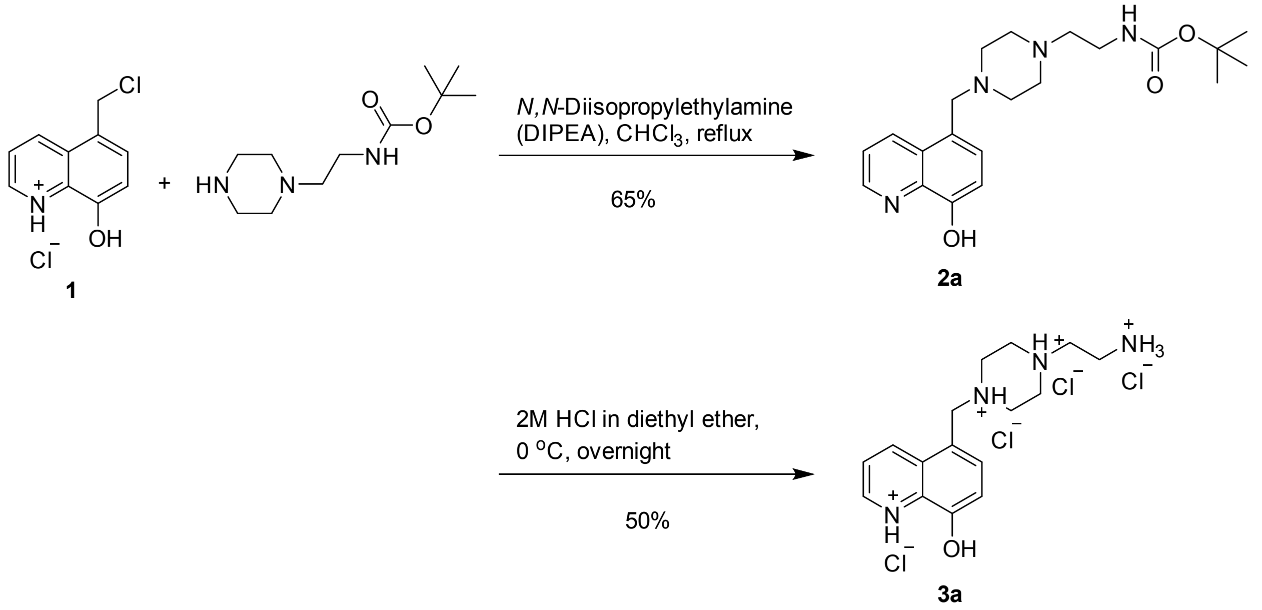


Synthetic route to primary amine hydrochloride salt.

#### Synthetic Procedures

Compound 1 was prepared according to published procedures with modification.[1]

### 2a

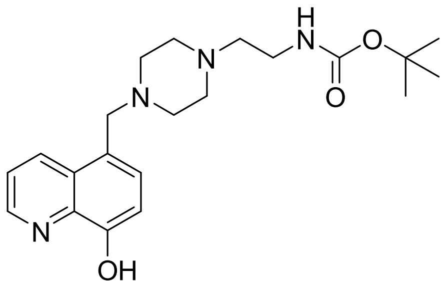

To a 100 mL round bottom flask, compound **1** (2006.7 mg, 8.7214 mmol), 1-(2-*N*-Boc-aminoethyl)piperazine (2000 mg, 8.7214 mmol), *N,N*-diisopropylethylamine (DIPEA) (4.558 mL, 26.1667 mmol) and 40 mL of chloroform were added. The reaction mixture was refluxed overnight. Following completion, the organic solvent was removed under reduced pressure. The crude product was purified with column chromatography on silica gel (dichloromethane:methanol, 200:1 to 10:1, v:v). The product was obtained as a yellowish viscous oil in 65% yield (**2a**, 2191.0 mg)

**$^1$H NMR** (400 MHz, $CDCl_3$) δ 8.74 (dd, *J* = 4.2, 1.6 Hz, 1H), 8.62 (dd, *J* = 8.5, 1.6 Hz, 1H), 7.41 (dd, *J* = 8.5, 4.2 Hz, 1H), 7.29 (d, *J* = 7.7 Hz, 1H), 7.04 (d, *J* = 7.6 Hz, 1H), 5.09 (s, 1H), 3.75 (s, 2H), 3.18 (t, *J* = 6.2 Hz, 2H), 2.63 – 2.12 (m, 10H), 1.42 (s, 9H).

**$^{13}$C NMR** (101 MHz, $CDCl_3$) δ 156.0, 151.9, 147.5, 138.7, 134.1, 128.9, 127.9, 124.6, 121.4, 108.8, 79.1, 60.6, 57.1, 53.0, 52.9, 37.2, 28.5.

**HRMS** (ESI) m/z: [M+H$^+$] Calculated for C21H31N4O3+ 387.23907; Found 387.23923.

### 3a

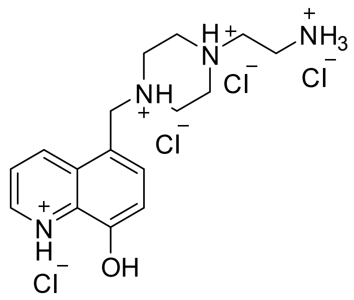

In a 50 mL round bottom flask, compound **2a** (100.0 mg, 0.2587 mmol) was dissolved in 4 mL dichloromethane. After cooling down to 0 $^{o}$C, 2 M HCl in diethyl ether (8.0 mL, 16 mmol) was added. The reaction mixture was stirred overnight. Following completion, the mixture was filtered and washed with chloroform and cold methanol under vacuum. The solid was then transferred to a scintillation vial and kept under vacuum to give dried primary amine hydrochloride salt in 50% yield (**3a**, 56.0 mg).

**$^1$H NMR** (400 MHz, $D_2O$) δ 9.38 (d, *J* = 8.8 Hz, 1H), 9.06 (d, *J* = 5.3 Hz, 1H), 8.18 (dd, *J* = 8.8, 5.4 Hz, 1H), 7.95 (d, *J* = 8.2 Hz, 1H), 7.42 (d, *J* = 8.1 Hz, 1H), 4.96 (s, 2H), 3.71 (t, *J* = 5.3 Hz, 4H), 3.63 – 3.47 (m, 4H), 3.47 – 3.36 (m, 4H).

**$^{13}$C NMR** (101 MHz, $D_2O$) δ 149.62, 143.16, 143.12, 136.64, 129.47, 129.16, 123.05, 115.55, 115.52, 55.45, 52.72, 49.28, 49.13, 34.09.

**HRMS** (ESI) m/z: [M+H$^+$] Calculated for C16H23N4O+ 287.18664; Found 287.18716.

### NMR Spectra

## 2a

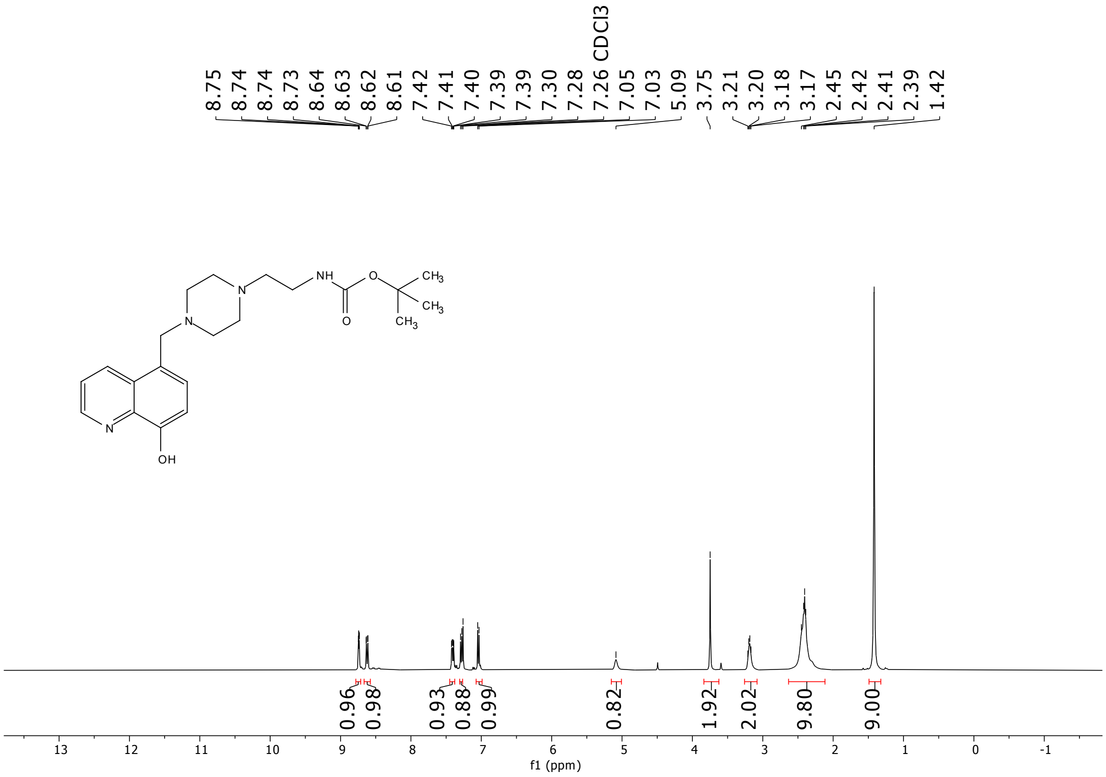


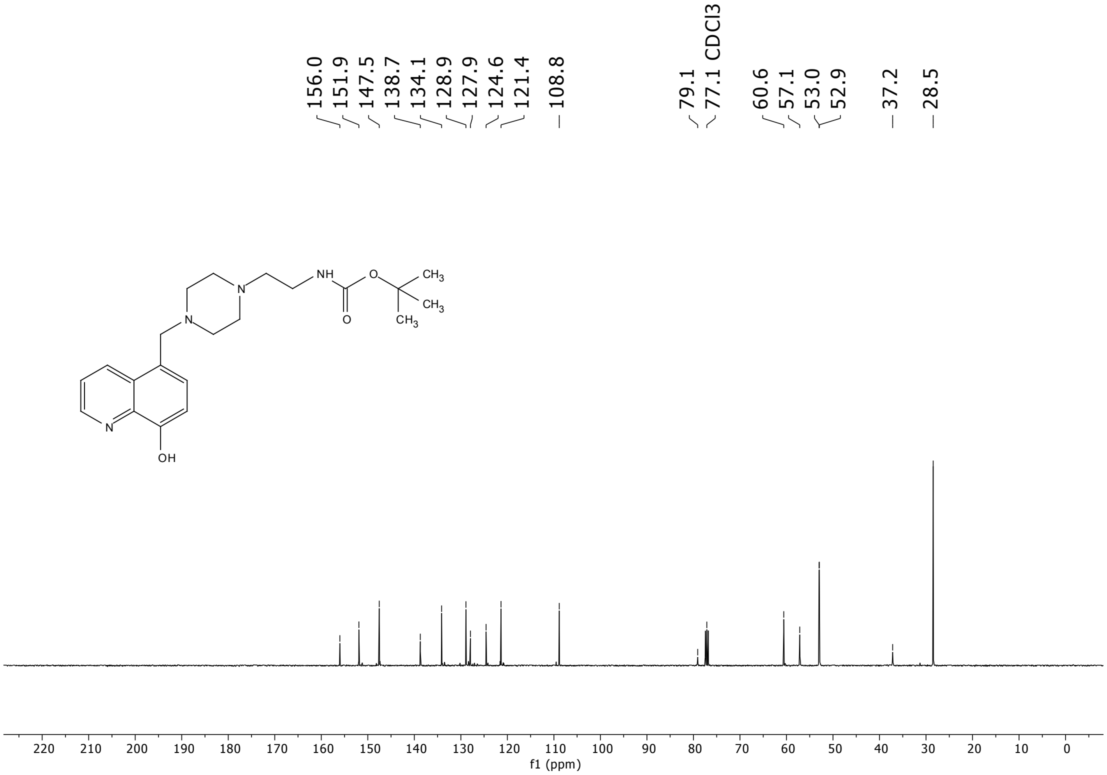

## 3a

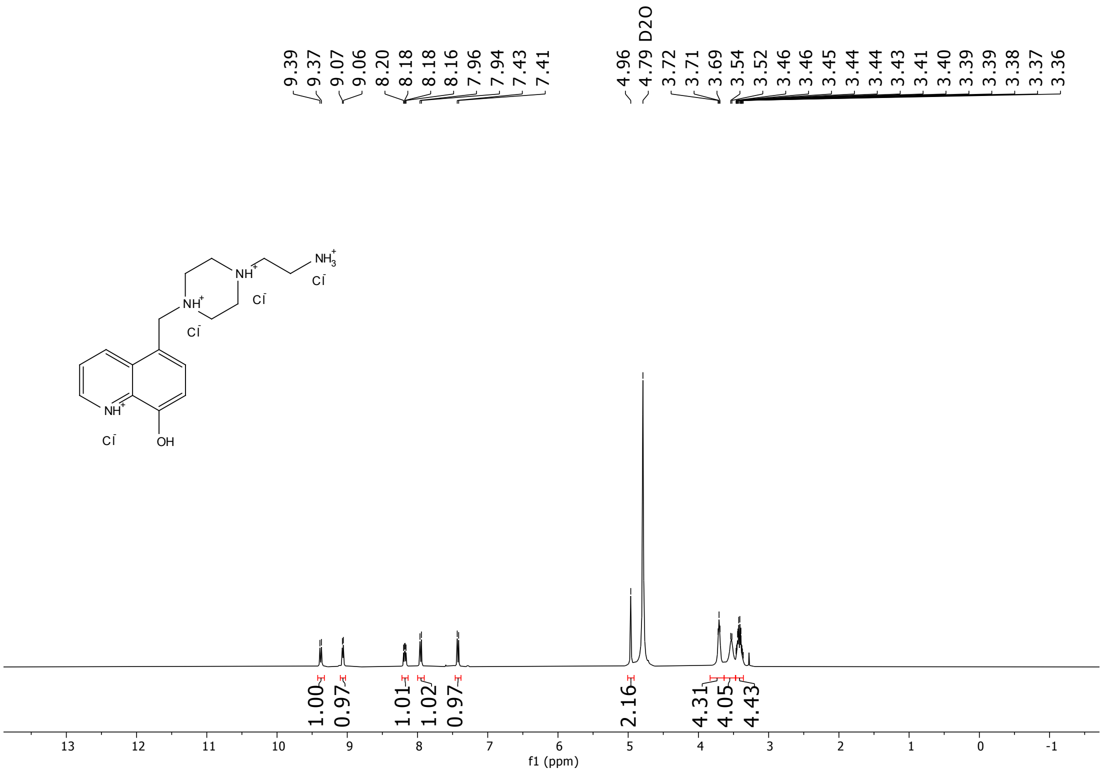
9.39
9.37
9.07
9.06
8.20
8.18
8.18
8.16
7.96
7.94
7.43
7.41
4.96
4.79 D2O
3.72
3.71
3.69
3.54
3.52
3.46
3.46
3.45
3.44
3.44
3.43
3.41
3.40
3.39
3.39
3.38
3.37
3.36
1.00
0.97
1.01
1.02
0.97
2.16
4.31
4.05
4.43
13
12
11
10
9
8
7
6
5
4
3
2
1
0
-1
f1 (ppm)

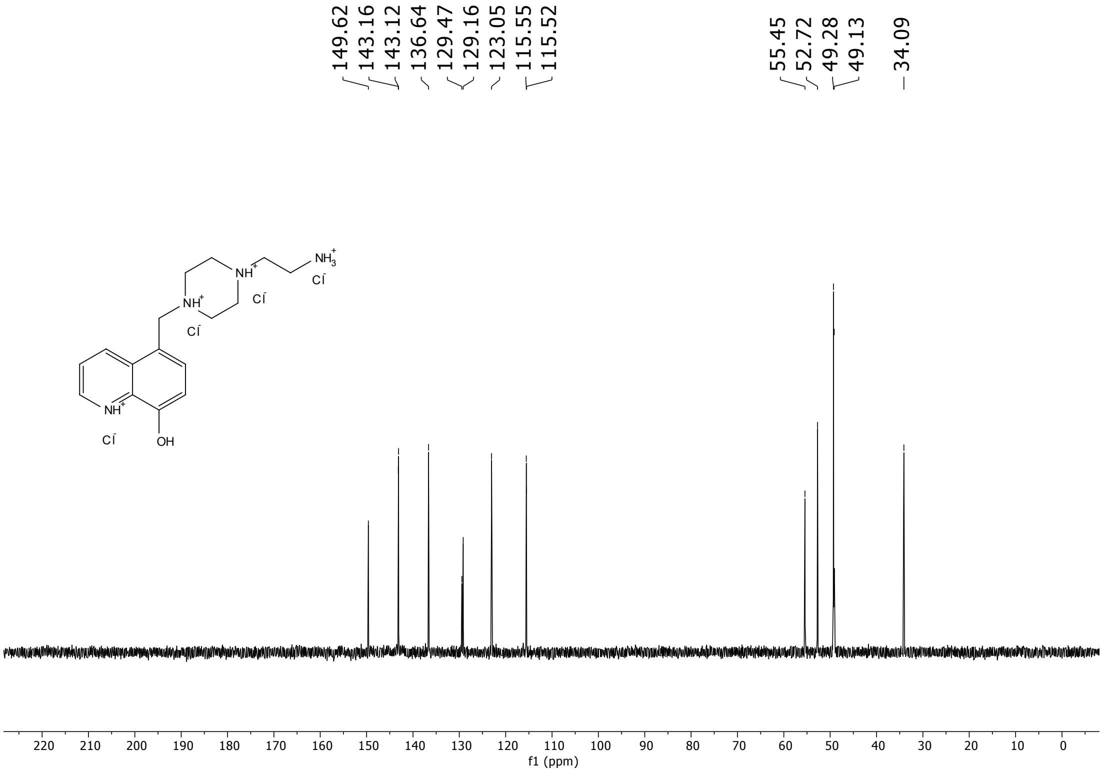
149.62
143.16
143.12
136.64
129.47
129.16
123.05
115.55
115.52
55.45
52.72
49.28
49.13
34.09
220
210
200
190
180
170
160
150
140
130
120
110
100
90
80
70
60
50
40
30
20
10
0
f1 (ppm)

### HRMS Spectra

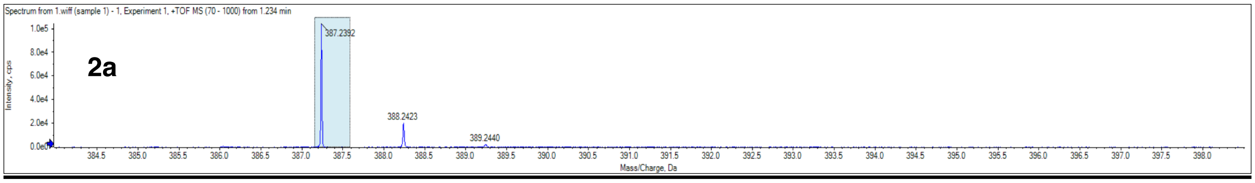


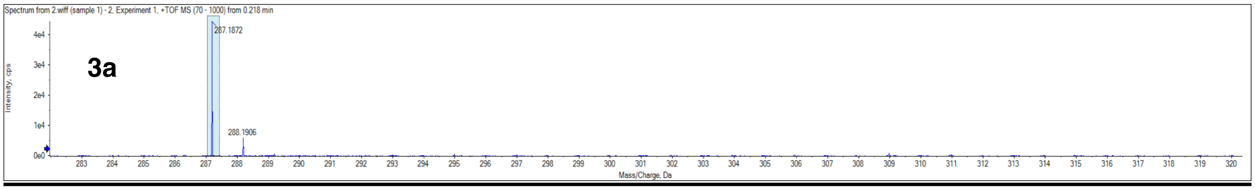

### Legends for Supplementary Videos

Title: Proteinaceous Metal-Binding Eph-Ephrin Tetramerization Is Modulated by Copper and Chelators

By: Khambete, Wang, Henkemeyer, & Henkemeyer

High resolution video files are available to download:

https://figshare.com/s/e34b45ec8d159e00a75b

Video S1 (Related to Figure 1A): Formation of EphB2-EphrinB2 circular tetramer. Movie using ribbon diagrams of the co-crystal structure illustrates how EphB2 (blue) and EphrinB2 (red) ectodomains first come together to form the dimer, two of which can then precisely fit together to form the circular tetramer.

Video S2 (Related to Figure 2B): EphB2 tetramer interface amino acids highlighted in a ribbon diagram of the unbound monomer crystal structure (1NUK). Amino acids identified include the electropositive arginine residues (orange) and other residues involved in making tetramer contacts (magenta).

Video S3 (Related to Figure 2B): EphrinB2 tetramer amino acids highlighted in a ribbon diagram of the unbound monomer crystal structure (1IKO). Amino acids identified include the key aspartic acid residue in the C-D loop (light blue) and other residues involved in making tetramer contacts (yellow).

Video S4 (Related to Figure 2C): EphB2 (blue, space-filled) - EphrinB2 (red, ribbon) tetramer interface. The EphB2-EphrinB2 co-crystal (1KGY) highlighting amino acids of each protein involved in forming the tetramer interface, with EphrinB2 residues identified. Green lines indicate hydrogen bonds with larger cylindrical length corresponding to increasing hydrogen bonding strength. Coloring is same as above.

Video S5 (Related to Figure 2C): EphB2 (blue, ribbon) - EphrinB2 (red, space-filled) tetramer interface. The EphB2-EphrinB2 co-crystal (1KGY) highlighting amino acids of each protein involved in forming the tetramer interface, with EphB2 residues identified. Coloring is same as above.

Video S6 (Related to Figure 2D): EphB2 (blue, ribbon) - EphrinB2 (red, ribbon) tetramer interface interactions focusing on key electrostatic interactions between Arg87 of EphB2 and Asp69 of EphrinB2, and other hydrogen bonds that form the interface. Ionic bonding interactions are not explicitly shown. Coloring is same as above.

Video S7 (Related to Figure 2D): EphB2 (blue, ribbon) - EphrinB2 (red, ribbon) tetramer interface interactions focusing on the receptor-ligand aromatic π-π stacking interaction between Phe128-Tyr37 and π interaction between Phe135-Glu34. Phe128 in EphB2 shows a large conformation change from monomer to tetramer. Phe128-Try37 and Phe135-Glu34 interactions are the second and fourth highest in energy within the tetramer interface, respectively (Extended Data Fig. 4d). Coloring is same as above.

Video S8 (Related to Figure 2I): Shows how EGTA (yellow) may dock into the EphB2 tetramer interface via interactions with both Arg87 and Arg89 (orange). Negative charged oxygens in the carboxylates of EGTA (red) tend to bind strongly to positive amino acids like Arg87 and Arg89. The electrostatic interaction map is shown on the surface of EphB2 where blue represents electropositive spaces and red represents electronegative spaces. This docking result represents the third highest docking score in the results section (Extended Data Fig. 2h).

Video S9 (Related to Figure 2I): Shows how 3511-0013 (yellow) may dock into the EphB2 tetramer interface via interactions with Arg89 (orange). The negative charged oxygen in the 8-HQ ring of 3511-0013 (red) binds to positive amino acid Arg89. This docking result represents the highest docking score in the results section (Extended Data Fig. 2h). Coloring is same as above.

Video S10 (Related to Figure 2I): Shows how A20 (yellow) may dock into the EphB2 tetramer interface via interactions with Arg 87 and Arg89 (orange). The negative charged oxygen in the 8-HQ ring of A20 (red) binds to positive amino acid Arg89, while the aromatic hydroxyquinoline has π interaction with Arg87. This docking result represents the highest docking score in the results section (Extended Data Fig. 2h). Coloring is same as above.

Video S11 (Related to Figure 2I): Additional movie showing how A20 (green) may dock into the EphB2 tetramer interface via interactions with Arg89. The negative charged oxygen in the 8-HQ ring of A20 (red) tends to bind strongly (purple line) to positive amino acid Arg89. This docking result was obtained using Schrödinger computational tools Glide and Maestro and are similar to that obtained using MOE docking.

Video S12 (Related to Figure 2I): NAMD simulation of EGTA interacting with EphB2 for 30 nanoseconds. EGTA docked to EphB2 is very stable and remained bound to the receptor for the entire 30 nanoseconds, likely because this multidentate chelator uses numerous oxygens (red) to interact with both the Arg89 and Arg87 residues within the protein's tetramer interface. Electrostatic interactions are indicated in green.

Video S13 (Related to Figure 2I): NAMD simulation of 3511-0013 interacting with EphB2 for 30 nanoseconds. 3511-0013 docked to Arg89 of EphB2 mainly through its oxygen (red) at 8-HQ position 8 and nitrogen (blue) at position 1, as expected for a bidentate chelator. The interaction of 3511-0013 with EphB2 is less stable than EGTA as it dissociated from the receptor after ~20 nanoseconds, likely because, as a bidentate chelator, it is only able to interact with one of the electropositive arginine residues within the receptor's tetramerization interface. Electrostatic interactions are indicated in green.

Video S14 (Related to Figure 3B): Shows the copper binding site within the EphrinB2 C-D loop. MOE energy minimization and SiteFinder analysis identified a copper-binding site within the C–D loop of the EphrinB2 tetramer interface, stabilized by an ionic interaction between $Cu^{2+}$ and Asp69, as well as coordinate-covalent bonds between $Cu^{2+}$ and both Lys71 and Thr72. Coloring is same as above.

Video S15 (Related to Figure 3B): NAMD simulation of EphrinB2 binding to copper for 30 nanoseconds. Copper was docked into the EphrinB2 structure using manual placement and energy minimization in MOE. The metal was observed to remain stably bound to EphrinB2 for the entire 30 nanosecond simulation, interacting with Asp69, Lys71, and Thr72 residues of the ligand C-D loop involved in tetramerization.

Video S16 (Related to Figure 4A): NAMD simulation of the EphB2-EphrinB2 dimerization interface for 30 nanoseconds.

Video S17 (Related to Figure 4A): NAMD simulation of the EphB2-EphrinB2 tetramerization interface for 30 nanoseconds. For this simulation, two EphB2-EphrinB2 dimers were juxtaposed to study the molecular dynamics of the two tetramer interface interactions as they form the circular tetramer.

Video S18 (Related to Figure 4C): The EphB2-EphrinB2 tetramer with electrostatic map shows the tetramerization interface is guided by electrostatic interactions. Coloring is same as above.

Video S19 (Related to Figure 4D): The EphB2-EphrinB2 dimer with hydrophobicity map shows the dimerization interface is driven by hydrophobic interactions. Coloring is same as above.

Video S20 (Related to Figure 4): Superimposition of EphrinB2 monomer (1IKO, green) and tetramer (1KGY, yellow and light blue) crystal structures highlighting changes in amino acid positioning of key residues that form the tetramer interface.

Video S21 (Related to Figure 4): Superimposition of EphB2 monomer (1NUK, green) and tetramer (1KGY, magenta and orange) crystal structures highlighting changes in amino acid positioning of key residues that form the tetramer interface.

Video S22: Shows the EphB2-EphrinB2 circular tetramer highlighting the tandem positioning of juxtaposed tetramerization interfaces that snap two dimers into the circular tetramer. Coloring is same as above.
